# Lineage origin and developmental context shape tumor states in MYCN-dependent zebrafish neuroblastoma

**DOI:** 10.1101/2025.10.13.682025

**Authors:** Nora Fresmann, Julia Köppke, Anton Gauert, Anis Senoussi, Luca Bramé, Pedro Olivares-Chauvet, Aliki Grammatikaki, Marie Schott, Lennart Höfer, Marvin Jens, Dieter Beule, Anton G. Henssen, Nikolaus Rajewsky, Bastiaan Spanjaard, Anja I. H. Hagemann, Jan Philipp Junker

## Abstract

Neuroblastoma, a neural-crest-derived malignancy of the peripheral nervous system, is a devastating pediatric disease characterized by high intra- and intertumoral heterogeneity. While several tumor expression programs correlate with patient outcome, the developmental and contextual determinants of their emergence remain incompletely understood. Here, we systematically dissected neuroblastoma transcriptional heterogeneity and measured how tumor expression programs are associated with lineage history and developmental context. To achieve this, we combined single-cell transcriptomics with high-throughput lineage tracing and tumor cell transplantation in zebrafish models of MYCN-driven neuroblastoma. We identified developmental cell-state-associated programs that align with sympathoadrenal development, as well as general cellular-process programs. Clonal analysis showed that developmental programs are preferentially clone-associated, whereas cellular-process programs are more context-responsive. Transplantation into an embryonic environment reshaped the activation of several initially clone-associated programs, consistent with developmentally determined state reconfiguration and/or selection of state-competent cells. Together, our results show that lineage history and developmental context jointly shape tumor-state architecture *in vivo*.

## Introduction

Transcriptional heterogeneity and phenotypic plasticity are increasingly recognized as drivers of tumorigenesis, metastatic dissemination and treatment evasion^1,2^. Phenotypic differences between tumor cells can be described by their transcriptional states, which are defined by expression of gene modules – groups of co-regulated genes that comprise both cell identity-specific as well as physiological programs^3,4^. Plasticity, the ability of a cell to switch between different transcriptional states, is crucial during for example development, but is largely lost in fully differentiated cells in healthy tissues. Cancer cells override these rules, exhibiting the capability to switch between different phenotypes^5^. A key question is how the gene expression programs they access relate to their cell of origin, lineage history and exposure to developmental or microenvironmental cues.

Recent pan-cancer studies suggest that tumor cells access a common set of transcriptional programs related to general cellular processes, such as stress response or cell cycle^3,4^. In contrast to this, cell identity programs are cancer type-specific and derive from the cell type of origin and developmentally related cell types^6^. Efforts to link tumor cell lineage and state in cancer animal models have elucidated that a single cell can give rise to a complex tumor with diverse cell identities, e.g. alveolar type I, type II and gastric-like states in lung adenocarcinomas^7–9^. This state diversity arises from the interplay of cell-intrinsic mechanisms, including genetic and epigenetic state and interaction with the tumor microenvironment.

The link between differentiation and tumor cell state is particularly relevant for pediatric cancers, which arise from developmental precursor cells and where cell of origin and early developmental environment can profoundly influence tumor behavior^10^. Neuroblastoma (NB) is a childhood cancer with heterogeneous disease progression, high metastatic potential and low survival rates for high-risk patients^11,12^. NB arises from cells of the developing sympathoadrenal lineage with the earliest tumorigenic events occurring in the first trimester of pregnancy^13^. Amplification of the MYCN oncogene is observed in 20 % of NB patients and is a strong predictor for poor prognosis^14,15^. MYCN-amplification is an early event in NB-formation and studies have shown that MYCN alone can induce NB in neural crest derivatives^13,16–18^. These features make MYCN-driven neuroblastoma a useful context to study how developmental lineage history and oncogenic transformation jointly shape tumor-cell states.

In cell culture, some NB models can occupy noradrenergic/adrenergic (ADRN) and mesenchymal-like (MES) states, and interconversion between these states has been demonstrated for a subset of models^19–21^. MYCN, together with the transcriptional co-activator LMO1, has been shown to reinforce the adrenergic core regulatory circuit of transcription factors and to thus keep NB cells in an undifferentiated neuronal progenitor state^21,22^. Extensive single-cell studies of human neuroblastoma and fetal adrenal development have mapped neuroblastoma cells onto normal sympathoadrenal developmental trajectories, with most malignant cells resembling late neuroblast / sympathoblast states and subsets engaging bridge-, Schwann-cell-precursor (SCP)-, chromaffin- or mesenchymal-associated programs^23–28^. Complementary human stem-cell models have further shown that NB-associated lesions, including ALK mutation, recurrent chromosomal copy-number alterations and MYCN activation, can perturb neural crest and sympathoadrenal differentiation and generate NB-like states^29,30^. These studies provide an essential developmental framework for interpreting NB cell states. However, because developmental origin has largely been inferred from transcriptomic similarity and perturbation of developmental model systems, it remains unresolved how transcriptional state relates to clonal lineage history within established tumors *in vivo*. In particular, we cannot distinguish whether developmental-state-associated programs represent persistent lineage memory from distinct transformed founders, or whether similar programs can be acquired convergently by unrelated tumor clones in response to developmental or microenvironmental context.

Here, we address these questions using well-established *dbh*-driven zebrafish models of MYCN-dependent NB^31,32^, which allow controlled induction of tumors within noradrenergic / sympathoadrenal-lineage populations and joint measurement of transcriptional state and clonal history via high-throughput lineage tracing. In these transgenic lines, human *MYCN* is specifically activated in sympathoadrenal cells, leading to growth of tumors that histopathologically resemble human NBs. We found that tumors in this model are composed of clones from multiple cells of origin, allowing us to directly measure the influence of lineage on gene expression within individual tumors. We identified a range of NB transcriptional states that are either related to general cellular processes or to developmental cell-state-associated-programs. We found that development-associated states tend to be determined by the cell of origin and are subsequently stably expressed within cells of one clone, suggesting an important role for the developmental state of the cell of origin for tumor state. By transplanting primary zebrafish NB cells into zebrafish embryos, we showed that even initially clone-associated transcriptional programs can be reshaped after exposure to a developmental environment. This highlights the role of developmental signals received during early tumorigenesis for tumor cell state and potentially disease severity.

## Results

### Dissecting tumor expression heterogeneity by multiplexed single-cell transcriptomics

We used single-cell transcriptomics to analyze the diversity of tumor transcriptional states in the two established transgenic zebrafish lines that closely reiterate the pathogenesis of human neuroblastoma: Tg(*dbh:MYCN, dbh:EGFP*)^31^ and the double transgenic combination of Tg(*dbh:MYCN, dbh:EGFP*)^29^ with Tg(*dbh:LMO1, dbh:mCherry*)^32^, hereafter called MYCN and MYCN;LMO1, respectively. Because *MYCN* expression in these models is driven by the *dbh* promoter, tumor initiation is restricted to *dbh*-expressing sympathoadrenal-lineage cells. This provides a controlled setting with a limited cell of origin population to measure how lineage history shapes tumor cell states. To enable later assessment of the influence of cell lineage on transcriptional state, we combined scRNA-seq with high-throughput lineage tracing using CRISPR/Cas9 induced lineage barcodes^33–35^ (Fig. 1A, details described later in Fig. 3). Single-cell expression profiles and lineage barcodes were read out jointly by scRNA-seq in tumors from adult fish, using the cell hashing method MULTI-seq^36^, processing up to 14 tumors or tumor sub-samples in one scRNA-seq run to minimize experimental batch effects (Fig. 1A, Tables S1-2).

**Fig 1:**
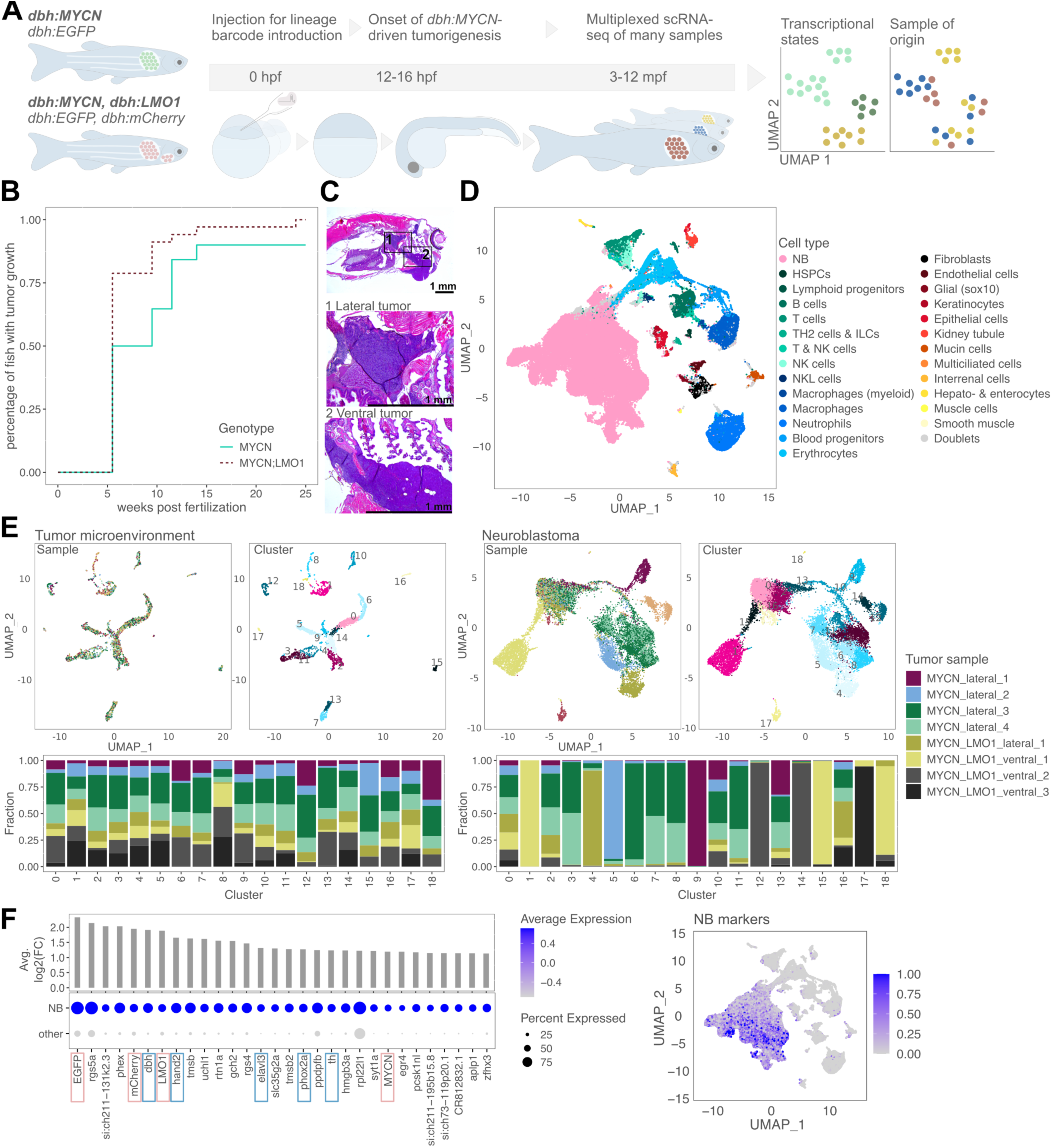
Multiplexed scRNA-seq to understand NB intra- and inter-tumor transcriptional heterogeneity. A) Overview of the main experiment: The two transgenic zebrafish lines MYCN and MYCN;LMO1 grow NB tumors in the interrenal gland (lateral site) and the latter also in the arch-associated complex (ventral site). We inject Cas9 and sgRNAs to later analyze cellular lineage relationships. We performed multiplexed scRNA-seq with MULTI-seq to gather single cell transcriptome data from several tumors at the same time with reduced batch effects. hpf = hours post fertilization, mpf = months post fertilization. B) Tumor incidence curves (monitoring between six and 25 wpf) for the two transgenic zebrafish models. C) H&E-stained sagittal section of a 3 mpf MYCN;LMO1 fish with magnified views of the lateral tumor in the middle and ventral tumor in the bottom image. D) UMAP of the entire dataset (around 150 thousand cells) colored by cell types: NB cells in pink, blood cells in green and blue hues, other stromal cells in brown to yellow hues. Cells with ambiguous marker gene expression are termed doublets and are shown in grey. E) 3451 TME cells (left) and 16482 NB cells (right) from one individual MULTI-seq run (multi_seq_09) sub-clustered and colored by tumor sample of origin or Louvain cluster in the UMAP. Barplots show the fraction of cells in each cluster made up of cells from a given sample. F) Top positively differentially expressed genes in NB cells when compared to all other cell types. Known noradrenergic / sympathoadrenal genes and transgenes are highlighted by blue and pink boxes, respectively. UMAP of all cell types shows expression score for the 46 most differentially expressed genes (signature ‘NB_markers’).

In agreement with previous reports, we found induction of tumors between 6 and 15 weeks post fertilization (wpf), with faster induction in the MYCN;LMO1 line (Fig. 1B). As expected, the fish developed tumors in the interrenal gland (IRG), which is equivalent to the human adrenal gland, and the superior cervical ganglion; we refer to these tumor locations as lateral. Additionally, only MYCN;LMO1 fish developed tumors in the arch-associated complex (AAC)^17,31,37^, a location we refer to as ventral (Fig. S1A). The AAC is a catecholaminergic, neural-crest-derived progenitor population adjacent to the pharyngeal arch vasculature that is described in zebrafish development as *th+/dbh+* ^17,37,38^. Thus, *LMO1* expression does not only increase MYCN-driven tumor penetrance, but also enables tumorigenic transformation in an additional cell population that *MYCN* expression alone cannot transform. Tumors in both locations showed the typical small round blue cell phenotype of NB^39^ (Fig. 1C). In total we sequenced the transcriptomes of 141,812 single cells from 60 tumors and 9405 cells from three healthy control samples (head kidneys with IRG) (Fig. 1D, Fig. S1B-D). We detected 101,872 NB cells, expressing sympathoadrenal and known NB markers *phox2bb*, *hand2*, *dbh* and tumor transgenes (Fig. S1C-D, Table S3). Robust expression of the *MYCN*-transgene is maintained in tumor cells over time as detected by scRNA-seq (Fig. 1F) and on the protein level (Fig. S1E)^40^. In the stromal and immune compartment of tumor samples, we detected 39,940 cells, including various kidney cell types (e.g. multiciliated cells) and steroidogenic interrenal cells from the IRG (Fig. S1B-C)^41^. Immune cells were likely derived from tumor immune invasion, but also partially from the hematopoietic tissue in the zebrafish kidney marrow, equivalent to human bone marrow (Fig. S1B).

Within individual MULTI-seq runs, technical batch effects were minimal with tumor microenvironmental (TME) cells from different tumor samples intermixing in clusters and on the UMAP (Fig. 1E, Fig. S1D). By contrast, tumor cells exhibited substantial inter- and intra-tumor transcriptional heterogeneity, reflected by clusters composed of cells from one or only few samples as well as cells of one tumor spread across multiple clusters. This is reminiscent of the pronounced phenotypic heterogeneity observed between patient samples.

### Zebrafish NB cells resemble sympathoadrenal developmental cell types

To characterize the overall transcriptional profile of NB cells, we performed differential expression analysis against all other cell types. NB cells expressed the tumor transgenes and noradrenergic / sympathoadrenal genes (including *dbh*, *hand2* and *elavl3/4*), whereas *pnmt*, the enzyme responsible for converting noradrenaline into adrenaline in differentiated chromaffin cells, was not detected (Fig. 1F, Table S4). The 100 most highly expressed genes in NB cells comprised almost exclusively ribosomal genes (signature *‘ribosomal_genes’*, Table S4), consistent with reported MYCN-driven increases in ribosome biogenesis^42,43^. We further calculated expression of the human cell line-derived ADRN/adrenergic and MES/mesenchymal NB signatures (van Groningen et al. 2017^19^). In line with findings in human primary tumors, the adrenergic signature was expressed in zebrafish NB cells, whereas the corresponding mesenchymal signature was more highly expressed in TME cells (Fig. S1F)^44^. Taken together, these findings show that zebrafish NB transcriptomes are overall noradrenergic and shaped by MYCN-activity.

As human neuroblastoma cells have been shown to transcriptomically resemble fetal sympathoadrenal developmental populations, including late neuroblast / sympathoblast, SCP-, bridge- and chromaffin-associated states^23–28^, we next sought to identify corresponding populations in zebrafish developmental scRNA-seq atlases. To compare zebrafish NB gene expression states with normal developmental programs, we extracted putative sympathoadrenal populations from the Daniocell and Zebrahub atlases (Fig. S2, Methods)^45,46^. In both datasets, these cells form transcriptomic continuum trajectories, spanning from SCP-like cells (*sox10*+) and a related cycling population, through a bridge state (*ascl1*+, *sox11a/b*+), to putative sympathoblasts (*elavl3/4*+, partially *dbh*+ and *th*+) and putative chromaffin-precursors (strongly *dbh*+ and *th*+, *elavl3/4*-low, *pnmt*-) (Fig. S2). This continuum functionally mirrors the established ontogeny of the zebrafish sympathoadrenal system, wherein multipotent SCPs give rise to both chromaffin cells and sympathetic neurons^47^. Furthermore, computational trajectory inference confirmed a directional developmental flow from these initial SCP-like progenitors toward both sympathoblast and chromaffin-precursor terminal fates, reinforcing this biological model (Fig. S3A-E)^48,49^. Projecting our zebrafish NB cells onto this developmental reference revealed assignments across all annotated cell types (Fig. S3F). However, the largest proportion of zebrafish NB cells mapped to putative chromaffin-precursors. This contrasts with human NB, which predominantly resembles neuroblasts / sympathoblasts^23^. Assigning zebrafish NB cells to their closest human developmental cell type resulted mainly in assignments to neuroblasts as well, suggesting that the observed discrepancy is likely caused by the technical limits of distinguishing between late-stage sympathoblasts and chromaffin cells using current zebrafish developmental atlases (Fig. S1G). Overall, this suggests that both zebrafish and human NB cells mainly resemble noradrenergic developmental populations.

Beyond the overarching NB transcriptomic profile, we next sought to dissect the observed heterogeneity within this population by performing a systematic *de novo* analysis of tumor gene expression programs.

### The spectrum of MYCN-driven NB transcriptional programs

Gene expression in tumor cells has been shown to be composed of multiple gene expression programs, which can be active to different degrees in individual cells^3,4^. Clustering alone may hence merge multiple sources of transcriptional variation into composite cell states. We therefore used non-negative matrix factorization (NMF^50^) to decompose zebrafish NB transcriptomes into gene expression modules, providing a modular representation of the tumor transcriptome.

In order to identify gene modules that capture both intra- and inter-tumor expression variation, we performed NMF on three different levels (Fig. 2A). In the first instance, we ran NMF on NB cells from individual tumors separately, resulting in a list of modules representing expression variation between NB cells within each tumor. We grouped these modules by their similarity in gene content to derive recurrently activated consensus modules (Fig. S4A, Methods^3,4,42^). We repeated this analysis using NB cells from individual MULTI-seq runs as input to identify gene modules that are differentially activated between tumors without suffering from batch effects in the data. Lastly, we also ran NMF on the whole NB cell population from all samples to identify gene modules that are only activated in few samples or cells. We annotated modules by associated Gene Ontology (GO)-terms and the functional annotation of individual genes contributing to them (Tables S4-7). Because NMF identifies networks of genes that strongly co-vary in their expression across single cells rather than grouping genes by strict functional ontology, these modules inherently capture broader transcriptional programs or comprehensive cell states, including core functional genes alongside genes involved in distinct cellular processes. We emphasize that descriptive module labels (e.g., “*catecholamine_production*”) serve as shorthand names based on the top-weighted driver genes and GO-terms. All three approaches showed some overlap in detected gene modules (Fig. S4B). We therefore compiled a final combined list of modules that are largely non-overlapping in terms of their gene content and which represent the spectrum of NB cell states (Fig. 2B, Table S6, Methods).

**Fig 2:**
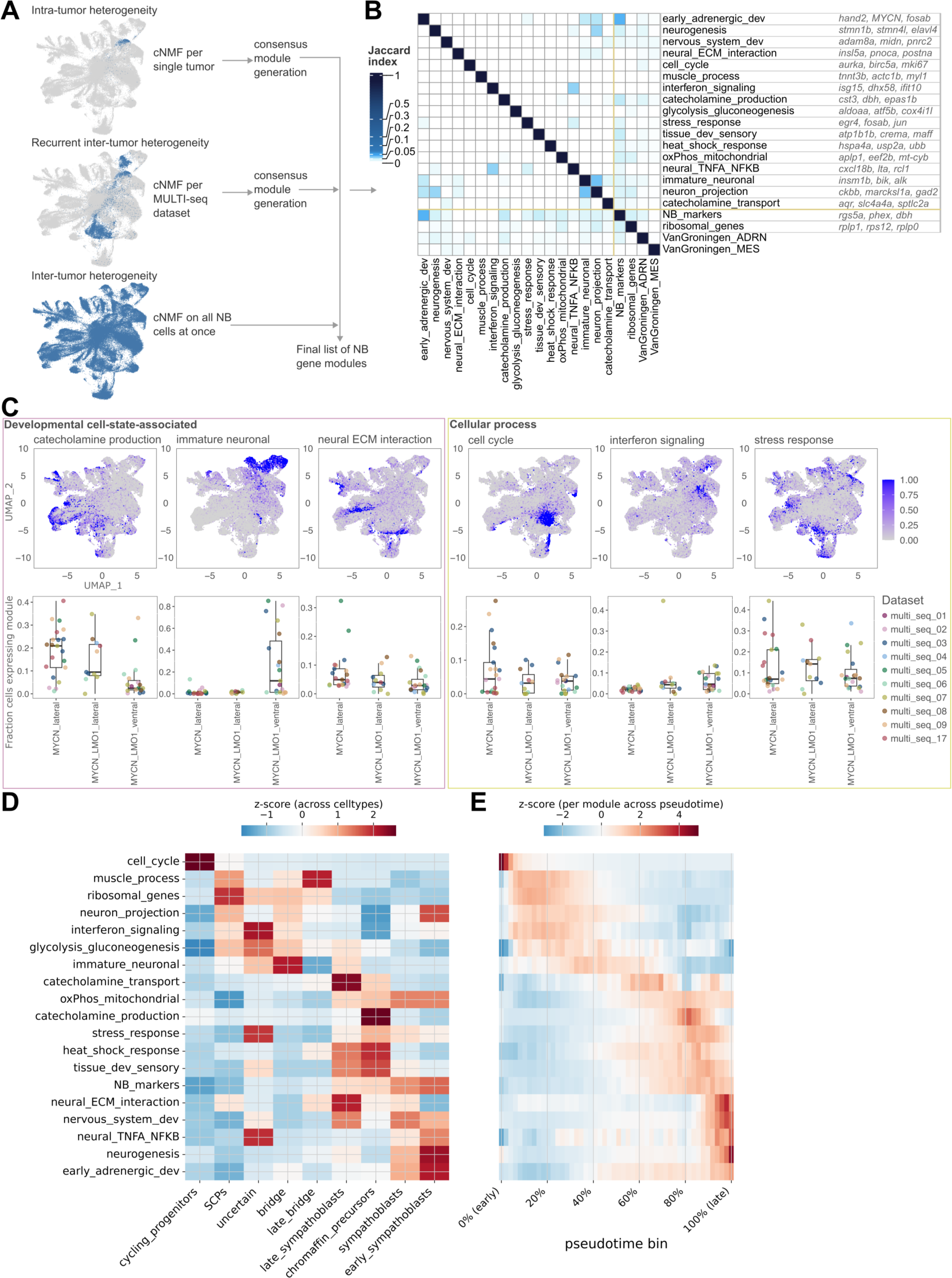
The spectrum of transcriptional states of MYCN-driven zebrafish NB. A) Gene expression module identification process: A three-fold approach to detect both modules describing intra-tumor heterogeneity (expressed in a subset of cells in most tumors) and inter-tumor heterogeneity (expressed only in some tumors and absent in others). Modules from all approaches were summarized to derive consensus-modules shown in B. B) Summary of final list of modules compiled from all three approaches in A with Jaccard index of gene content overlap for all modules as well as the NB differential and high expression signatures (‘*NB_markers*’ and ‘*ribosomal_genes*’ as in Fig. S1E) and the published human ADRN/adrenergic and MES/mesenchymal NB signatures (as in Fig. S1F). The top three genes contributing to each gene module are shown on the right. C) UMAPs of NB cells with expression scores for the indicated modules. Box plots and jitter show the fraction of cells that express a given module per tumor. Modules were grouped into developmental cell-state-associated (pink box) and general cellular processes (yellow box). D) Z-score of expression of each of the 19 gene modules in zebrafish developmental cell types, transferred from NB cells in a shared transcriptomic embedding. E) Z-score of module expression along pseudotime (modules as in D) determined by transfer of pseudotime from developmental cells to NB cells in a shared transcriptomic embedding.

The resulting 17 gene modules comprised both sympathoadrenal and neuronal identity signatures (termed “developmental cell-state-associated” programs, e.g. *catecholamine_production*, *immature_neuronal*) and general, non-cell-type-specific cellular processes (e.g. *cell_cycle*, *interferon_signaling*) (Fig. 2C, Fig. S4C, Table S5-6). To interpret the developmental modules, we benchmarked them against the annotated zebrafish sympathoadrenal reference^45^. Specifically, we assessed module expression across discrete developmental cell types (Fig. 2D) and along developmental pseudotime (Fig. 2E). In the zebrafish developmental reference, the *neurogenesis* module was highly enriched in sympathoblasts, whereas *catecholamine_production* mapped predominantly to chromaffin-precursor cells, and *neural_ECM_interaction* was associated with late sympathoblasts. The *immature_neuronal* module peaked earlier in pseudotime than most other developmental programs, showing highest expression in bridge cells. Notably, the *immature_neuronal* module includes *alk*, a developmental receptor tyrosine kinase with known relevance in NB biology^51–53^. To validate these findings in the context of human biology, we scored expression of the modules in human fetal adrenal cells^25^. Individual modules mapped to distinct human developmental states, partially mirroring the zebrafish trajectories: *neurogenesis* correlated with sympathoblastic cells, while *catecholamine_production* aligned with chromaffin-precursor cells (Fig. S5A). Together, these findings indicate that our identified gene modules capture transcriptomic variation along a sympathoadrenal differentiation axis.

### Zebrafish NB gene modules capture human NB transcriptional variation

We found that all detected modules also varied across individual cells and tumors in our zebrafish NB dataset, validating that they at least partially capture the observed tumor expression heterogeneity (Fig. 2C, Fig. S4C). Some modules also showed a clear dependence on tumor location: *immature_neuronal* was preferentially activated in ventral tumors, whereas *catecholamine_production* was more highly activated in lateral tumors, consistent with a relative shift of lateral tumors toward stronger catecholaminergic differentiation. Multiple modules contain higher levels of the *MYCN* transgene (*early_adrenergic_development*, *neuron_projection*, *immature_neuronal*), supporting the notion that MYCN drives several distinct zebrafish NB cell transcriptional states. Expression of the *early*_*adrenergic_development* and *immature*_*neuronal* modules was particularly associated with *MYCN* expression levels and the expression of published MYCN-driven genes (Fig. S6A)^54^, while *catecholamine_production* was more weakly associated with *MYCN* expression. To further test the relevance of these modules for human cancer, we compared them to gene modules from recent cancer studies and scored their expression in published human NB datasets (Tables S8, S9). As expected, programs like *stress_response* and *interferon_signaling* resembled general programs activated in many cancers^3,4^ (Fig. S6B). Conversely, multiple developmental cell-state-associated zebrafish modules overlapped with broad adrenergic / neuronal programs derived from human NB^19,27,44,55^ (Fig. S6C). Expression scoring of the zebrafish NB modules in published scRNA-seq and bulk RNA seq data of human NB showed that some activity of all modules is detectable and that expected risk associations, such as higher expression of the *cell_cycle* module in high-risk disease are recapitulated (Fig. S6D-G, Tables S9-10)^11,24,25,56–61^.

Together, these findings demonstrate that the zebrafish-derived modules capture key oncogenic and differentiation-associated programs relevant to human NB biology. We hypothesized that activation of some modules might be clonally determined, for instance by the cell of origin that a tumor cell was derived from, while others may be regulated in a niche-dependent manner. We further speculated that the degree of lineage determination might be higher for modules related to developmental cell states compared to modules related to general cellular processes. In order to test this, we next analyzed the clonal structure of zebrafish NB tumors using lineage tracing.

### High-throughput lineage tracing identifies multiple cells of origin per tumor

To experimentally measure the influence of lineage on transcriptional state, we combined scRNA-seq with high-throughput lineage tracing using CRISPR/Cas9 induced lineage barcodes^33–35^, which are created by injection of Cas9 and sgRNAs targeting lineage recording sites into zebrafish embryos at the one-cell stage (Fig. 3A, Methods). In our system, lineage barcodes are created on multiple integrations of a cassette of three Cas9 target sites in the 3′-untranslated region (UTR) of a dsRed transgene (Fig. S7A-C) as well as in the 3′**-**UTRs of seven endogenous genes (Fig. 3A). Lineage barcodes predominantly consist of small insertions or deletions around the Cas9 target site (Fig. 3B). We determined that these barcodes are generated within the first 8 hours post fertilization (hpf) (Fig. 3C), preceding the onset of *dbh* expression at 14 hpf^45^ (Fig. 3D). Consequently, activation of the tumor-inducing *dbh:MYCN* transgene occurs after lineage barcode creation, ensuring that all progeny of a given transformed cell inherit the same lineage barcodes^17^ (Fig. 3A). By contrast, tumor cells derived from different cells of origin can be distinguished based on differing lineage barcodes.

**Fig 3:**
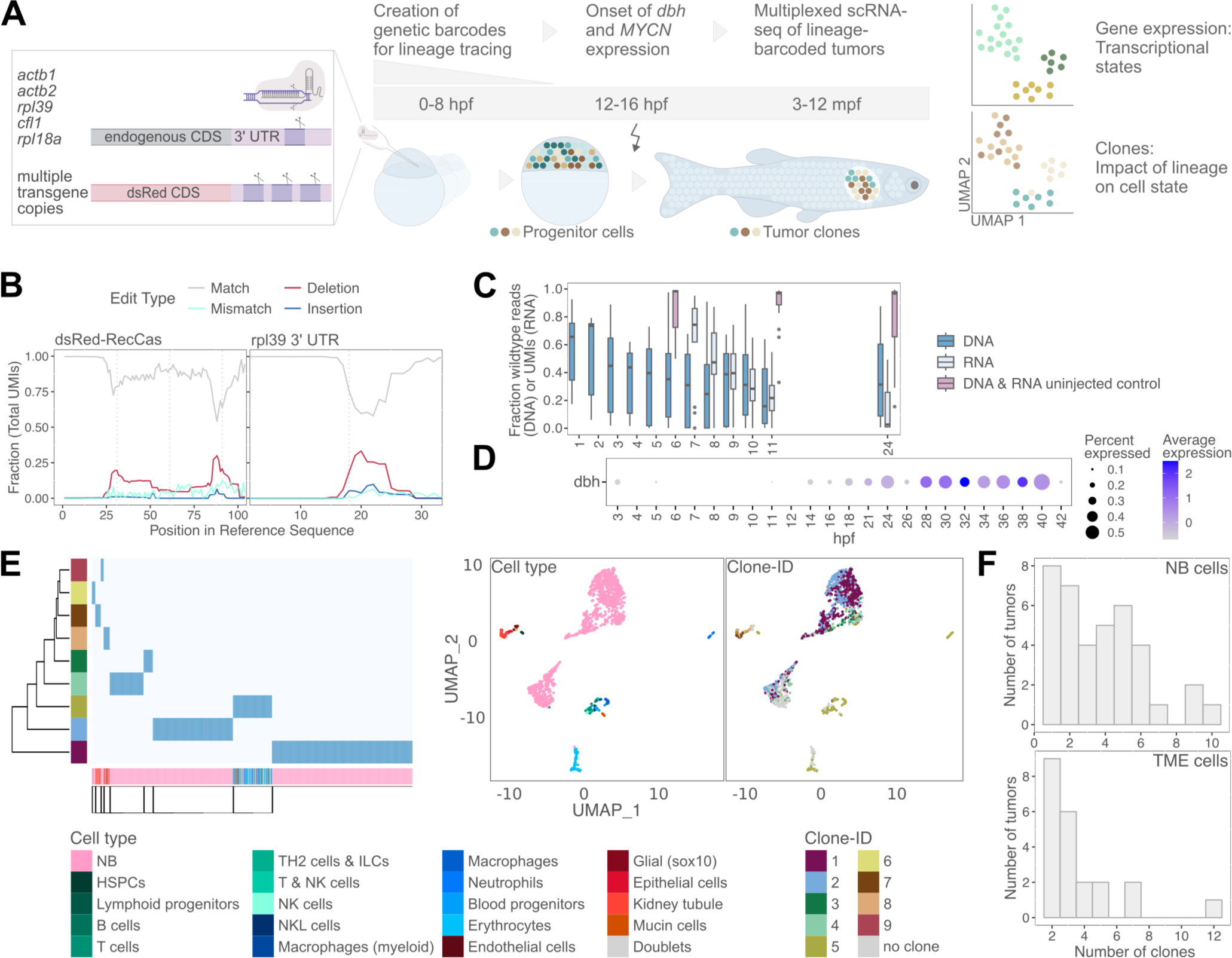
Tracing tumor cell clones derived from distinct cells of origin. A) Experimental strategy for early developmental lineage tracing via CRISPR/Cas9. Cas9 and sgRNAs targeting a transgene carrying three target sites in the 3’ UTR as well as single target sites in the 3’ UTR of the listed highly expressed endogenous genes are injected at the one-cell stage. The repair of Cas9-induced DNA-lesions results in the formation of indels used as lineage barcodes. Lineage barcodes are read out together with the cell transcriptomes, when tumors are processed via scRNA-seq. B) Base position plots for one NB MULTI-seq dataset showing the fraction of UMIs with edits in a given base in the transgenic recording cassette locus (dsRed-RecCas, left) and the endogenous rpl39 3’ UTR locus (15 bases upstream and downstream of the expected Cas9 cut-site). Border between target (spacer) and PAM-sequence for each target is marked by a vertical grey line. C) Lineage barcode creation dynamics show a rapid loss of uncut alleles and thus introduction of lineage barcodes in the first hours of development after CRISPR/Cas9 injection. Lineage barcode introduction on DNA saturates around 5 hpf and unedited RNA is largely replaced at 11 hpf. Samples from at least two separate injection experiments were taken for each time point and assay. D) Expression dynamics of *dbh* in scRNA-seq of zebrafish development (Daniocell atlas^45^). *dbh* expression is only observed in few individual cells prior to broad activation at around 14 hpf. E) Hierarchical clustering of 1099 cells from a single tumor (MYCN_lat_m9_1) based on assigned clone-IDs (dark blue heatmap color: assigned to clone). Cells are annotated by cell type at the bottom of the heatmap. We found clear lineage splits between one immune/blood cell clone and two stromal cell clones and identified six distinct NB cell clones. The UMAPs show cells from the same tumor including those that could not be assigned to a clone (total of 1410 cells) colored by cell type or clone-ID. F) Number of clones detected in NB cells (top) or the TME cell compartment (bottom) per individual tumor (n = 38).

Joint analysis of single-cell transcriptomes and lineage barcodes revealed that NB cells and TME cells typically have separate lineage barcode profiles, in line with the distinct lineage origins of these cell types (Fig. 3E). Furthermore, we found that the NB cells of a single tumor were composed of cells with multiple distinct lineage barcode profiles and were hence derived from multiple cells of origin (Fig. 3E). We attribute this to the strong effect of *MYCN* in our genetic models, which induces tumorigenic transformation in many cells. We hereafter refer to these groups of cells from different origins as clones. We clustered NB cells from all 38 tumors with lineage information into clones according to their lineage barcode pattern across all lineage reporter sites, focusing on maximizing clonal resolution (Methods, Fig. S7D-G). We typically found between 2 and 6 NB cell clones (and hence cells of origin) per tumor (Fig. 3F, S7H-L). We found that ventral and lateral tumors originating from the same fish always had completely distinct clonal composition, indicating different lineage origins between these two tumor types (Fig. S8A-B). This is in line with the AAC-region tumors being independent primary lesions from a distinct developmental population, as opposed to these tumors originating from dissemination from lateral tumors. The multi-cellular origin of individual tumors now allowed us to study gene expression differences between NB cell clones in a shared environment, and thus assess lineage effects on transcriptional state independently of confounding effects related to differences between individual tumors.

### Clonal analysis separates clone-associated developmental programs from context-responsive programs

We next sought to use this approach for quantifying to which degree transcriptional heterogeneity within one tumor is driven by different cells of origin. In an extreme scenario, tumor states are fixed by the cell of origin, with no expression state transitions, leading to co-segregation of cells by gene expression and clone (Fig. 4A). In the opposite scenario, gene expression is highly context dependent (e.g. spatial position, cell-cell interactions), and tumor states are independent of the cell of origin. Intermediate to this, distinct cell states of origin may interact in specific ways with contextual signals.

**Fig 4:**
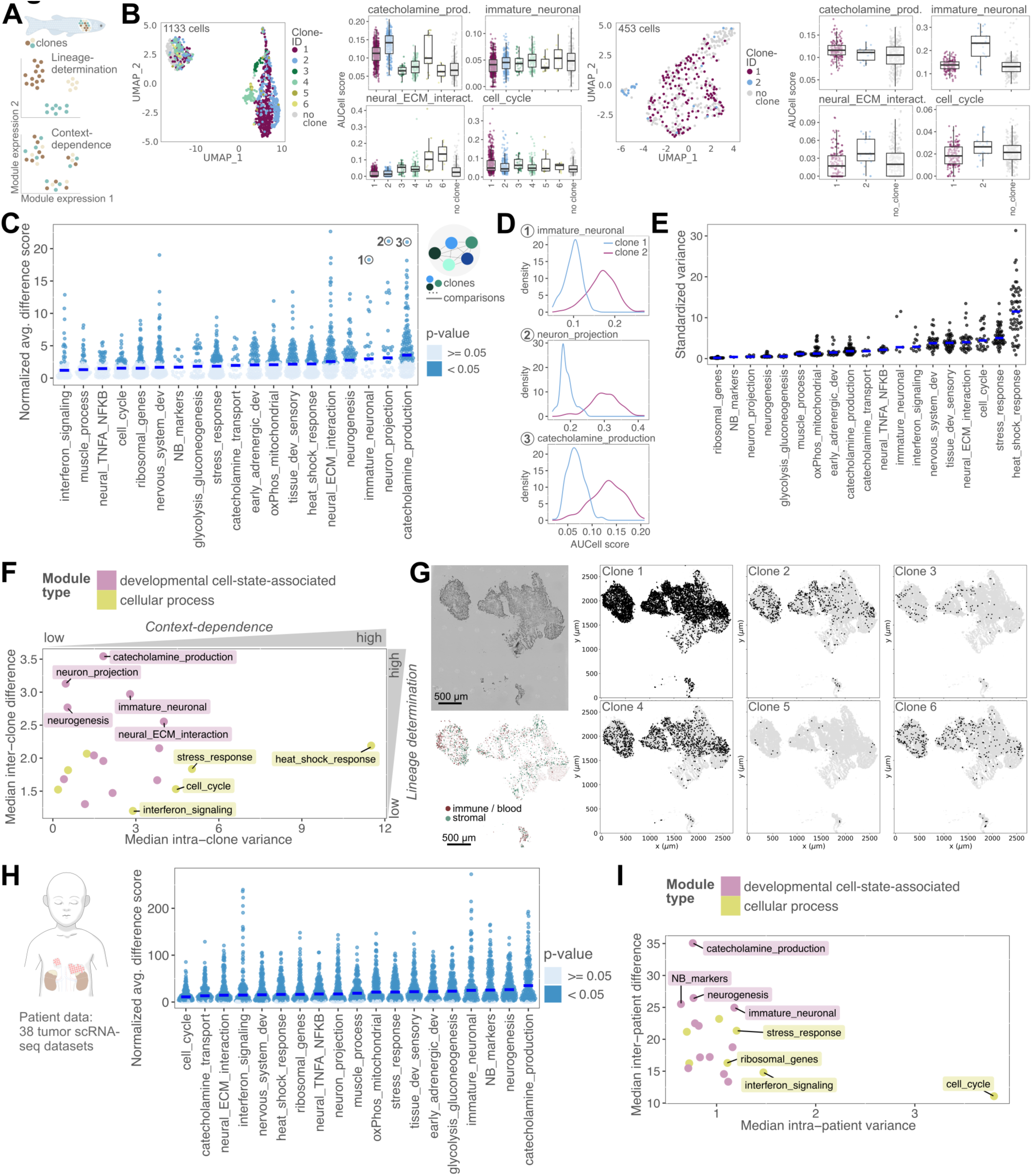
Clonal analysis of NB reveals clone-associated transcriptional programs. A) Opposing scenarios of tumor state regulation: In the first scenario, module expression is determined by the cell of origin (i.e. clone, indicated by color). In contrast, if tumor states depend on the current conditions in the tumor niche, cells dynamically take on different tumor states, regardless of their origin. B) UMAP showing NB cells from one tumor colored by the assigned clone for a lateral tumor (left) and a ventral tumor (right). The AUCell-determined expression scores for four modules in NB cells grouped by clone is shown right to the UMAP. C) Differential module expression scores for comparisons between pairs of clones coming from the same tumor (= one dot) with the median differential expression score indicated by a blue line. Only pairs with detectable module expression in at least one clone were included (Methods), ranging from 18 pairs for module ‘*NB_markers’* to 188 pairs for module ‘*ribosomal_genes*’. The distributions of module expression scores for three example comparisons are shown in D. D) Distribution of AUCell-determined module scores for the pairs of clones compared in the highlighted examples in C. E) Standardized intra-clone variance of gene module expression with the median variance indicated by a blue line. Only clones with detectable module expression were included (Methods), ranging from 2 clones for module ‘*NB_markers*’ to 63 clones for module ‘*ribosomal_genes*’. F) Summary plot showing median inter-clone difference (as in C) versus median intra-clone variance (as in E) for each module, highlighting developmental cell-state- and cellular process associated modules. G) Clonal distribution in space using spatial transcriptomics data. Left: Microscopic image of the tumor section and indication of spots assigned to immune or blood cell and stromal cell types. Other spots are mainly NB cells. Right: Spatial outline of the tumor section highlighting spots, in which lineage barcode sequences representative of the indicated clones (derived from scRNA-seq data) were found. H) Differential module expression scores for comparisons between pairs of tumors from different patients from a compendium of 38 human NB scRNA-seq datasets with the median differential expression score indicated by a blue line. I) Summary plot showing median inter-patient expression difference (as in H) versus median intra-patient expression variance for each module.

To assess the clonal determination of specific states, we first examined expression of the modules identified in Fig. 2 in the different clones of selected individual tumors. This modular representation enabled us to move beyond asking whether entire clusters are clone-associated, and instead quantify clone-association separately for each transcriptional program. We found that some modules, such as *cell_cycle*, were expressed at similar levels across the different clones of a tumor, while developmental cell-state-associated modules (e.g. *catecholamine_production*) tended to vary between the clones of a tumor (Fig. 4B, Fig. S8A). For a systematic analysis of tumor state association with clonal origin, we next calculated the differential module expression between all clone pairs within the same (sub-)tumor (Methods, Fig. 4C-D, S8C). The inter-clone differences reported by this analysis correspond to the effective state determination by the clonal origin. We observed considerable differences in clone-association between the gene modules, with a subset of developmental cell-state-associated programs such as *catecholamine_production, immature_neuronal* and *neurogenesis* being particularly clone-associated (63 %, 56 % and 53% significant comparisons respectively), whereas several modules related to general cellular activity (e.g*. interferon_signaling, cell_cycle* and *stress_response*) were more context-responsive (25 %, 28 % and 38 % significant comparisons respectively). Conversely, when we calculated differential module expression between groups of cells from the same clone residing in different sublocations of the same tumor, we found that the modules with most frequent expression differences were cellular process modules, further indicating their activation is more context-dependent, for example on the location of a cell in the tumor (Fig. S8C).

The inter-clone expression differences we observed suggest that developmental cell-state-associated modules tend to be determined by their clonal origin and will likely be activated at a similar level across cells within a clone. To test this hypothesis, we next determined the intra-clone expression variance of the gene modules. Indeed, we found a tendency for developmental cell-state-associated modules to exhibit lower intra-clone variance than modules associated with general cellular processes (Fig. 4E, Methods). Together, these metrics suggest stable clone-associated deployment of developmental programs and stronger local context dependence of general cellular process programs (Fig. 4F, Table S11).

### Inter-clone expression differences occur in the absence of genetic or niche differences

The observed differences between cancer cells from different cells of origin might originate from different genomic alterations in these populations. However, we detected neither additional mutations in whole exome sequencing (WES) on two new tumor samples nor copy number variants when inferring copy numbers from scRNA-seq data (Fig. S9). We additionally performed whole genome sequencing (WGS), taking into account that zebrafish tumors are multi-clonal and that clonal CNVs may be more difficult to detect. We found that in MYCN lateral tumors, eight out of twelve tumors contain at least one clone with a relative size of 25 % or more with differential expression of a gene module (Fig. S10A). We performed WGS on five MYCN zebrafish and found that across all five tumor/normal WGS pairs, no genomic segment ≥ 1 Mb showed a log2 copy number ratio outside the range (−0.15, 0.1) (Fig. S10B). All segments remain within the range expected for a clone fraction below 25 %. In the five tumors assessed, these data indicate the absence of somatic copy number changes of sufficient amplitude and extent to account for the clone-specific module expression observed by scRNA-seq. Given this data, the chance that copy number changes can explain the existence of the observed clones with differential expression of such programs is p = 0.02 (Fisher’s test, one-sided).

To further exclude the possibility that an uneven distribution of clones within a tumor would confound the observed inter-clonal expression differences, we performed Open-ST^62^-based spatial transcriptomics to assess the localization of clones in a proof of concept experiment. The spatial data was analyzed at the level of 5 µm diameter bins. All major cell types identified in scRNA-seq data were also found in the spatial data and we found most of the previously detected modules activated in spots across the tumor mass (Fig. S11, Table S8). To identify clones in the spatial transcriptomics data, we extracted clone-specific lineage barcode sequences from scRNA-seq data of the same tumor and analyzed their distribution in space (Fig. 4G, Fig. S12). All six NB cell clones identified in the scRNA-seq data were found in the spatial data. Cells from all these clones were spread across a large area of the tumor section and were well intermixed. Co-occurrence of distinct lineage barcodes in spatial spots confirmed that cells from different clones often occupied the same neighborhoods (Fig. S12F). This example indicates that clones show expression differences even when they occupy the same region of a tumor, emphasizing the importance of the cell of origin for emergence of inter-clonal expression differences.

Spatial transcriptomics further allows the analysis of effects of intercellular interactions on tumor cell transcriptional states. Spatial analysis revealed that TME composition has an overall effect on NB transcriptional states with most NB expression modules correlating positively with TME cell presence, whereas the *ribosomal_genes* module was enriched in the tumor core and inversely associated with presence of most TME cell types (Fig. S13A-B, Methods). In the scRNA-seq data, we found several likely immunosuppressive interactions between NB cells and immune cells (Fig. S14A)^63^. Expression scores for multiple modules correlated positively with the expression of signaling factors in NB cells, suggesting that activation of these modules is associated with a more immunosuppressive phenotype (Fig. S14B). We also observed a correlation between the abundance of multiple macrophage populations as well as cytotoxic T-cells and the activation of the *interferon_signaling* module, indicating TME cell types can influence context-dependent modules (Fig. S14C-E). Overall, these analyses support an association between tumor niche composition and tumor-cell transcriptional state, both regarding tumor area (core vs. periphery) as well as presence of certain immune cell populations. However, in contrast to lineage origin, it does not explain the specific activation of most modules.

In summary, these analyses reveal a spectrum from clone-associated developmental programs to dynamically regulated cellular-process programs, rather than a binary division between fixed and plastic tumor states. Among the stably clonally activated modules, we identified the more differentiated *catecholamine_production* module, as well as the more primitive, *alk*-containing *immature_neuronal* module. This suggests that such distinct developmental states, which carry potential relevance for disease progression, are either inherited from the cell of origin or established very early in tumorigenesis.

### Lineage-associated gene modules are differentially activated between human tumors

In patient data, module expression determined by the cell of origin would manifest in differential expression between tumors, assumed to be of mono-cellular origin. We tested inter-tumor module expression differences and intra-tumor expression variance using published scRNA-seq data from 38 tumors (Fig. 4H-I, Table S12)^24,25,56–58^. Developmental cell-state-associated modules were still consistently the programs with highest inter-tumor expression differences. Despite less clear separation of module types by intra-patient variance, variance for many developmental cell-state-associated modules was low. This suggests that, despite a higher likelihood for complex tumor evolution and presence of genetically distinct tumor subclones in patient tumors, activation of programs associated with developmental cell types may also be relatively stable and potentially influenced by the cell of origin or early events in tumorigenesis in patients.

### Tumor cell transplantation into an embryonic environment

We next asked how stable clone-associated tumor programs remain after exposure to a developmental environment. To test this, we transplanted batches of around 150 lineage-barcoded primary tumor cells into wildtype zebrafish embryos at the blastula stage. Transplantation of cells from the same clones into multiple embryos enabled their subsequent recovery from different hosts at several timepoints. By processing and transplanting cells from multiple primary tumors at once, we ensured sufficient material for transplantation and increased the number of clones observed per experiment. We profiled cells with scRNA-seq directly after primary tumor dissociation and at two engraftment stages: Tumor cells were FACS-enriched and sequenced alongside larval host cells 2 days post transplantation (dpt; early allografts) and whole graft tumors were processed several months post transplantation (mpt; late allografts) to track clonal states over time (Fig. 5A). Zebrafish NB cells spread throughout the host larvae after transplantation and formed secondary tumors two to six months later (Fig. 5B, Fig. S15A-B), most commonly in the orbital cavity of the eye, near the AAC (heart-proximal) and the superior cervical ganglion – sites populated by neural crest derivatives or known to be NB primary or metastatic sites^64,65^ (Fig. 5C). Integrated analysis of scRNA-seq data of all primary tumors and graft samples revealed that tumor cells from the grafts, identified by the presence of lineage barcodes, cluster together with the primary tumor cells (Fig. 5D-E, Fig. S15C-E). Cell-cell communication analysis further showed that NB cells continue to express putative immunosuppressive signaling factors as observed in the primary tumor (Fig. S16A-B). Thus, NB cells retain a broad noradrenergic / neuroblastic identity.

**Fig 5:**
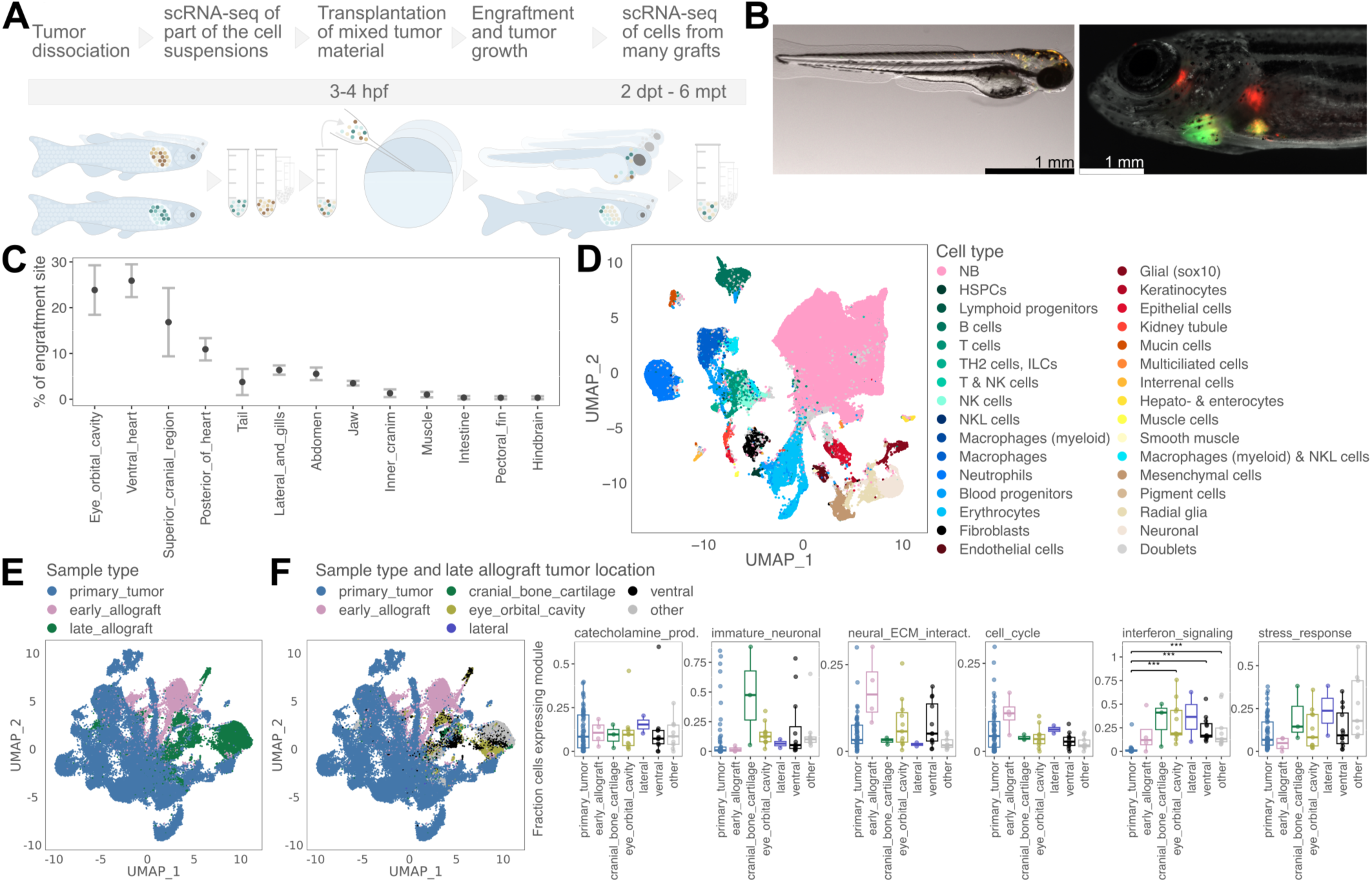
Tumor cell transplantation into an embryonic environment. A) Experimental workflow for transcriptional profiling of clones over multiple transplantation time points: Lineage-barcoded cells from multiple tumors are extracted and a part is sequenced, while the rest is pooled and transplanted into many wildtype zebrafish blastulae. Transplanted cells are extracted for sequencing two days or months after transplantation (d/mpt). B) Fish engrafted with a mix of MYCN (green) and MYCN;LMO1 (red and green) tumor cells three days after transplantation (left) or 2.5 months after transplantation (right). C) Mean percentage of tumors in different engraftment sites three months after transplantation in three separate transplantation experiments. Error bars denote the standard error of the mean. D) UMAP of all cells from primary tumors, 2 dpt allografts and late allograft tumors colored by cell type (around 208 thousand cells). Cell types derived from healthy tissues of the host larvae at 2 dpt are colored in beige tones (mesenchymal cells, pigment cells, radial glia, neuronal). E) UMAP of NB cells from all time points (around 131 thousand cells) colored by sample type (primary tumor, early allograft, late allograft). F) UMAP of NB cells from all time points colored by sample type (primary tumor, early allograft) or by tumor location for samples derived from late allografts. Fraction of cells per tumor expressing the indicated module with samples grouped as in the UMAP. Significance is only shown for significant comparisons as determined by pairwise Wilcoxon rank sum test (* p < 0.01, ** p < 0.001, *** p < 0.0001).

### Embryonic exposure reshapes clone-associated developmental programs

To compare gene expression modules before and after transplantation, we repeated the NMF-based gene module identification on both early and late graft tumor cells (Methods, Fig. S16C). We found that the vast majority of modules represented processes also identified in the primary tumors, indicating that they do not reflect novel transcriptional states induced by engraftment (Fig. S16D). Analysis of module expression across time points and locations revealed a generally high degree of transcriptional variation (Fig. 5F), both in cellular process- and in developmental cell-state-associated modules. We thus continued to use the list of modules identified in primary tumors for all further analyses

To determine how tumor programs respond to transplantation, we tracked individual clones over time using lineage barcodes. We assigned graft cells to primary tumor clones based on shared lineage barcode patterns, allowing us to track 9 clones over three and 18 clones over two time points (Methods, Fig. S17A-B). Overall, grafted NB cells transiently upregulated genes related to interferon signaling and the cell cycle, reflecting a short-term cellular response to the early developmental environment (Fig. 6A, Fig. S17C). Interestingly, we observed multiple cases in which clone-associated modules such as *catecholamine_production* or *immature_neuronal* were down- or upregulated upon transplantation, indicating that developmental exposure can reshape the activation of initially clone-associated developmental programs after transplantation. We then looked at an individual clone in more detail: Clone 6_8 in dataset #1 – derived from a lateral MYCN;LMO1 tumor – which had contributed to seeding of multiple graft tumors in different host fish and tumor locations. Depending on the specific graft tumor, cells from this clone exhibited varying expression levels of programs previously found to be stably expressed within a single primary tumor clone (e.g. *catecholamine_production*), further suggesting that even initially clone-associated programs can be reshaped after transplantation (Fig. 6B).

**Fig 6:**
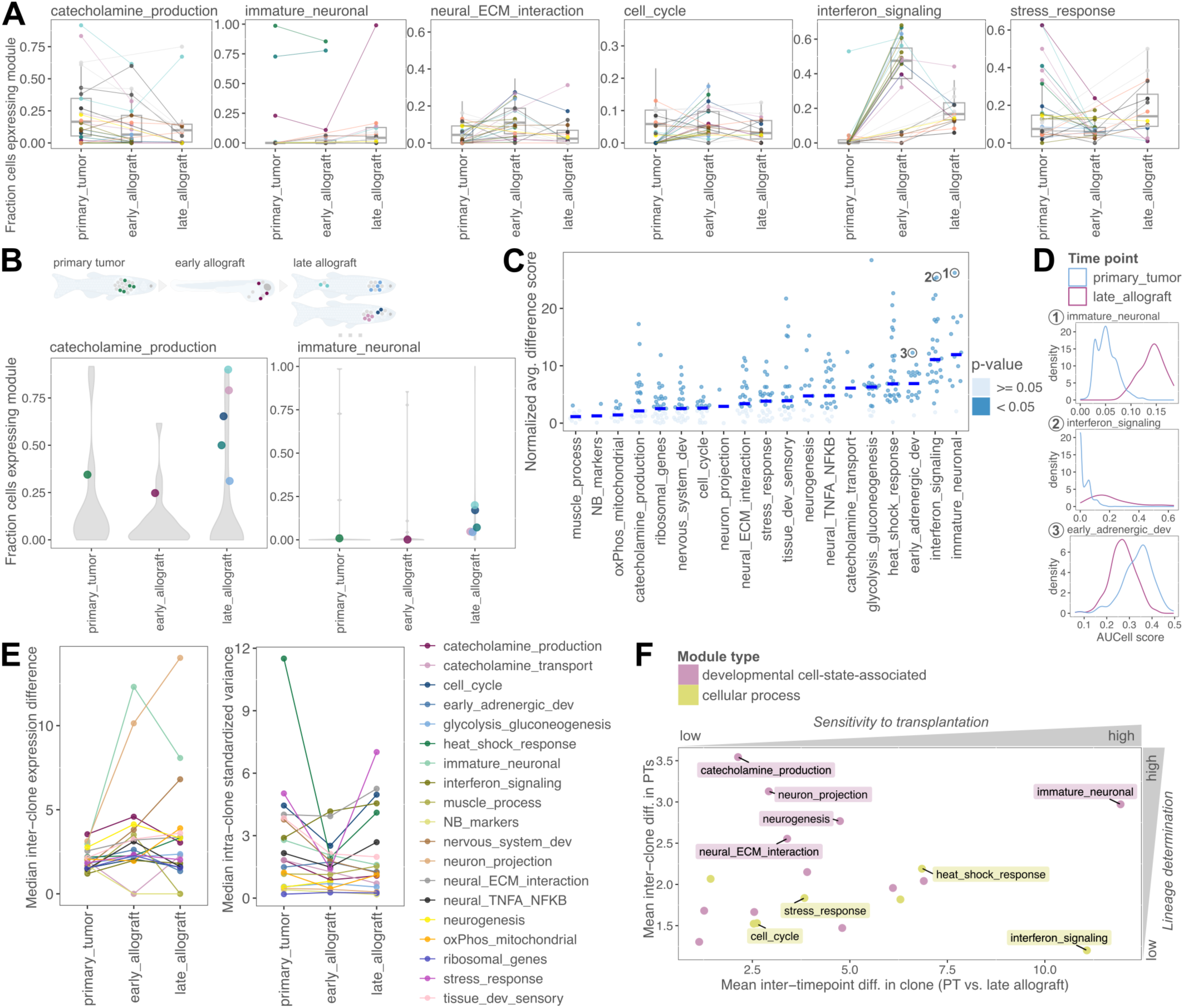
Embryonic exposure reshapes clone-associated developmental programs. A) Fraction of cells per clone and time point that express a given module. Each dot shows a clone in a specific time point and groups of cells from the same clone in different time points are connected via lines. B) Fraction of cells from a clone in a time point and location expressing the indicated modules. Each dot represents cells of the clone in distinct locations (as illustrated in the sketch above the plot: primary tumor, early allograft timepoint, multiple late allograft timepoint tumors). Grey violins show background distribution of module expression fraction for all clones in that dataset. C) Differential module expression scores for comparisons between cells from one clone found in two different timepoints (one dot = comparison between group of cells from one clone in primary tumor with group of cells of the same clone in late allograft tumor). Only pairs of cell groups with detectable module expression in at least one group were included (Methods), ranging from 2 pairs for module ‘*neuron_projection*’ to 19 pairs for module ‘*stress_response*’. Module expression score distributions for three examples are shown in D. D) AUCell-determined module expression distribution for the respective module in the cells from one clone in the primary tumor and the late allograft tumor is shown for three examples highlighted with circles in C. E) Median inter-clone difference (left) and median intra-clone variance (right) for each module across clones in the three sampled timepoints. Inter-clone difference was calculated for pairs of clones found in the same tumor or allograft sample. For both measures, only clones with detectable expression of a given module were included, as described in Fig. 4C and E. F) Summary plot showing median module expression difference between cells from one clone in two time points (primary tumor and late allograft, as in C) versus median inter-clone difference in the primary tumor (as in Fig. 4C). Highlighted modules are classified as developmental cell-state- or cellular process-associated.

To substantiate these findings, we next performed a systematic analysis of differential module expression between primary tumor and late allografts using all clones captured in these two timepoints. Unlike the analysis in Fig. 4C which was focused on pairwise comparison of clones within one time point, we now computed the expression differences within individual clones across time points (Fig. 6C-D, Fig. S17D). This longitudinal analysis represents the sensitivity of specific tumor programs to a new embryonic niche. The modules *interferon_signaling* and *immature_neuronal* exhibited the most profound expression differences between primary tumors and late allografts. Comparison with the intermediate “early allograft” time point revealed distinct kinetics: *interferon_signaling* is rapidly upregulated immediately after transplantation, whereas the *immature_neuronal* program is markedly altered only in late allograft tumors, suggesting activation later than two dpt (Fig. S17D). Consistent with this interpretation, moscot-based transition inference linked primary tumor cells with high *catecholamine_production* activity to late allograft cells with *immature_neuronal* activity (Fig. S18)^66^.

These transplantation-associated changes prompted us to ask whether module activity remained associated with clone identity within graft tumors. Comparing distinct clones within the same graft tumor showed that several developmental cell-state-associated programs again displayed clone-associated expression differences after transplantation, whereas several cellular-process programs showed weaker clone-associated structure (Fig. 6E, Fig. S17E, Table S13). Intra-clone variance showed a similar overall pattern across primary and graft tumors, suggesting that some transcriptional programs regain clone-associated organization after transplantation-associated state changes.

Integrating clone-associated module differences in primary tumors with module changes between primary and late allograft tumors separated programs into distinct empirical groups (Fig. 6F, Table S14). For example, *immature_neuronal* showed both strong clone-association in primary tumors and high transplantation-associated change, whereas *catecholamine_production* was clone-associated but less sensitive to transplantation. In contrast, *interferon_signaling* showed weaker clone-association but strong transplantation-associated change, while *cell_cycle* showed comparatively low values in both metrics. Thus, tumor programs differ in the degree to which they are associated with clone identity and in their sensitivity to transplantation into an embryonic environment.

## Discussion

By leveraging well-controlled experimental conditions in the zebrafish model, our systematic analysis of intra- and inter-tumor heterogeneity allowed us to determine the spectrum of *MYCN*-driven NB tumor states consisting of 17 distinct gene expression modules, which represent both developmental cell state specific and general biological processes that are co-opted by tumor cells. Developmental benchmarking showed that, rather than occupying a single generic ‘sympathoadrenal’ state, the tumors span SCP-/bridge-like, sympathoblast-like and chromaffin-precursor-associated programs, with most zebrafish NB cells overall closest to more differentiated sympathoblast or chromaffin-precursor states. This developmental organization is consistent with prior human fetal-adrenal and stem-cell model studies that placed neuroblastoma within sympathoadrenal developmental trajectories and showed that oncogenic lesions can perturb neural crest / sympathoadrenal differentiation^23–30^. By combining this developmental-state framework with clonal barcoding, our study extends these observations by directly testing which tumor programs are associated with lineage history and which remain responsive to developmental context *in vivo*.

Overall, zebrafish NB cells show expected features of MYCN-driven neuroblastoma, including activation of ribosomal, cell-cycle and ADRN-associated neuroblastic programs. However, the activity of individual tumor programs varied strongly with clonal and anatomical context, indicating that MYCN-driven transformation does not impose a single uniform state even in this controlled model. The *immature_neuronal* module containing *alk* and *MYCN* was enriched in ventral AAC-region tumors, was associated with a less differentiated developmental bridge state^38,46^, and could also emerge after transplantation into an embryonic environment. While the function of tumor programs requires further investigation, this suggests that expression modules with potential relevance to high-risk disease can arise through non-genetic, context-dependent mechanisms *in vivo*.

In our clonal analysis, we found that modules associated with general cellular processes tend to exhibit a high propensity for context-responsive state reconfiguration, while modules associated with sympathoadrenal tissue development – and thus likely the cell state of origin – are mostly clonally determined. Assessment of clones in space showed that cells from multiple clones are spatially intermixed. Therefore, their distinct states are driven by clonal origin rather than local spatial positioning within the tumor tissue. These distinct clonal states may reflect remnants of the cell of origin or, alternatively, divergent states induced by MYCN during neoplastic transformation that subsequently stabilize. Their stable activation at an early timepoint in tumorigenesis suggest that the differentiation state and location of the cell at tumorigenic transformation are important factors in determining tumor phenotype. It is conceivable that specific tumor cell states arise from distinct effects of *MYCN*-overexpression on different cell of origin states, or that tumor cell state diversity is shaped by the differentiation status of the cell of origin, with less differentiated progenitors giving rise to a more diverse tumor cell population. Addressing these questions will be important for advancing our understanding and the treatment of NB.

Transplantations allowed us to determine how tumor cells from the same cell of origin behave in a different environment. Interestingly, some cellular process modules, as well as specific developmental cell-state-associated modules were particularly affected by transplantation. In our established developmental framework, these changes are consistent with movement along a sympathoadrenal differentiation axis and/or selective expansion of state-competent cells, rather than proving direct one-step fate conversion in every case. Within fully developed graft tumors, many developmental cell-state-associated programs were once again clone-associated, suggesting that transplantation-associated state reconfiguration is restricted to a distinct time window, after which cellular states stabilize. Overall, this analysis showed that even initially stable, clone-associated tumor programs can be reshaped upon exposure to a different signaling environment. Mechanisms of plastic adaptation have also recently been proposed to drive NB metastatic dissemination and emergence of drug tolerant persister cells^67,68^. Thus, understanding the molecular mechanisms that drive state transitions, by, e.g. perturbing transcription factors or signaling pathways associated with specific modules, will be a crucial step toward blocking transitions into more aggressive states.

## Limitations

While we did not detect mutations by WES or WGS, we cannot rule out that undetected mutations may influence the observed states. However, the observed sustained polyclonality and the alteration of tumor states upon transplantation suggests that modules are largely determined in a non-genetic manner. Furthermore, we only consider MYCN-driven NB; while this represents an important high-risk disease type, other NB genetic subtypes may behave differently. A related limitation is that the zebrafish models used here express *MYCN*, and in the MYCN;LMO1 model also *LMO1*, under control of the *dbh* promoter. Normal *dbh* expression in zebrafish is localized to the sympathetic ganglia, the developing IRG, and the AAC^17,38^. Thus, transformation is experimentally restricted to *dbh*-expressing noradrenergic / sympathoadrenal-lineage cells within these specific anatomical structures. This approach does not address whether earlier neural crest, SCP-like or other developmental populations could act as cells of origin under different oncogenic conditions. At the same time, this restriction is advantageous for the central question of our study, because it limits the range of initiating populations and thereby enables a controlled analysis of how transcriptional programs differ between clonally distinct tumor founders within a defined sympathoadrenal lineage context.

Because site-resolved normal reference data for the AAC, IRG and sympathetic ganglia are not currently available in zebrafish, our developmental annotation relies on public whole-body developmental atlases and cross-species comparison to human sympathoadrenal references; we therefore use the zebrafish developmental labels as an interpretive framework rather than as strict one-to-one cell-type assignments.

Here, we used spatial transcriptomics to assess the spatial distribution of clones within tumors. In a proof-of-concept experiment we found that clones were spatially intermixed in an individual tumor. Larger numbers of samples and more advanced analysis of spatial gene expression patterns would be required to investigate the effect of the TME in more detail.

Finally, while we can associate tumor states with human disease progression, we do not have a direct measurement of a tumor state driving tumor aggression. Additionally, the set-up of our allografting experiment is not suited for a quantitative assessment of selective advantage of individual tumor states. To address such questions, it would be necessary to analyze tumor state changes upon treatment e.g. in patient-derived xenografts and to identify strategies for inducing targeted state transitions.

## Acknowledgements

We thank Jana Richter for help with cloning and zebrafish line establishment, and Jutta Proba for help with allografting experiments. We acknowledge Janita Mintcheva for support and helpful discussions in the establishment of the Open-ST workflow for zebrafish tissues. We also thank Anika Neuschulz and Frank Westermann for helpful feedback on the manuscript. Furthermore, we would like to thank Chris McGinnis and Prof. Zev Gartner’s laboratory for generously providing a batch of reagents for MULTI-seq. The transgenic lines tg(*dbh:MYCN*,*dbh:EGFP*) and tg(*dbh:LMO1*,*dbh:mCherry*) were a kind gift from Thomas Look (Dana Faber Cancer Institute, Boston, USA). We would further like to acknowledge support from the MDC/BIMSB Protein facility and the MDC/BIH Genomics facility as well as the zebrafish facilities at MDC/BIMSB and Charité.

This work was supported by the Deutsche Forschungsgemeinschaft (DFG, German Research Foundation) within the Collaborative Research Center CRC1588 (project number 493872418, to J.P.J, A.H.H., A.G.S., N.R., D.B.), the European Research Council (ERC CoG 101043364 to J.P.J.) the DFG Forschergruppe FOR5628 (project number 513752256, to A.H.H.) and the Bruno and Helene Jöster Foundation within the collaborative research grant "Tumor Evolution and Plasticity in Childhood Cancer - TEP-CC". A.S. was supported by an EMBO Long-Term Fellowship (ALTF 972-2022) and a Marie Skłodowska-Curie Actions Postdoctoral Fellowship (ID 101106181 sc-LAB2FATE) from the European Union’s Horizon Europe programme.

## Author contributions

J.P.J., B.S. and N.F. conceived and designed the project. P.O.-C. and N.F. designed, established and tested the dsRedLinrecorder transgene and fish line. J.K. and L.H. performed histology and fluorescence microscopy. N.F. performed bulk and single-cell sequencing experiments with support from A.G., J.K. and L.B.. N.F. developed the computational lineage data analysis approach and performed single-cell transcriptome and lineage data analysis. N.F. generated whole exome sequencing data, J.K. generated whole genome sequencing data and B.S. performed the data analysis. A.S. performed spatial transcriptomics experiments and analysis with support from M.S.. A.S. developed the experimental and computational approaches for lineage barcode detection in spatial transcriptomics with support from N.F.. A.G. curated and analyzed human NB scRNA-seq data and was supported by M.J. and D.B.. J.K., A.G., L.B., L.H. and A.I.H.H. established, optimized and performed tumor cell transplantations and supported extraction of allograft tumor cells for sequencing. L.B. performed the moscot-analysis. M.S. was supported by N.R.. J.P.J., B.S., A.I.H.H. and A.G.H. supervised the project. J.P.J., B.S. and A.I.H.H. guided experiments and J.P.J. and B.S. guided data analysis. N.F. and J.P.J. wrote the manuscript in close interaction with B.S., A.I.H.H. and A.G.H. and with input from all authors.

## Methods

### Ethics statement

All experiments were performed in accord with the legal authorities approved license ‘G 0325/19’ and ‘G0020/25’ and were conducted in accordance with the European Community Council Directive of November 24, 1986 (86/609/EEC).

### Zebrafish lines

Zebrafish (*Danio rerio*) were raised and maintained according to standard protocols at 28°C with a 14/10 hour light-dark cycle^69^. Experiments in this study used the zebrafish wild-type strain AB. For lineage tracing, we used the transgenic tg(*ubi:zebrabow-M*)^70^, tg(*bActin2:dsRed_LinRecorder*) and tg(*hsp70:dsRed_LinRecorder*) (created in this study). Transgenic neuroblastoma lines used were tg(*dbh:MYCN, dbh:EGFP*) and tg(*dbh:LMO1, dbh:mCherry*)^31,32^.

Adult zebrafish of random sex were included in an experiment, when they had apparent growth of a dissectible tumor. Fish were euthanized immediately before tumor dissection by hypothermic shock as described by Wallace et al.^71^.

### Histological staining and imaging

Specimens were fixed in 4% phosphate-buffered formaldehyde (Labochem, L01-LC64701) for 48 hours, then washed in PBS and decalcified in 0.25M EDTA for 24 hours. Sections were mounted on microscope slides, deparaffinized, rehydrated, and stained with hematoxylin (Agilent Dako, CS70030-2) and eosin (Sigma-Aldrich) in Coplin jars following this protocol: 5 min xylene, 10 min xylene, 5 min 95% EtOH, 2x 2 min 80% EtOH, 5 min ddH_2_O, 6 min hematoxylin, dip into tap water, 6 min running tap water, 3 min eosin, dip into tap water, 1 min 70% EtOH, 1 min 80% EtOH, 1 min 95% EtOH, 2x 1 min 100% EtOH, 3 x 5 min xylene. Sections were mounted with Eukitt quick-hardening medium (Sigma-Aldrich, 03989) and a glass coverslip.

### Immunohistochemistry

Fixed and mounted specimens were deparaffinized in xylene (2 × 5 min) and rehydrated through a graded ethanol series (100% for 2 × 5 min, 70% for 5 min) and rinsed in ddH2O. Permeabilization was performed using 0.3% Triton-X (Fisher BioReagents, 10254583) in TBS-T for 5 min. For antigen retrieval, slides were heated in citrate buffer via microwaving for 20 min (sub-boiling) and cooled for 30 min. Nonspecific binding was blocked with 1% BSA in TBS for 1 h at RT in a humidity chamber. Slides were then incubated with an anti-N-Myc primary antibody (1:300; Santa Cruz, sc-53993) in 1% BSA/TBS at 4°C for 48 h in a humidity chamber. After washing in TBS-T (2 × 5 min), a Goat Anti-Mouse Alexa Fluor® 488 secondary antibody (1:300; Abcam, ab150113) was applied for 1 h at RT in the dark. Following a 10-min TBS-T wash, sections were counterstained with DAPI (1:1000 in PBS; Thermo Fisher Scientific, 10184322), washed twice in PBS (3 min each), and mounted with Prolong GOLD Antifade (Invitrogen, P36934). Slides were cured for 24 h at RT and stored in the dark at 4°C until imaging.

### Cloning of the lineage tracing recording cassette

The dsRedExpress coding sequence was sequence-optimized for zebrafish. We designed the recording cassette by placing three 23 bp sequences (including PAM) from RFP that had been tested for CRISPR/Cas9 editing before in an array interspersed by 7 bp spacers^33^. Sequences containing multiple restriction sites were placed on either side of the cassette. We further added the 10x capture sequence 1^72^ downstream of the cassette to enable more efficient capturing of transcripts in the 10x 3’ GE assay. Both sequences were synthesized by Twist Bioscience. The dsRed and the recording cassette were then inserted consecutively into the MCS of pME with the NEBBuilder HiFi DNA Assembly (NEB, E2621L), after linearizing the vector with KpnI+HindIII or BamHI+XbaI, respectively. The transgene was then cloned downstream the bActin2- or hsp70-promoter and upstream of a polyA-signal by Gateway LR reaction (Thermo Fisher, 11791020) into a pDest carrying the Tol2-TIRs for insertion of transgenes into the zebrafish genome. Finally, two integration barcodes were inserted into the transgene flanking the recording cassette. A 7-base-pair random sequence with a stable G in the middle flanked by 20-basepair overhangs complementary to the integration site was obtained as a single-stranded DNA oligo (IDT). The plasmid was linearized with EcoRI upstream of the cassette and the barcode oligo was inserted using the NEBBuilder HiFi DNA Assembly. A plasmid library with high barcode diversity was isolated from transformed bacteria. This process was repeated to integrate a second barcode downstream of the recording cassette after linearizing with NruI. The final plasmid library was subjected to Sanger sequencing to confirm a near-complete insertion of the barcode and the presence of barcode-diversity in the library.

### Generation of a lineage tracing line

The plasmid (at 6.25 ng/µl) was injected together with Tol2-mRNA (25 ng/µl) in 2 nl droplets into the yolk of wildtype zebrafish at the one cell stage. Larvae showing strong and widely spread red fluorescence in the body at 48 hpf to 5 dpf were selected for raising. Adult founders producing large fractions of red-fluorescent offspring were selected. Individual larvae were genotyped by next generation sequencing to identify integration barcodes and thereby number of integrations per larva. Two pairs of one male and one female founder each conferring several integrations with distinct integration barcodes to their offspring were bred to produce two F1-lines. F1 fish were mated with the transgenic NB lines for the lineage tracing experiments shown here.

### Cas9 and sgRNA injections for high-throughput lineage tracing and bulk barcode creation dynamics

For experimental batches 1 and 2 (Fig. S7H), we used a similar approach for sgRNA/Cas9 preparation as described before^33,34^. For experimental batch 3 Alt-R crRNAs (100 µM) were ordered from IDT and annealed to the constant tracrRNA (100 µM, IDT) individually by incubation of the mix at 85 °C for five minutes, followed by cooling at room temperature and subsequently on ice. Hybridized gRNAs for different targets were then pooled in multiple batches at the desired ratio. SpCas9 (MDC protein facility) was diluted to 26.8 µM in Cas9 freezing buffer (20 mM Tris-HCl pH7.5, 600 mM KCl, 20 % glycerol; all nuclease-free). Each gRNA-pool was mixed with an equal volume of diluted spCas9 and guides were loaded onto the protein by incubation at 37 °C for five minutes. Batches of loaded ribonucleoprotein complexes (RNPs) were then mixed at the desired ratio, aliquoted and frozen at −80 °C. 2 nl of freshly thawed RNPs were injected into one cell stage offspring of a cross between a transgenic lineage tracing line and tg(*dbh:MYCN, dbh:EGFP, dbh:LMO1, dbh:mCherry*). Successfully injected specimens were selected after 48 hours based on loss of pigmentation induced by cleavage of the tyr-CDS by Cas9 with a gRNA included in each mix.

### Sample collection and library preparation for bulk lineage barcode creation dynamics

Tg(*bActin2:dsRed_LinRecorder*) zebrafish were injected with 2 nl RNPs at the one-cell stage. Batches of embryos or larvae were transferred to Eppendorf tubes at selected time points. For RNA-extraction, zebrafish samples were homogenized in TriZol reagent (Thermo Fisher, 15596026) and RNA was extracted following the manufacturer’s instructions. Up to 3 µg of RNA were used as input for reverse transcription with SuperScript IV (Thermo Fisher, 18090010) and a poly-dT primer as used for CELseq2^66^, but with a 22 bp UMI. After second strand synthesis, cDNA from multiple samples was pooled and cleaned up as in the CELseq2 protocol^66^. Each target sequence of interest was enriched by a three-round nested PCR approach with NEBNext High Fidelity Master Mix (NEB, M0541) using target-specific primers, thereby also introducing the overhang sequences needed for sequencing. For DNA-extraction, zebrafish samples were incubated in 50 µl lysis buffer (10 mM Tris-HCl pH 8.0, 1 mM EDTA, 0.3 % Tween-20, 0.3 % Triton-100; all nuclease-free) with 1 mg/ml proteinase K (Invitrogen, 25530049) at 55 °C overnight, followed by 30 minutes at 85 °C to inactivate proteinase K. gDNA was precipitated using isopropanol, washed twice with 70 % ethanol and finally resuspended in 10 mM Tris-HCl pH 8.0. gDNA was directly used as input for a two-round nested PCR approach using gene-specific primers. All libraries were sequenced on Illumina NextSeq2000 200 bp kits (R1 28 cycles, R2 155 cycles).

### Analysis of lineage scar creation dynamics

Sequencing reads from individual libraries were demultiplexed using bcl2fastq (v.2.19.0.316). A custom python script was used to further demultiplex reads from each individual sample based on barcodes introduced during PCR and sequenced on a non-index read. FASTQ files were aligned to references of the target genes using bwa mem (v.0.7.17)^67^. Sequences with less than three reads were removed. Reads were shortened to a sequence identifier 30 bases around the expected Cas9 cut site for endogenous targets and to a 92 base sequence identifier covering all three target genes for dsRedLinRecorder. Sequence identifiers matching the reference sequence were classified as wildtype, while all other sequence identifiers were classified as edited by Cas9. These assignments were used for calculation of the fraction of wildtype UMIs or wildtype reads for RNA-based and DNA-based libraries, respectively.

### Whole exome sequencing

Genomic DNA from tumor and control tissue (clipped fins of the same fish) was extracted with the DNeasy Blood & Tissue Kit (Qiagen, 69504) and WES libraries were constructed with the SureSelect XT HS2 DNA Reagent Kit (Agilent, G9981A) following manufacturer’s instructions with zebrafish-specific probes (SSXT Zebrafish All Exon, Agilent 5190-5450). Libraries were sequenced on the NovaSeq system (R1 150 cycles, R2 150 cycles).

FASTQ files were trimmed and aligned to the zebrafish genome GRCz11 using bwa mem and read duplicates were removed. Somatic mutations for a tumor sample taking into account the individual’s matched healthy tissue were called using GATK’s Mutect2^73^. These variants were used to calculate variant allele frequencies.

Segmental copy number variants were called with the CNVkit software^74^. The genome was first split into bins containing an equal number of bait-targets (excluding centromeric and telomeric regions), in which reads were piled up. Pile-ups were compared between tumor and matched control samples. For calling CNVs the in-built circular binary segmentation approach was used^75,76^.

### Whole genome sequencing

For whole-genome sequencing (WGS), tumor and matched control tissues (entire clipped caudal fins) were isolated from five *tg[dbh:MYCN, dbh:EGFP]* transgenic zebrafish. Genomic DNA was immediately extracted using the DNeasy Blood & Tissue Kit (Qiagen, Cat. No. 69504). PCR-free, tagmentation-based genomic DNA libraries were prepared at the Genomics Platform of the Berlin Institute of Health (BIH/MDC) using a LMN DNA PCR-Free LPK (96 SPL) kit (Illumina, #20041795). Short-read whole-genome sequencing was subsequently performed on 1 lane of 25B NovaSeq X Plus flowcell in PE150 (150+10+10+150) sequencing mode. Tumor and matched normal samples were sequenced to mean depths of 35-45x and 75-95x, respectively.

Sequencing data was aligned to the zebrafish genome GRCz11 using bwa mem. We used CNVkit to determine copy numbers of the tumor relative to the normal, with a minimum segment size of 1Mb.

### Tissue dissociation

Tumors were excised from adult zebrafish, carefully cleaned from non-fluorescent tissues and placed into ice-cold PBS. The tissue was then minced into small pieces. Tissue fragments were pelleted and resuspended in dissociation solution. Primary tumors used for scRNA-seq only were dissociated in 0.01% papain (Sigma-Aldrich, 1495005), 0.1% dispase II (Sigma-Aldrich, D4693-1G), 0.01% DNase I (AppliChem GmbH, A3778) and 12.4mM MgSO4 in calcium- and magnesium-free Hank’s balanced salt solution at room temperature for 25 min with trituration through a micropipette tip every 5 minutes. Primary tumors used for transplantations as well as larval samples and allograft tumors were dissociated in 500 µl 30 mg/ml type II Collagenase (Sigma-Aldrich, C2-28) (approx. 700 U/ml), 25 mM HEPES, 10% FCS. The tissue was incubated at 37 °C for 30 minutes with trituration through a micropipette tip every 5 minutes. 200 µl 2U/ml dispase II (Sigma-Aldrich, D4693-1G, in 50 mM HEPES, 150 mM NaCl, pH7.4) was added for the last five minutes of dissociation. Following dissociation with either protocol, the opaque cell suspension was pipetted on a 5mL polystyrene Falcon® round-bottom tube with a 30 µm-mesh cell strainer cap (Corning, 352003) pre-filled with 500 µl ice-cold PBS with 1 % BSA and centrifuged at 500 g at 4 °C for 5 minutes. Cells were washed once with 500 to 1000 µl ice-cold PBS depending on the cell pellet size. Cells were then pelleted again and resuspended in the desired volume for downstream processing. Live and dead cells were counted manually using a Neubauer counting chamber and Trypan Blue as a stain for dead cells. The dissociation protocol used for each tumor sample is listed in Table S2.

### MULTI-seq labelling

MULTI-seq lipid-modified oligos (LMOs) and co-anchor were kindly gifted by Zev Gartner’s lab and later acquired from Sigma Aldrich (LMO001)^36^. After counting cells in the suspension, an equal number of live cells from each tumor (200 to 500 thousand, depending on material availability) were incubated in 100 µl to 200 µl 200 nM LMO-Barcode-oligo mix in PBS on ice for five minutes. Then the co-anchor was added and mixed into the cells at 200 nM, followed by another 5 minute incubation on ice. The labeling reaction was quenched by addition of 800 µl PBS with 1 % BSA. Cells were pelleted and washed once before pooling in a large volume of PBS with 1 % BSA. Before droplet encapsulation or FACS cells were again passed through a 30 µm strainer.

### Gene expression, MULTI-seq barcode and lineage library preparation

Single cells from individual tumors or samples pooled after MULTI-seq labelling were counted manually using a counting chamber and Trypan Blue as a stain for dead cells. Cells were then used for scRNA-seq library preparation with the 10x 3’ GE kit (V3.0 or 3.1), aiming for 10,000 cells per library. The gene expression library was prepared following the 10x protocol. For MULTI-seq runs, the small fraction of the cDNA containing most barcode molecules was cleaned up in addition to the large cDNA fraction used for the gene expression library. To this end, the supernatant remaining during cDNA clean-up was incubated with SPRIselect beads (Beckman Coulter B23318, final ratio 3.2X) and isopropanol (final ratio of 1.8X) and cDNA was eluted in EB (Qiagen, 19086) after two ethanol washes. The small cDNA fraction was then used for library preparation using a two-step PCR protocol with primers amplifying the MULTI-seq barcode oligos specifically. For specific amplification of genetic lineage barcodes, we used an approach similar to the one previously described^33,34^. Briefly, a targeted library was prepared for each lineage tracing target gene using a three-round nested PCR approach with gene-specific primers and 100 ng 10x cDNA as input material. All libraries were sequenced on Illumina NextSeq500, NextSeq2000 or NovaSeq. 10x gene expression libraries and MULTI-seq LMO libraries were sequenced with a minimum Read-2 length of 90 cycles. Lineage tracing libraries were sequenced with a minimum Read-2 length of 120 cycles.

### ScRNA-seq data processing and mapping

All zebrafish single cell sequencing data was demultiplexed using bcl2fastq (v.2.19.0.316). Gene expression data was mapped to a zebrafish transcriptome (GRCz11, Ensembl release 92) extended to include all transgenes present in the fish lines used (*MYCN*, *LMO1*, *EGFP*, *mCherry*, *dsRedLinRecorder*) using Cell Ranger (v.7.0.0). Lineage tracing libraries were further processed using custom pipelines as described below.

### Zebrafish NB transcriptome analysis

Gene expression data was analyzed using Seurat v.4.0.0. Cells with over 10 % mitochondrial transcripts (genes named ‘mt-’) were removed. Datasets derived from one individual sample using the standard 10x workflow were filtered to only contain cells with at least 250 distinct genes and 500 UMIs (Table S1). Cells from MULTI-seq datasets were filtered more leniently and subsequently assigned to a sample of origin using the classification approach described in the following section. Cells that could not be assigned a sample of origin or that were classified as doublets in this process were removed, thereby ensuring that only Bonafide cells are kept for downstream analysis.

The data was then processed using the standard Seurat pipeline with log-normalization, scaling and identification of highly variable genes. The list of highly variable genes was filtered to remove batch-associated genes (genes expressed in 80 % of cells from a dataset and log2FC > 0.2 compared to cells from all other datasets) and known cell cycle markers^72^ (translated to zebrafish genes using https://www.flyrnai.org/diopt) to reduce their impact on dimensionality reduction and clustering. This was followed by PCA and selection of a suitable number of components to use for Louvain clustering and UMAP. Differentially upregulated genes were used to assign a cell type to each cluster (Table S3). (Sub-) cell type assignment was further refined iteratively by sub-clustering all non-NB cells, followed by a further separation of blood / immune cells and other stromal cells (Fig. S1B). cNMF^50^ was used to identify gene modules active in each cell subset. Modules and differentially upregulated genes were used to assign cell types and specific functions. Marker genes for cell type classification were mainly taken from two previous zebrafish scRNA-seq studies^41,77^. Copy numbers were inferred from scRNA-seq data with inferCNV using default parameters using a zebrafish reference with addition of the transgenes as individual chromosomes.

### MULTI-seq sample demultiplexing

MULTI-seq libraries were first processed using the deMULTIplex^36^ pipeline to obtain a count matrix of observations of each sample barcode in each cell. Cells were then assigned to a sample of origin based on three different classification approaches. First, the deMULTIplex R package (https://github.com/chris-mcginnis-ucsf/MULTI-seq) was used to classify cells. Second, a manual thresholding approach was used, where cells are assigned to a sample of origin, if the associated barcode is found in them at a high frequency. Cells that passed the threshold for multiple barcodes were labelled as doublets, whereas cells below the thresholds for all barcodes were labelled negative. In the third approach, cells were classified according to the dominant barcode found in them. Cells were assigned to a sample, if the associated barcode had at least 2.3 times as many molecules as the second-most highly detected barcode and if the molecules from the dominant barcode made up over 40 % of the total molecules found. Cells that did not fulfill these criteria were classified as doublets or negative if their total amount of barcode molecules was above or below the mean of total barcode molecules across all cells, respectively. To make sure cell-sample-assignment was stringent, a consensus of the three approaches was taken. Only cells with a matching assignment in at least two of the classification approaches were classified according to this label. Cells with ambiguous classification across the three approaches were labelled doublets. Doublets and negative cells were removed from further analysis.

### Identification of sympathoadrenal cells in zebrafish developmental scRNA-seq atlases

Daniocell and Zebrahub zebrafish developmental scRNA-seq data were obtained as filtered count matrixes^45,46^. Each dataset was first subset for cells expressing at least one key marker gene of neural crest and sympathoadrenal development (*sox10*, *mpz*, *elavl3*, *elavl4*, *isl1*, *hand2*, *phox2a*, *phox2bb*, *dbh*, *th*, *chga*, *chgb* and in Zebrahub additionally *ascl1a*, *ascl1b*). We clustered these cells and calculated expression scores for marker genes of human fetal adrenal medullary development^23^. Clusters with substantial expression of at least one of these gene modules were manually selected, subset and sub-clustered again. This process was repeated multiple times. Additionally, clustered were inspected for expression of the individual sympathoadrenal marker genes. Finally, we annotated clusters based on expression of human developmental gene modules and individual marker genes as well as developmental stage of cells contributing to it. Clusters that could not be annotated were excluded. Assessment of the entire developmental atlas datasets showed that the selected groups of cells were located in few clusters and in close proximity to each other in the entire UMAP embedding.

### Pseudotemporal ordering and lineage inference

Developmental reference cells (Daniocell zebrafish sympathoadrenal lineage) were normalized, and the 2,500 most highly variable genes were retained for downstream embedding. Data were scaled, and PCA (50 components) followed by a shared nearest-neighbor graph (15 neighbors, 30 principal components) and UMAP embedding were computed using default scanpy parameters. Three complementary pseudotime estimates were calculated: CytoTRACE (via the CellRank CytoTRACEKernel), diffusion pseudotime (DPT; 15 diffusion components, 10 used for branching), and Palantir (10-component diffusion map, multiscale space, 500 waypoints, k = 30 nearest neighbors). The root cell for DPT and Palantir was defined as the cell with the highest CytoTRACE score among cells assigned to the earliest developmental stage. Agreement between pseudotime estimates and the annotated developmental stage was assessed by Spearman and Pearson correlation.

Cell-fate trajectories were inferred with CellRank2 (GPCCA estimator). A pseudotime kernel (Palantir pseudotime) was combined with a connectivity kernel in an 0.8:0.2 weighted sum to build the transition matrix. Macrostates were identified from the leading 20 Schur vectors and partitioned into 13 macrostates, annotated against the reference cell-type labels; macrostates corresponding to the same biological population were manually merged into 9 terminal states, and initial (progenitor) states were set manually (cycling progenitors, Schwann cell precursors) based on their pseudotime distribution. Fate probabilities toward each terminal state were computed for every cell and lineage-driver genes were identified per terminal state as the genes whose expression most strongly correlates with the corresponding fate probability.

Transfer to NB tumor cells: Developmental cell-type labels, pseudotime values, and per-cell fate probabilities were transferred from the reference to tumor cells by a cosine-similarity k-nearest-neighbor approach (k = 15) in a shared, jointly computed PCA space (30 components; top 2,500 shared highly variable genes after joint normalization of reference and query cells). Categorical cell-type identity was assigned to each tumor cell by majority vote among its k reference neighbors, with assignment confidence defined as the fraction of neighbors agreeing with the majority label. Continuous developmental quantities (pseudotime, fate probabilities) were transferred as distance-weighted (inverse-distance) averages over the same k neighbors, with fate probabilities renormalized to sum to one per cell.

### Identification of gene expression modules in NB cells and identification of consensus modules

To capture both intra- and inter-tumor variation in gene expression, we performed NMF on NB cells on three levels: individual tumor samples, individual MULTI-seq datasets and the entire dataset. Modules from all individual tumor samples or MULTI-seq runs were aggregated into recurring consensus modules following approaches similar to those described in^3,4,42^. Count matrices and highly variable genes lists as determined in Seurat for NB cells from a) an individual tumor sample, b) an individual MULTI-seq run or c) all samples were passed as input to cNMF^50^. We ran cNMF with output module numbers (k) from 5 to a) 30 for individual samples and MULTI-seq runs or b) 55 for the entire NB cell dataset. We ran cNMF with 200 iterations for each sample, from which the algorithm builds a consensus result and a measure of stability for the results obtained for a given number of modules k. A suitable k displaying good stability across NMF-iterations was chosen for each sample in the range of 5 to 15 modules per sample or MULTI-seq run.

For analysis of the entire NB cell dataset, we chose a k of 47, as this displayed a local maximum in stability over NMF-iterations, and would enable recovery of highly resolved gene expression programs that could possibly add to the results obtained from individual samples and MULTI-seq runs. To define final gene lists from the NMF-output matrices, genes for each module k were ranked according to their contribution weight. For each module, genes were added starting with the most highly weighted gene. If a gene had a higher rank in another module, it was flagged. Up to four flagged genes were allowed before stopping addition of genes to the final module. Modules were named by evaluating individual contributing genes and associated GO-term analysis in g:Profiler (https://biit.cs.ut.ee/gprofiler/) using ranked genes as input.

For consensus module generation, all modules from individual tumor samples (or all MULTI-seq runs) were compared to each other in terms of their gene content using Pearson correlation across all genes with a positive z-scored contribution to one of the modules. (Figure S3A). Modules with a Pearson’s R of at least 0.1 with at least two other modules were selected. Modules that passed this filter were clustered using hdbscan from the dbscan package^78^ and a consensus module was generated for each cluster by keeping genes found in over 25 % of all modules in that group. Each consensus module was named according to the contained genes evaluating individual genes and associated GO-terms. If necessary, we considered gene contributions to a consensus module determined as the median z-score gene weight across modules contributing to a consensus module (Table S7). Modules were further merged, if they were assigned the same function as well as modules with strong overlap in gene content. Finally, ambiguous genes that were found in multiple modules were removed: For each module, a gene that is found in a more prominent position in another module flagged. One such duplicate was allowed, but all duplicates from the second onwards were removed. The final list of modules was compiled by adding MULTI-seq derived modules to those derived from individual tumors, followed by modules derived from whole dataset NMF (Figure S3B). Modules that overlapped strongly with existing modules as well as modules that were only very spuriously expressed or could not be assigned a function were removed (Table S5).

### Gene module expression scoring

Two different approaches were used for gene module expression scoring. To classify cells into those that express a module and those that do not and to thus be able to determine the fraction of cells expressing a module in a given population, we scored the expression following the approach published by Barkley et al^3^. Briefly, the average centered expression of the module and 1000 random gene lists was calculated. The expression score for the module of interest was defined as −log10(fraction of random modules higher than module of interest) and rescaled linearly to [0,1]. Here, a score higher than 0.5 means that the module of interest scored higher than half of the random gene sets and this value was used as a cut-off to determine whether a module is expressed in a cell. To derive expression scores that can be used for differential module expression calculations, scores were calculated using AUCell^79^. AUCell uses the “Area Under the Curve” (AUC) to calculate whether a given list of genes is enriched in the expressed genes of a cell. The AUCell score is relative measure of gene module expression and can thus be used to compare module expression levels between cells. Scores were calculated in NB cells from each tumor sample or early allograft dataset separately. Only modules containing more than five genes present in the respective scRNA-seq dataset were considered.

### Translation of zebrafish to human orthologs and vice versa

Zebrafish gene modules were input into DIOPT (https://www.flyrnai.org/diopt) to get a list of orthologs. Only orthologs with the following metrics were kept: rank = high OR rank = moderate, best score = yes, DIOPT-score >= 10 (max = 19). Mitochondrial genes were manually translated using the same DIOPT criteria to make sure that current gene name versions are used. Human modules were similarly translated using DIOPT keeping matches with rank = high. Importantly, all comparisons between zebrafish and human gene modules were carried out in ‘human gene space’, as assigning a single human gene match to a zebrafish gene is more reliable than vice versa, due to the genome duplication in teleosts. Zebrafish NB modules translated to human orthologs are shown in Table S10.

### Human cancer gene expression modules

Published gene expression modules were retrieved from the indicated sources. In Fig. 2D and Fig. S6A, the signature ‘MYCN_upregulated’ corresponds to the signature determined as constitutively upregulated MYCN-targets across cell cycle phases in Ryl et al^54^. The PRC2-targets signature consists of the intersection of the 300 top EZH2-bound genes in a ChiP-seq assay from two human NB cell lines as reported in Chen et al^80^.

### Human NB and developmental scRNA-seq analysis

ScRNA-seq data of human fetal adrenal medullary cells was downloaded in processed form https://www.neuroblastomacellatlas.org^25^. Zebrafish NB module scores were calculated using AUCell.

Patient NB scRNA-seq data from four studies was collected as count tables from the NBAtlas^24,25,56,58,81^. Count tables from Patel et al. were downloaded from thehumantumoratlas.org^57^. The data was analyzed in Scanpy (v.1.11.5). Only datasets generated from whole single cells on the 10x Chromium platform were kept. Samples with a median UMI count below 1000 were removed. Only cells with more than 200 unique genes, more than 500 UMIs and with at most 25 % mitochondrial counts were kept. Gene names were harmonized across datasets and made unique using the R package HGNChelper (v.0.8.1). After merging datasets, genes with zero counts across all cells were removed.

Gene module expression scores were calculated for each NB cell using the AUCell scoring approach, as described above for zebrafish data. Inter-tumor module expression differences and intra-tumor module expression standardized variance were calculated as described for zebrafish NB clones below. Briefly, module expression differences between two tumors were determined with a Wilcoxon rank-sum test and the significance of expression difference was determined by comparing the test result to that of 1000 tests with randomly permuted cell group labels. Expression-standardized variance of module expression within a tumor was calculated together with the variance of all individual genes in the dataset using the Scanpy’s sc.pp.highly_variable_genes. No comparisons or individual tumors were filtered out before plotting due to the relatively small number of samples (38).

### Human bulk RNA-seq analysis

Bulk gene expression data for 498 NB samples from the SEQC cohort was retrieved from GEO (GSE49711) as log2(1 + FPKM). Genes expressed in less than four samples were removed. Gene expression data from the TARGET-NBL-cohort was retrieved from the GDC data portal (https://portal.gdc.cancer.gov/) as STAR gene counts. Information on *ALK* mutational status was retrieved from Brady et al.^59^. Entrez gene IDs were translated to gene symbols. If this introduced duplicated gene symbols, the one with the higher variance was kept. Genes with less than 10 counts in less than five samples were removed. Counts were then normalized with a variance standardized transformation (VST) as implemented in DESeq2 (v.1.30.1)^82^.

In all datasets, expression scores were calculated following an approach implemented by Decoene et al.^83^, similar to the approach developed for single cell data by Tirosh et al.^84^. Only modules containing more than five genes found in the respective RNA-seq dataset were considered. To test for significance, pairwise Wilcoxon rank sum tests between expression scores in groups were carried out and significance was adjusted for multiple comparisons using Bonferroni correction.

### Mapping and filtering of lineage barcode libraries

The lineage tracing libraries for endogenous targets were processed as previously described^34^. Briefly fastqs were aligned to individual references of the endogenous targets using bwa mem (v.0.7.17). Reads associated with a valid cell barcode present in the transcriptome library were kept. Subsequently, scar sequences were filtered to remove PCR and sequencing errors or sequences arising from doublets, following the assumption that there can be at most two distinct alleles of an endogenous target within a cell. Cells, in which one or two sequences made up 80 % or more of the gene-specific transcripts were kept and only these top sequences were kept.

The lineage tracing libraries for transgenic targets were processed similar to our previously described approach^33^. Briefly, sequencing reads were aligned to individual transgene references using bwa mem and only reads associated with a valid cell barcode in the transcriptome data were kept. Scar sequences with only one read were removed. Following this, for each combination of cell barcode and UMI, only the scar sequence with the highest number of reads was kept. Furthermore, sequences derived from sequencing errors were reduced by comparing sequences found within a cell and removing those that had a low Hamming distance to others and a comparatively low read number. Finally, scar sequences with a relatively low number of reads (determined by distribution of reads for all scar sequences that passed previous filtering steps) were removed.

Sequences derived from the dsRedLinRecorder were subsequently split by integration ID. Only sequences carrying a valid integration ID barcode were kept. These were determined as sequences that contain a G in the middle position and have a considerably higher number of reads than invalid sequences. As a given transgene integration in an individual cell can only carry a single scar sequence, ambiguous sequences and cells were removed. Only cells, in which a single scar sequence contributed to 60 % or more of the detected transcripts, were kept. In these cells, only the sequence with the highest number of UMIs was kept. Cells and sequences that passed these filters were used as input for the clone calling pipeline.

### Clone calling

Clone calling is illustrated in Fig. S7C-G and was performed for each fish individually, as the possibility that scar sequences are created in multiple fish hampers a joint analysis. Clone determination starts out with a separate analysis of each endogenous target gene, similar to our previously developed approach^34^. Briefly, the sequences were shortened to a sequence-ID of 30 bases around the CRISPR target site. Sequence-IDs that were only observed once across all cells were removed. First, only cells with two distinct alleles (one being wildtype is allowed) were kept. For each sequence-ID, the fraction of the total observations of this ID that occur in cells together with a given other sequence-ID is determined. If at least 80 % of observations of one or both sequence-IDs were found in a specific combination, this combination is kept. If possible, a hierarchy of a sequence-ID that was created first (‘parent scar’) and one that was created later (‘child scar’), was determined on the fractions of co-occurrence. Subsequently, cells with only a single sequence ID are assigned to a group defined by a combination of sequence IDs, provided that this single ID could be unambiguously matched to that combination. Finally, cells that have multiple UMIs of wildtype sequence IDs only are labelled as wildtype cells. Cells and sequence-ID combinations that passed these filters are passed on to clone calling based on all target genes.

For final clone calling, information from all targets is merged, with input for transgenic targets directly taken from scar filtering step. Here, each endogenous target is represented with one joint combination of two sequence-IDs. Each transgene integration (as distinguished by integration ID barcode sequence) is input as an individual target gene. Wildtype sequences are excluded. Each contributing sequence (or sequence combination) is hereafter called a seq-ID. The overlap in associated cell barcodes is calculated (Jaccard index) for each seq-ID pair and a threshold of 0.3 was determined to derive a binary adjacency matrix for all seq-IDs. This is used as input for an undirected graph, serving as a basis for clustering of seq-IDs. Cell barcodes and seq-IDs are aggregated by cluster, but this often leaves several distinct clusters with overlap in cell barcodes. Therefore, two overlap fractions are calculated for each cluster pair: the two clusters’ cell barcode intersection size divided by the total cell barcode count of either one of the clusters. The higher of the two values is kept. Based on this adjacency matrix, clusters are flagged for merging using an overlap threshold, set to 0.8 by default (i.e. 80 % of cells associated with one cluster are also associated with the other cluster). A cluster is flagged as ambiguous, if it overlaps with multiple other clusters, but those clusters share barcodes with each other only below a secondary threshold (default 0.6). Such ambiguity can arise from two scenarios: a) a lineage barcode (or combination) was created multiple times in independent events or b) a cluster is defined by lineage barcode(s) (‘parent’) that were created early and overlap with multiple clusters represented by lineage barcode(s) that were created later (‘child’). To account for the latter case and to avoid removal of many cells that only carry an early lineage barcode, larger clusters can optionally be treated as a ‘parent’ and smaller ‘child’ clusters can be merged into it. This option leads to the establishment of lower resolution clusters as used for analysis in Fig. 6 (Fig. S15A). If this option is not activated, ambiguous clusters are removed, e.g. two independent ‘child clusters’ would be kept, while the ‘parent cluster’ overlapping with both of them is removed, leading to higher resolution clustering (as used for analysis in Fig. 3 and 4). Remaining clusters flagged for merging are merged. Cell barcode overlap between these merged clusters is again determined by Jaccard similarity and if new clusters with significant barcode overlap (default Jaccard index of 0.3) have emerged, these are marked as ambiguous and removed. The remaining clusters are the final clones. Finally, cell barcodes that were assigned to multiple clones are removed.

Once clones had been determined, a seq-ID is classified as being clone-specific, if 90 % of cells it was observed in came from one specific clone in a given experiment. Clone-specific seq-IDs were later used as identifiers to match allograft-cells to primary tumor clones.

### Differential module expression analysis

Differential module expression analysis was performed in a pairwise manner between cells from two different groups. In the analysis of primary tumors, cells were grouped according to clone and tumor (sub-)sample to allow for comparison of clones within one tumor location and of cells from one clone in different sub-samples. In the analysis of allograft tumor cells, cells from the early allograft were grouped by clone only, whereas cells from the late allograft tumors were again grouped by clone and (sub-)sample to enable comparison between clones across time in one late graft tumor or between several graft tumors. For all comparisons, only groups of at least 10 cells were considered, where the two combined groups contained at least 25 cells. The differential expression score for each module was calculated based on module AUCell expression scores using a Wilcoxon rank sum test (as implemented in the Seurat function FindMarkers). To assess significance, group assignments of the tested cells were randomly shuffled 1000 times, while preserving group sizes, and the differential expression test was repeated for each shuffle. All differential expression values were ranked and significance *p* was determined as the rank of the test group of interest relative to the 1000 permutations and the p-value was calculated as this rank divided by 1,000. This means that *p* < 0.05 is equivalent to the test group of interest exceeding 95 % of random outcomes. In order to make results comparable between different modules, the differential expression score was normalized to the overall expression level of the module. To this end, the differential expression score for the comparison of interest was divided by the mean differential expression score of all random permutations for that module. Only results with *p* < 0.05 were considered. Furthermore, comparisons, in which a given module was expressed in less than 5 % of cells in both clones were removed.

### Calculation of module expression variance

In order to obtain module expression variance measures that are comparable between different modules, we used the expression-standardized variance values generated by Seurat’s FindVariableFeatures function (selection.method = ‘vst’) (Seurat v.4.0.0). First, counts for all genes in a module were summed to get raw module expression scores that were added to the gene count matrix. The counts matrix containing genes and gene module counts was then log-normalized using Seurat’s NormalizeData function. Log-normalized expression values were used as input for the FindVariableFeatures function, which fits the mean-variance relationship across genes and rescales observed variances by the expected variance at a given mean expression level. Expression-standardized variance was calculated per clone or other group of cells of interest.Groups of less than 30 cells were removed as well as groups of cells, which only had module expression in less than 5 % of cells.

### Tissue processing, library preparation and data preprocessing for spatial transcriptomics of zebrafish tumors

#### Samples

The section from spatial ‘Tumor 1’ was taken from an embedded piece of the same tumor used for generation of scRNA-seq sample ‘MYCN_lat_m17_3’. This tumor carried lineage barcodes. Spatial ‘Tumor 2’ corresponds to a tumor sample without lineage barcodes that was not processed with scRNA-seq.

#### Open-ST spatial transcriptomics and sequencing

Dissected entire tumors or pieces of tumor tissue (if another piece was used for scRNA-seq) were embedded in optimal cutting temperature compound (O.C.T., Tissue-Tek, 4583) in plastic cryomolds. The filled mold was frozen by placing it on a flat metal surface cooled down with dry ice. Frozen samples were subsequently stored at −80 °C. Cryosections were then cut at 10 µm thickness and mounted on Open-ST capture areas. Tissue handling and spatial barcoding were performed following the Open-ST protocol^62^. Brightfield images of H&E-stained sections were acquired using a Keyence BZ-X700 to assist with downstream image registration and background removal. Following cDNA elution, the whole transcriptome library was prepared following the Open-ST protocol. Gene-specific libraries were generated using a two-round nested PCR approach using target-specific primers. All final products were size-selected on a BluePippin HT system (Sage Science) Spatial transcriptomics libraries were sequenced on an Illumina NextSeq 2000 using a 200-cycle kit (R1: 37 cycles, R2: 191 cycles).

#### Alignment and generation of count matrices

Raw spatial transcriptomics data were processed and aligned using SpaceMake^85^, which produced a gene-by-spot count matrix from sequencing reads. Individual tiles were stitched, and expression was aggregated on a hexagonal grid with 5 µm diameter bins using custom Python code.

#### Image-based spot filtering

The brightfield image of the tissue section was used to create two images: one inverted in Fiji for alignment and one thresholded (black and white) for spatial filtering. Using a custom Python code, the first image was aligned to a synthetic transcriptomic image rendered by showing the number of spots aggregated into each hexagon of the grid. Manual landmarks were selected on both optical and spatial transcriptomic images. An affine or homography transformation was computed using OpenCV and applied to the binarized version of the optical image. Only spots falling within foreground tissue regions were retained for downstream analysis.

#### Transcriptomic filtering and normalization

Following image-based subsetting, spatial transcriptomic data were filtered to remove spots with fewer than 5 detected genes and genes expressed in fewer than 5 spots. Total counts per spot were normalized to 10,000 and log-transformed using log1p. Highly variable genes were selected using Scanpy’s variance-based method, retaining the top 2,500 genes.

### Spatial analysis of cell type and tumor modules

#### Cell type label transfer

Single-cell RNA-seq reference data were integrated on shared highly variable genes using Scanpy’s implementation of Harmony^86^, and spatial spots were projected into a shared PCA space. For each spot, a k-nearest neighbor model was used to infer a probability distribution over reference cell types, resulting in per-spot soft cell type scores.

#### Module scoring

Gene modules were quantified by computing the fraction of total expression per spot attributable to each module. For each spot, expression of all valid module genes was summed and divided by the total spot-wise expression.

#### Spatial correlation

To assess local co-variation of cell type or module scores, values were smoothed across spatial neighborhoods defined by a fixed Euclidean distance using a cKDTree search. Pearson correlations were then computed between smoothed scores across spots.

#### Proximity to tissue boundaries

To evaluate spatial positioning relative to tissue borders, connected component analysis was used to define tissue regions. We manually retained the three largest regions of the tumors. A Euclidean distance transform was applied to compute each spot’s distance to the nearest external boundary, and Spearman correlations were computed between distance and module scores within each region.

### Spatial clonal analysis

Spatial gene-specific libraries were used for spatial clonal analysis: Scar gene barcodes were extracted from read 1 and assigned to spatial coordinates based on their title and lane identifiers. Read 2 was aligned to reference scar genes using bwa mem, and a spatial barcode-scar sequence count matrix was constructed. Barcode matrices were filtered using the same imaging-derived tissue mask applied previously. Clone identities inferred from matched scRNA-seq data were transferred to spatial spots by aligning shared scar sequences (seqIDs) across modalities. Clone presence at each spot was binarized by thresholding to ≥ 1 supporting read per scar gene.

### Allogeneic transplantations into zebrafish embryos

The pool of dissociated tumor cells from multiple tumors was divided between the workflow for scRNA-seq and allogeneic transplantation. For transplantation, the cells from multiple tumors were counted and mixed. Cells were centrifuged through a 20 µm mesh filter at 500 g at 4 °C for 5 minutes. The supernatant was removed almost entirely and cells were resuspended in a tiny volume of PBS to keep the suspension very dense. Glass pipettes with a 20 µm outer diameter (BioMedical Instruments) connected to an air-pressure injector (IM-400) were used to inject 100 - 150 cells into each embryo 4 h after fertilization. Embryos were allowed to recover in E3 medium (5mM NaCl, 0.17mM KCl, 0.33mM CaCl, 0.33 mM MgSO4, pH 7.4) supplemented with 1% penicillin/streptomycin for 1 h at 28 °C before manually sorting for transplantation success on a M165 FC stereomicroscope (Leica Microsystems).

### Isolation of tumor cells from larvae via fluorescence activated cell sorting (FACS)

Larvae at 2 dpt (days post transplantation) were collected in batches of 15 to 20 and placed on ice for 10 minutes. Larvae were then washed once in ice-cold HBSS and dissociated with collagenase II and dispase as described above. If multiple larval batches were processed at once, cells were then labelled with MULTI-seq barcodes as described above. After quenching of the labelling reaction and thorough washing with PBS with 1 % BSA, cells were resuspended in PBS with 0.05 % BSA for FACS sorting (BD Biosciences FACSAria III). Cells were selected by first gating for GFP-fluorescence, followed by side- and forward-scatter gating. Cells were sorted into a cooled 1.5 ml eppendorf tube pre-filled and coated overnight with 500 µl PBS with 2 % fetal calf serum. Cells were sorted with a flow rate below 4 and an event-rate below 2000 events per second. Every 20 minutes, the sort was halted and the source cell suspension was vortexed. The receiving tube was closed and inverted before placing it back into the sorter. Sorting was done until either a) 50000 cells had been sorted into the tube, b) at least 20000 cells had been sorted and the cell suspension was close to running out, c) sorting time reached two hours. In the two latter scenarios, the gate sorting for fluorescence was inactivated, so that all live cells were sorted until 40000 total cells had been sorted into the tube. After sorting the receiving tube was inverted a few times before centrifugation at 4 °C, 1000 g for 5 minutes. Most of the supernatant was removed, leaving an estimated 10 µl around the cell pellet. The cells were resuspended after addition of 50 µl PBS and subsequently counted in a counting chamber prior to single cell encapsulation.

### Cell-cell communication analysis

We ran CellChat (v2.2.0) using the inbuilt zebrafish ligand-receptor interaction database with standard settings. For each sample group of interest (all primary tumors, all late allograft tumors or only late allograft tumors in ventral or orbital location), individual tumors were considered as biological replicates. We filtered predicted interactions to only retain those observed in at least four tumors in cell types with at least 10 assigned cells for primary tumors or in at least three tumors in cell types with at least 7 assigned cell types in late allograft tumors.

### Assignment of allograft cells to primary tumor clone

Lineage tracing target-specific libraries were processed as described for primary tumors. Sequences from endogenous target genes were shortened to a sequence-ID of 30 bases around the CRISPR target site. Cells with two distinct sequence-IDs for a given target were selected. For transgenic targets, valid integration ID barcodes were extracted and other sequences were removed. Thereafter, sequences were filtered, so that each cell retained at most one sequence per integration ID as described for the primary tumors. Combined sequence IDs from the endogenous targets as well as individual sequence-IDs from each transgene integration were used to match allograft cells to a primary tumor clone defined by one or more of these sequence-IDs. Here, we used primary tumor clones called at slightly lower resolution to increase the number of allograft cells that could be assigned to a clone (Fig. S17A-B). Ambiguous assignments of one graft cell to multiple primary clones were removed.

### Analysis of clonal state transitions across transplantation time points

To infer transcriptional state transitions across transplantation time points, we applied moscot (v0.4.2) to the clone-resolved single-cell RNA-seq multi_seq_07_6_8 and multi_seq_07_1_3^66^. Cells were grouped according to sampling time point using the sample_type annotation, comprising primary_tumour (PT), early_allograft (EA) and late_allograft (LA) cells. The analysis was performed on the integrated representation after dimensionality reduction by PCA on the integrated assay. Pairwise optimal-transport problems were constructed between consecutive time points, comparing primary_tumour to early_allograft and early_allograft to late_allograft cells. Transport maps were computed using the moscot BirthDeathProblem framework with a Sinkhorn solver, using unbalanced transport (tau = 0.95) and epsilon = 1e-2, and convergence was inspected for each transition. To summarize inferred transitions at the transcriptional program level, each cell was assigned to the gene module with the highest module expression score. Cell-to-cell transport probabilities were then aggregated by source and target module identity to obtain module-level transition matrices for each time interval. The resulting transition tables were exported and visualized as Sankey plots, with low-abundance transitions removed using a predefined filtering threshold where indicated. In parallel, cell-level transport-derived metrics, including transition entropy, concentration, maximum transition probability and progression-axis scores, were calculated to assess whether transitions were focused toward specific target states or diffusely distributed across the transcriptional landscape. To identify genes associated with inferred state transitions, gene expression values were correlated with transport-derived progression metrics separately for the primary_tumour-to-early_allograft and early_allograft-to-late_allograft transitions. Genes were ranked according to the direction and strength of their correlation with these scores, allowing identification of genes positively or negatively associated with focused transition behavior and progression along the inferred transport axis.

## Data availability

The sequencing data generated in this study is deposited in the Gene Expression Omnibus under GSE301974 (scRNA-seq data), GSE301860 (lineage tracing data from scRNA-seq), GSE301869 (spatial transcriptomics data).

## Code availability

Custom code is available through Github at https://github.com/junker-lab/neuroblastoma_plasticity.

---

**Fig S1:**
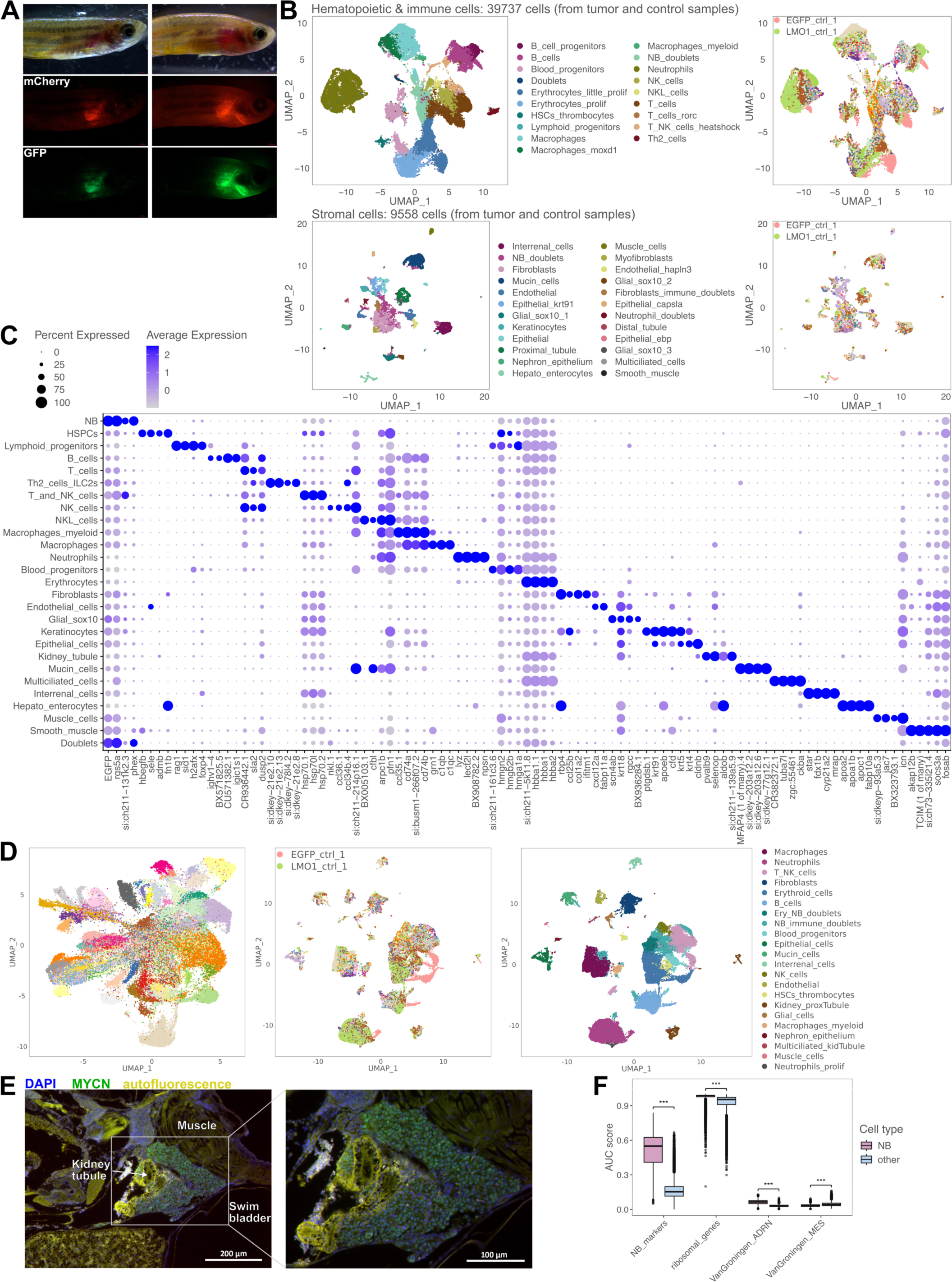
Information on zebrafish NB models, cell type composition and annotation. A) Stereomicroscopic images of two MYCN;LMO1 fish with a lateral tumor (both fish) and a ventral tumor (fish on the right). B) UMAP showing sub-clustered immune and blood cell types (upper plots) or sub-clustered stromal cell types (lower plots) with detailed sub-cell type annotation (left) and indication of the sample of origin (right). In the plots showing samples of origin, only the colors used for control samples are highlighted, as these contributed a large amount of immune and blood cells that separate slightly from tumor-derived cells on the UMAP. These cells are derived from the hematopoietic tissue of the head kidney. C) Dotplot showing expression level and prevalence of the top four cell type marker genes (as determined by differential expression analysis, Table S3) for each final cell type. D) UMAP of all NB cells (left) and all other cell types (middle and right) colored by the sample of origin (left and middle) or cell type (right). In the plots showing samples of origin, only the colors used for control samples are highlighted, as these samples show larger differences compared to TME cells. E) Immunohistochemistry for MYCN protein expression (green) in sagittal FFPE section of a 2 mpf old MYCN-fish with tumor in the interrenal gland (IRG) of the head-kidney. MYCN in green (IRG tumor), nuclei in blue, autofluorescence in yellow. F) Expression scores for the genes most differentially upregulated (‘*NB_markers*’ as in F) or most highly expressed (‘*ribosomal_genes*’) in NB cells, as well as for published ADRN/adrenergic and MES/mesenchymal gene signatures derived from human NB cell lines. Significance of differential expression between NB cells and all other cells was determined with Wilcoxon rank sum test (*** p < 0.0001).

**Fig S2:**
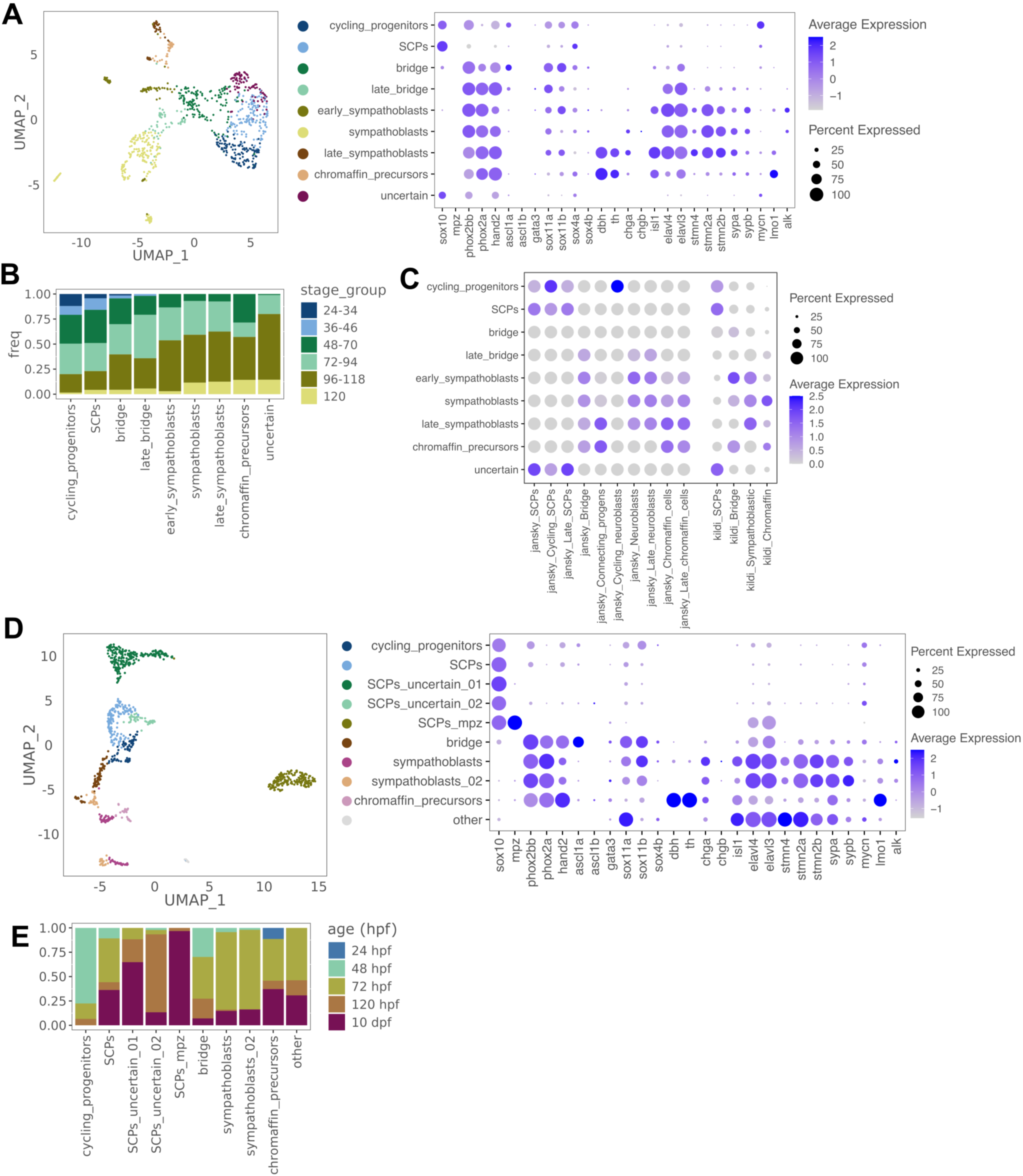
Identification of developmental sympathoadrenal populations in zebrafish scRNA-seq data. A) UMAP of putative sympathoadrenal cells and assigned cell types extracted from the Daniocell atlas. Expression of known marker genes of different sympathoadrenal developmental cell states is shown in the dotplot. B) The barplot shows fraction of cells of a given cell type extracted from a given embryonal /larval age (in hours post fertilization). C) Expression (AUCell scores) of human sympathoadrenal marker gene sets (from Jansky et al. and Kildisiute et al.^23,25^) in the zebrafish developmental cell types. D) UMAP of putative sympathoadrenal cells and assigned cell types extracted from the Zebrahub atlas^46^. Expression of know marker genes of different sympathoadrenal developmental cell states is shown in the dotplot. E) The barplot shows fraction of cells of a given cell type extracted from a given embryonal /larval age (in hours post fertilization).

**Fig S3:**
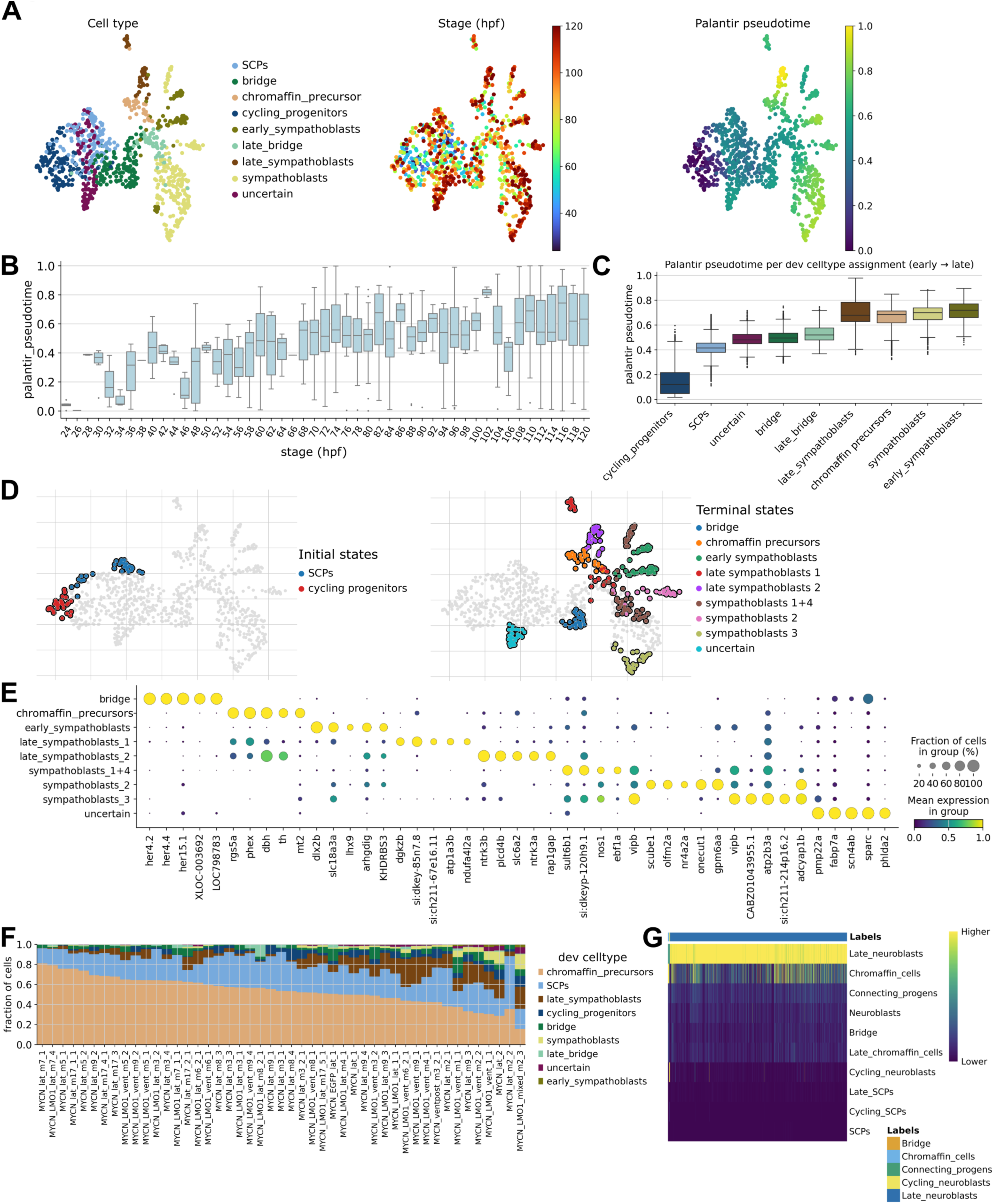
Zebrafish sympathoadrenal developmental pseudotime and correspondence to NB cell states. A) UMAP of sympathoadrenal cell types extracted from the Daniocell dataset colored by cell type, stage in hours post fertilization or Palantir pseudotime. B) Palantir pseudotimes assigned to zebrafish cells correlates with real developmental time determined by stage of the sample of origin in hours post fertilization. C) Boxplots showing Palantir pseudotimes assigned to developmental sympathoadrenal cell types. D) UMAP as in A colored by initial and terminal states in CellRank2-determined cell-fate trajectories. E) Five lineage-driver genes per terminal cell state as determined as the genes whose expression most strongly correlates with the corresponding fate probability. F) Fraction of NB cells from each tumor assigned to a given developmental cell type by transcriptomic similarity. G) SingleR label transfer from human fetal adrenal medullary cell types to zebrafish NB cells using cell type marker genes from Jansky et al^23^. The heatmap shows similarity scores for each developmental cell type in each NB cell (columns). Assigned labels are highlighted in the color legend above the heatmap.

**Fig S4:**
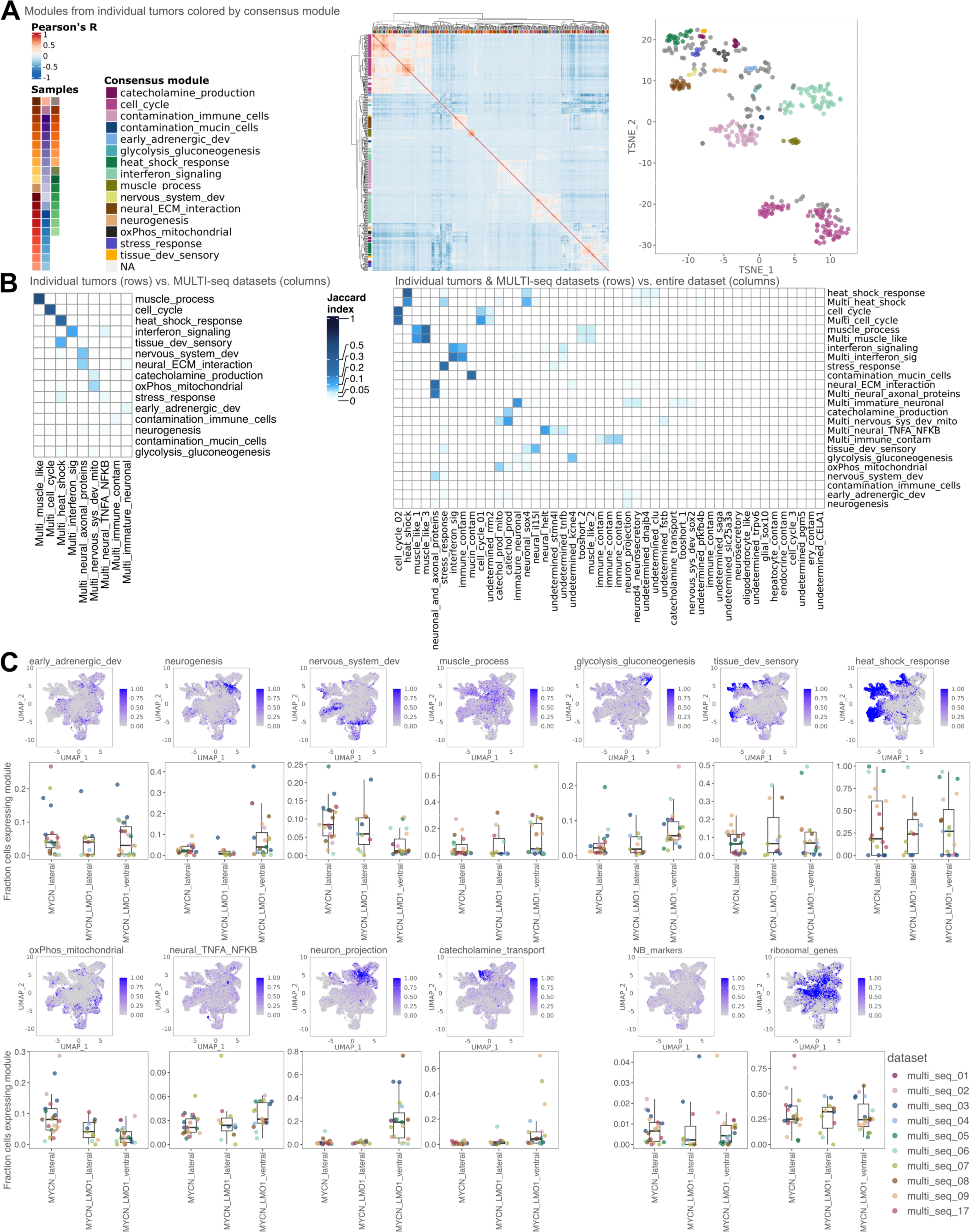
Detection of NB gene expression modules using a three-level NMF approach. A) Heatmap and T-SNE showing modules detected in the analysis of individual tumors plotted according to their pairwise correlation to each other (Pearson). Modules in the heatmap are annotated by the individual tumor sample they were derived from (color legend in top row) and by the consensus module they were assigned to after clustering with HDBScan (color legend on the left side). Modules in the T-SNE plot are colored according to the consensus module they were assigned to after clustering with HDBScan. Modules that were not assigned to any consensus module are labelled NA and colored in grey. B) Overlap in terms of gene content (measured by Jaccard index) for all consensus modules derived from the analysis of individual datasets (as in A), MULTI-seq datasets or classical cNMF analysis of all NB cells from the entire dataset (Table S4). C) UMAPs of NB cells with expression scores for those of the 17 curated consensus modules not shown in the main figure 2 as well as the signatures for NB cell differential and NB high expression (‘*NB_markers*’ and ‘*ribosomal_genes*’, respectively). Box plots and jitter plots show fraction of cells expressing a given module per tumor. Tumors are grouped by genotype (MYCN or MYCN;LMO1) and primary tumor location (lateral or ventral).

**Fig S5:**
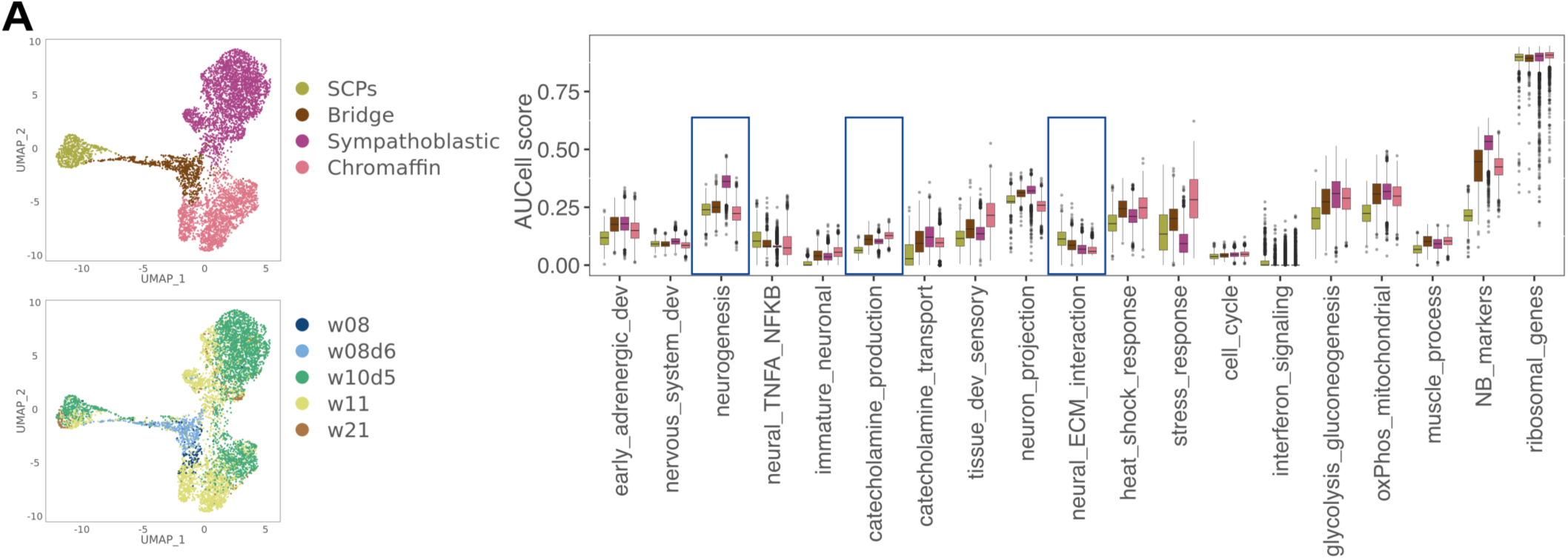
NB module expression in human developmental cell types. A) UMAP of human fetal adrenal cell types showing assigned cell type or sample age in weeks post conception (from Kildisiute et al.^25^). Boxplots show AUCell scores for the zebrafish gene modules in the four assigned cell types. Blue boxes highlight modules with tendency for higher expression in one of the four cell types.

**Fig S6:**
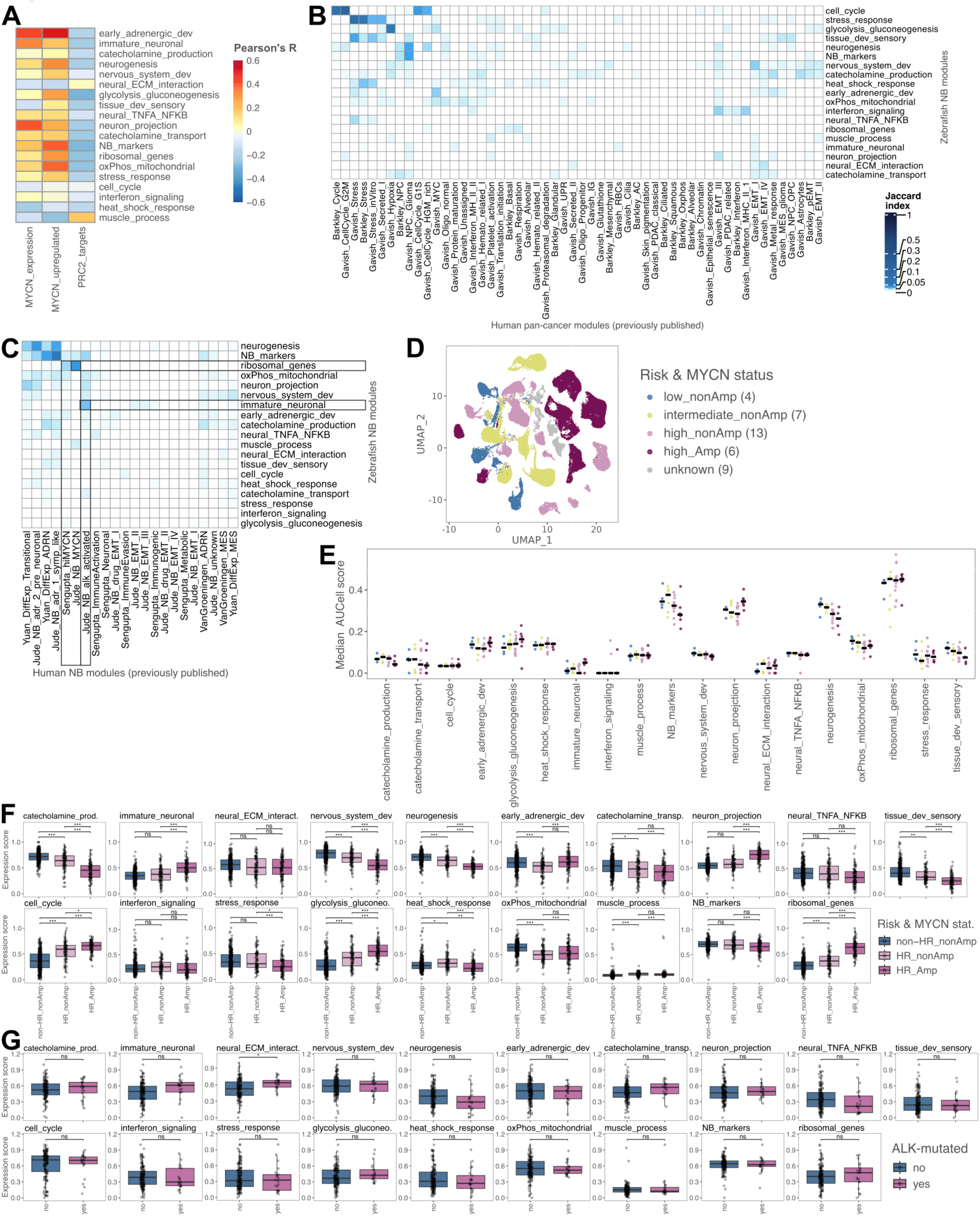
Similarity between zebrafish NB and human NB gene expression programs and expression of zebrafish NB modules in human NB. A) Correlation between expression scores of all zebrafish NB-derived gene modules (rows) and expression of *MYCN* (‘MYCN_expression’) or expression scores for human MYCN-driven or PRC2-target genes (measuring MYCN-mediated gene silencing via EZH2^80^), across all zebrafish NB cells. B) Gene content overlap as measured as Jaccard index between zebrafish NB gene modules (rows) and gene expression modules derived from analyses of many human cancer types (columns, named by study of origin and module function). C) Gene content overlap as measured as Jaccard index between zebrafish NB modules (rows) and gene expression modules derived from human NBs (columns, named by study of origin and module function). Color scale as in B. D) UMAP of integrated human NB cells from 39 tumors colored by risk (low, intermediate, high) and MYCN mutational status (non-amplified or amplified). E) Median per-tumor module expression score for tumors shown in D (excluding those with missing metadata), with tumors grouped by MYCN-status and disease risk. Black lines indicate the median value across tumors from a given group. F) Expression score of zebrafish NB modules in bulk RNA-seq data from the SEQC NB cohort (n = 498), grouped by risk (HR = high-risk, non-HR = non-high-risk) and MYCN-status (non-amplified or amplified). Significance of inter-group differences in E and F were tested with a Wilcoxon rank sum test (ns = non-significant, * p < 0.01, ** p < 0.001, *** p < 0.0001). G) Expression score of selected zebrafish NB modules in bulk RNA-seq data from the TARGET NB cohort (n = 151), grouped by ALK mutational status.

**Fig S7:**
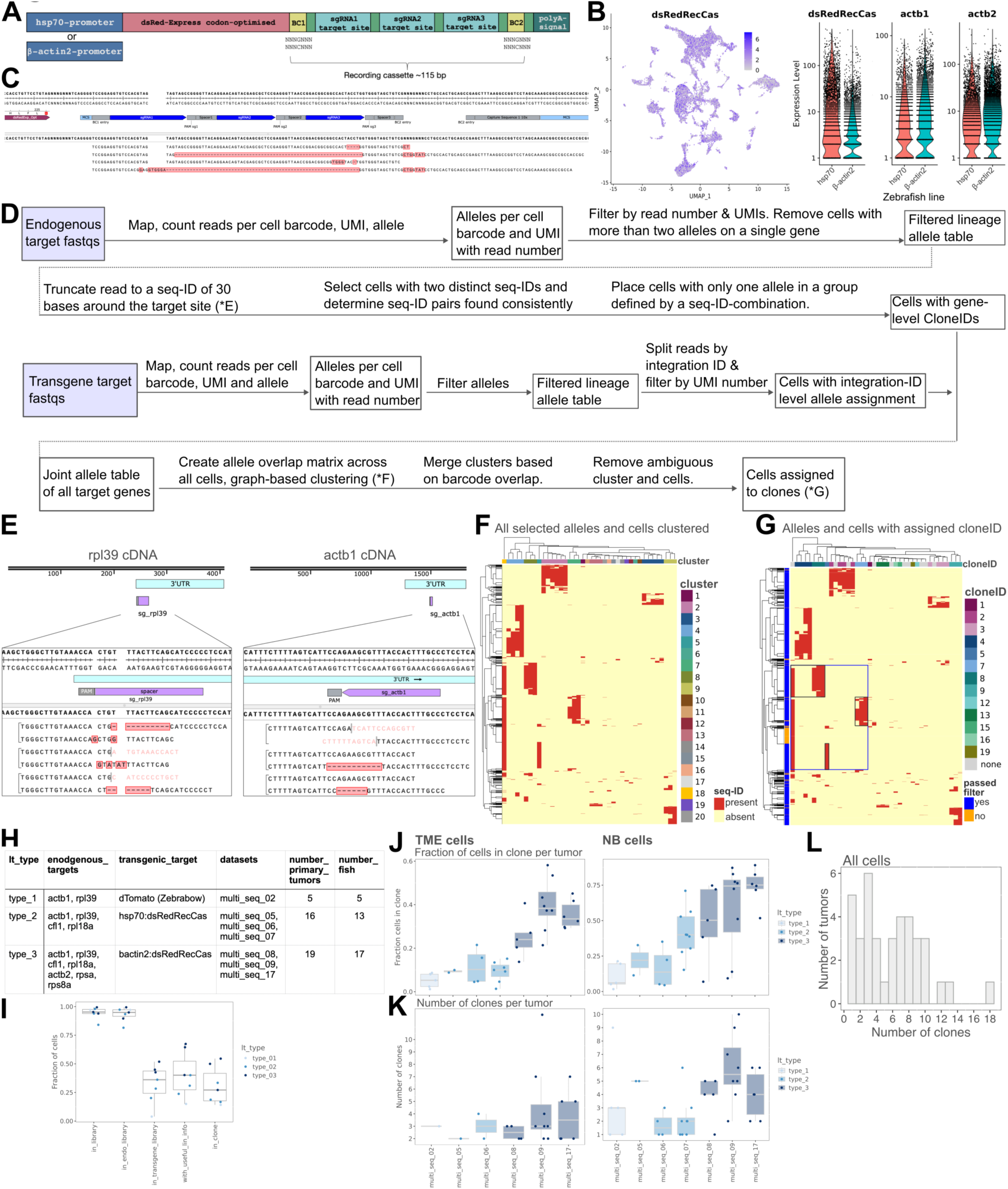
Lineage tracing target design, data processing, clone calling and clone detection statistics. A) Lineage tracing cassette design: Three Cas9-target sites are encoded in the 3’ UTR of a dsRed-gene sequence, optimized for expression in zebrafish, flanked by two 7 basepair barcode sequences (BC1 and BC2) that function as integration IDs to distinguish multiple genome insertions of the transgene. B) Expression levels of dsRed-RecCas (considering reads from the full length of the gene) and endogenous *actb1* and *actb2* in scRNA-seq data derived from zebrafish larvae of the hsp70:dsRedLinRecorder or bActin2:dsRedLinRecorder lines. Violin plots show expression of the transgene and two highly expressed endogenous genes in the two lines. C) Sequence of dsRed-RecCas with aligned CRISPR/Cas9-edited example reads. Edits include small alterations at one cut site as well as larger deletions induced by cutting of multiple target sites. D) Workflow for pre-processing and filtering of lineage target sequencing data from endogenous and transgenic targets as well as clone calling. E-G show data representations of an example dataset at various stages of the workflow and are referenced in the corresponding steps in the flowchart. E) Pairs of sequence-IDs found in three different cells on *rpl39* (left) and *actb1* (right) 3’ UTRs. F) Heatmap of cells (rows) and alleles (columns) clustered by their presence (red). Data represents all cells from one fish that passed filtering for at least one target gene and seq-IDs from all target genes (columns). Seq-IDs were clustered based on cell barcode overlap with clusters indicated above the heatmap. G) Heatmap as in F. Final clone-assignments are indicated above the heatmap with seq-IDs that did not pass filtering shown in grey. Cells that could unambiguously be assigned to a single clone are highlighted in blue on the left side of the heatmap. The heatmap represents high resolution clone calling. Cells that could be assigned to a clone are marked blue in the color bar left of the heatmap. One seq-ID is shared between many cells from multiple different clones (large blue box). Cells that only carry the parental seq-ID are filtered out (orange on the side of the heatmap), whereas cells with additional seq-IDs are grouped into sub-clones (small black boxes). In order to increase the number of cells and lineage barcodes used, the clone calling resolution can be lowered, so that all cells in the large box would be merged into one clone. H) Conditions used for lineage tracing injections in three experimental rounds. Different endogenous 3’ UTRs and transgenic loci were targeted in the conditions. Adult fish from one injection round were used for one or multiple MULTI-seq runs (datasets). I) Fraction of cells per MULTI-seq dataset (dots) with lineage information at different steps of the filtering and clone calling pipeline (in library = cells with at least two UMIs with more than one read each in all targeted libraries; in_endo_library = cells with at least two UMIs with more than one read each in targeted libraries for endogenous genes; in_transgene_library = cells with at least two UMIs with more than one read each in targeted library for transgene target; with_useful_lin_info = cells with a valid combination of alleles on any endogenous target and/or a valid lineage barcode (uncut) on a transgenic target; in_clone = cells that could be assigned to a high resolution clone). lt_types as in H. J) Fraction of cells assigned to a high-resolution clone per tumor (dots) grouped by MULTI-seq dataset considering only TME cells or only NB cells (right). lt_types as in H. K) Number of high-resolution clones per tumor (dots) grouped by MULTI-seq dataset considering only TME or only NB cells (right). lt_types as in H. L) Number of clones detected considering all cells (TME and NB) per individual tumor (as in Fig. 3F for NB and TME cells separately).

**Fig S8:**
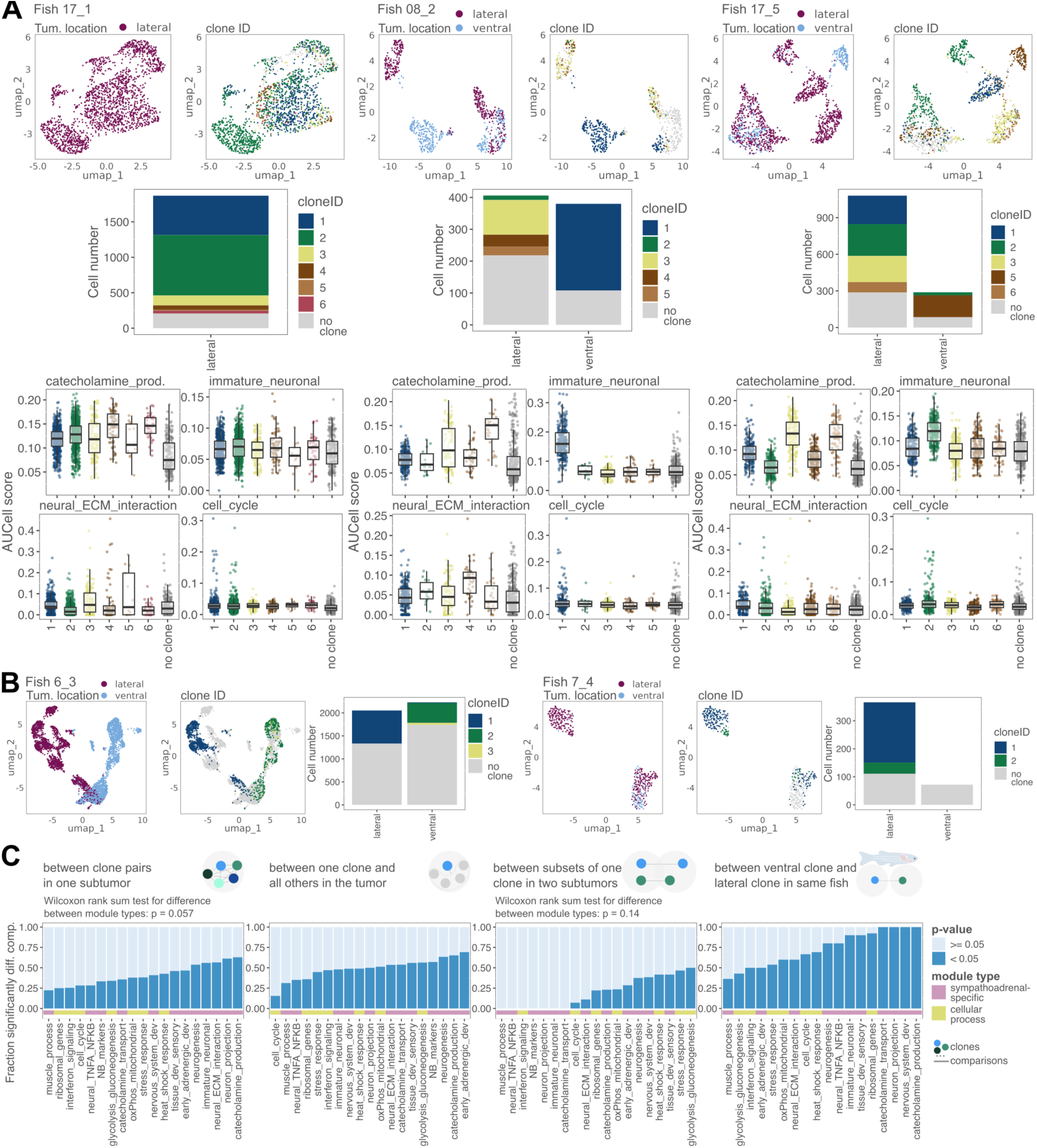
Lineage-dependence of gene expression in zebrafish NB tumors. A) UMAPs of NB cells taken from three example fish colored by the tumor location or the clone-ID. Fish 17_5 and 08_2 had a ventral and lateral tumor each that show clear separation on the UMAP as well as distinct clonal composition. Clonal composition for each tumor site is also shown in barplots. Expression scores of selected gene modules are shown as box- and jitter-plots. Inter-clone expression differences are particularly strong for clones found in distinct tumor locations. B) UMAPs of NB cells as in A for the two remaining fish that had a lateral and ventral tumor each. Clonal composition for each tumor site is shown in barplots. C) Fraction of significantly different module expression scores between groups of cells in four different comparisons. First pairwise differences between clones in a single tumor (sub-)sample are shown (as in Fig. 4C). Comparison of one clone vs. cells from all other clones in the same (sub-)sample show similar results. Comparison between cells from the same clone found in different sub-locations of the tumor shows overall lower differences. Comparisons between different clones found in the same fish, but in ventral and lateral locations show overall larger differences.

**Fig S9:**
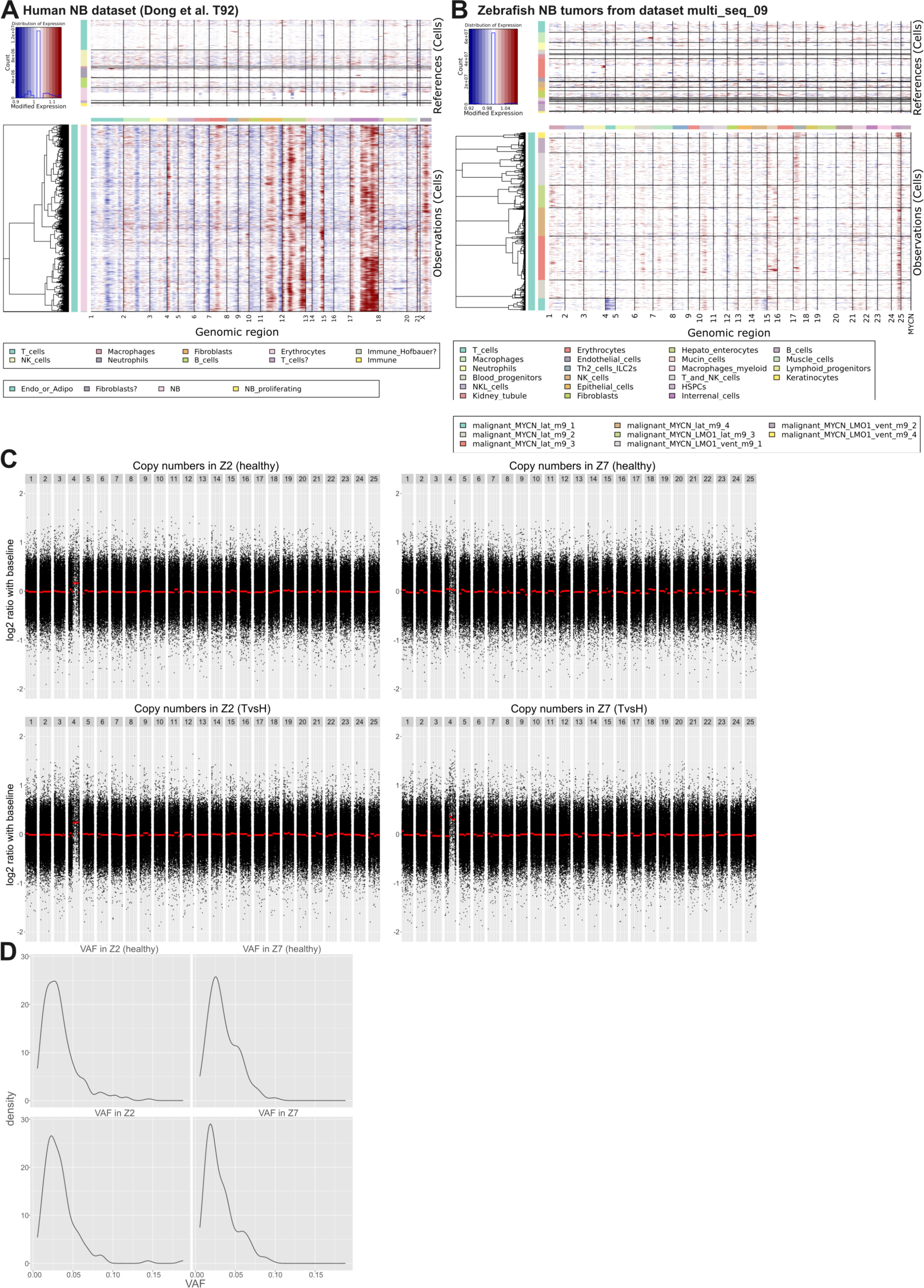
Genetic analysis of zebrafish NB based on inferred CNVs and whole exome sequencing. A) inferCNV result for a human NB scRNA-seq dataset (T92 from Dong et al.^24^). Reference cells and background CNV probability profiles are shown at the top. The tumor cells in the bottom part of the plot show strong signals of CNVs, e.g. for an amplification of part of chromosome 17, which is frequently observed in NB. B) inferCNV result for all zebrafish NB tumors from one MULTI-seq run (multi_seq_09). Reference cells and background CNV probability profiles are shown at the top. Cells from different tumors are grouped and marked with a distinct color in the legend on the left. There is no strong CNV-signal. C) WES copy number profiles for two samples (Z2 on the left, Z7 on the right). CNV profiles are shown for healthy control tissue taken from the fin (top) and tumor tissue (bottom). D) WES variant allele frequencies for the Z2 and Z7 healthy control tissue and tumor tissue.

**Fig S10:**
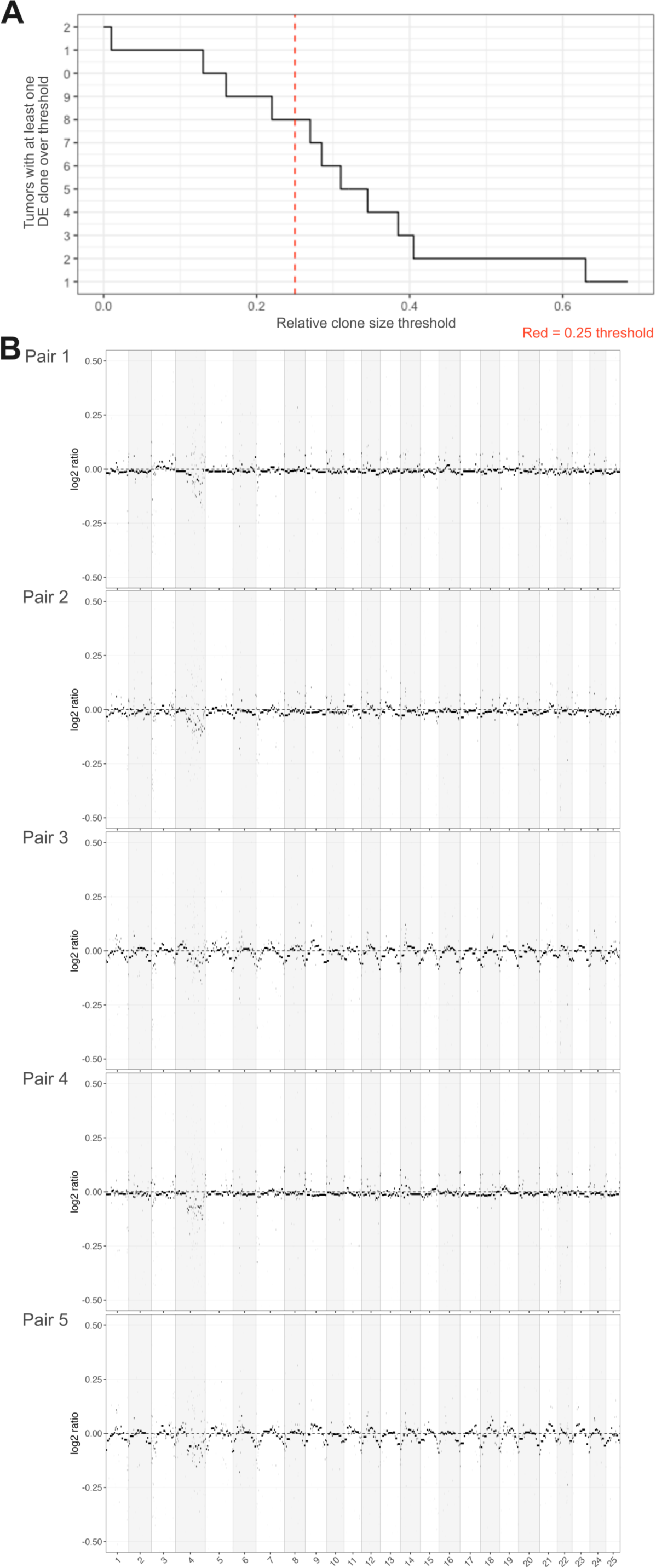
Clone sizes and WGS in MYCN-tumors. A) Size distribution of clones in MYCN-tumors with differential developmental cell-type specific module expression. Clone sizes are calculated as the number of cells in the clone divided by the total number of cells (cancer and TME) sequenced from the tumor. Eight clones (to the right of the threshold line) have a relative clone size of > 0.25. B) Copy-number profiles derived from WGS for each of five lateral MYCN-tumors, with respect to normal tissue samples from the same animal

**Fig S11:**
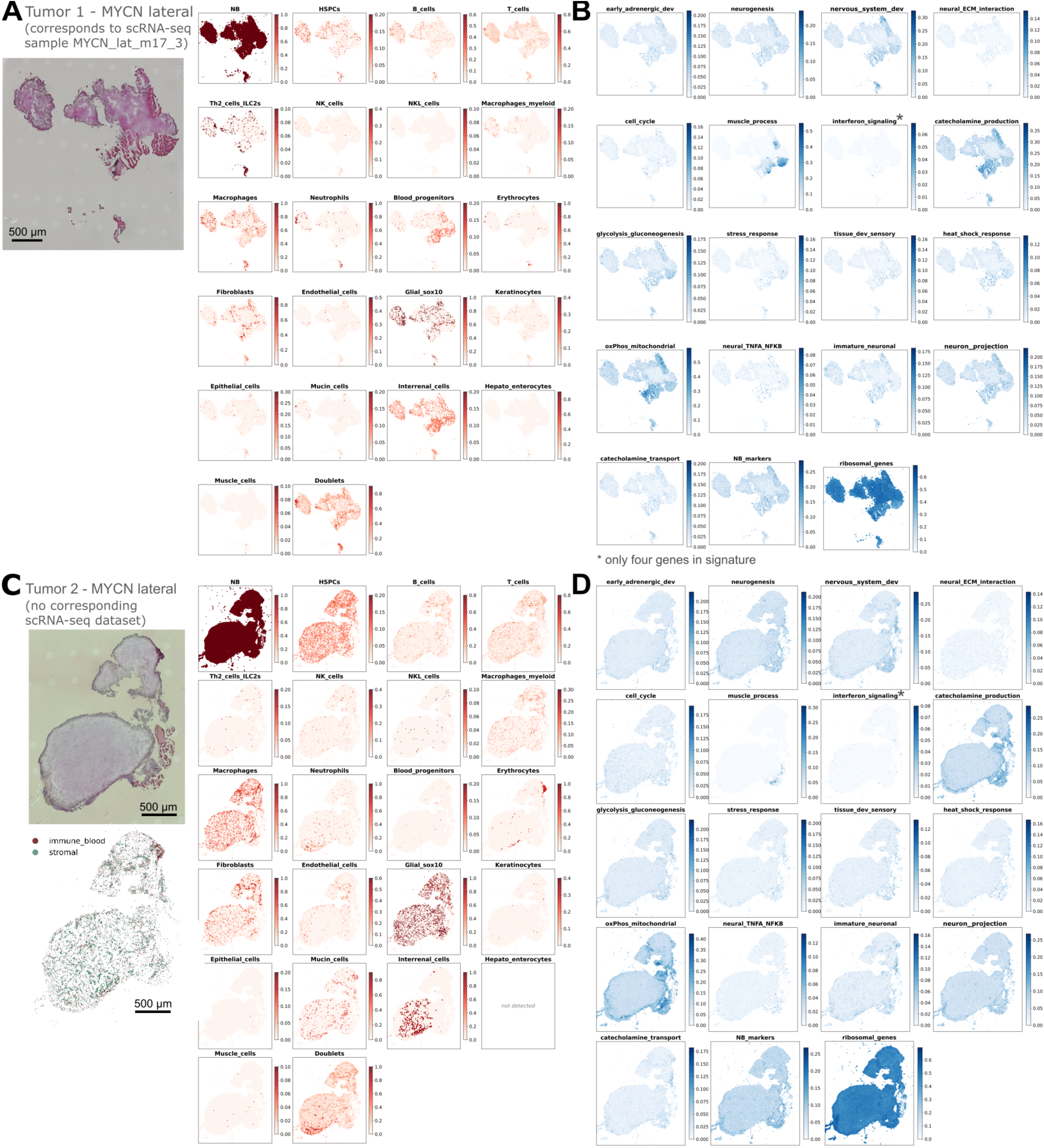
Spatial transcriptomics of two lateral MYCN-tumors. A) Light microscopic image of the H&E-stained section taken for Open-ST from tumor 1 as well as reference cell type similarity scores in 5 µm spots, using cell type marker genes derived from scRNA-seq data (as described in Methods section ‘Spatial analysis of cell type and tumor modules’). B) Expression score for NB gene expression modules in 5 µm spots across the tumor section. C) Light microscopic image of the H&E-stained section taken for Open-ST from tumor 2 as well as reference cell type similarity scores in 5 µm spots, using cell type marker genes derived from scRNA-seq data. D) Expression score for all NB gene expression modules in 5 µm spots across the tumor section.

**Fig S12:**
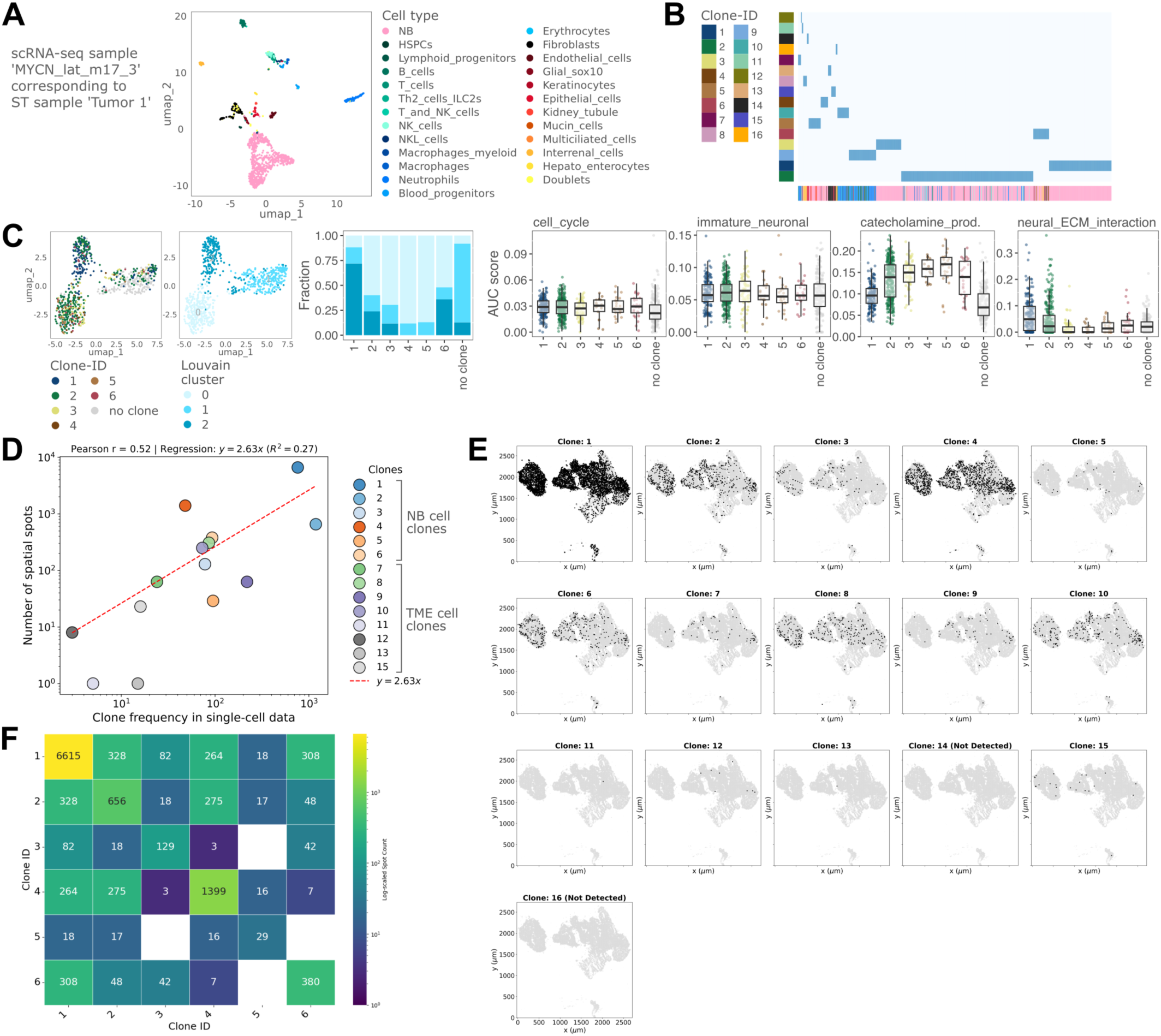
Spatial transcriptomics with lineage tracing. A) UMAP of single cell transcriptomes derived from scRNA-seq from the same tumor that was profiled with spatial transcriptomics with cells colored according to cell type. B) Hierarchical clustering of those cells from the tumor shown in A that were assigned to one of 16 clones (dark blue heatmap color: assigned to clone). Cells are annotated by cell type at the bottom of the heatmap. Clones 1 to 6 contain many NB cells while other clones are composed of TME cells. C) UMAP of sub-clustered NB cells from the tumor shown in A. Cells are colored according to the clone or Louvain cluster that they were assigned to. The barplot shows the fraction of cells from each clone assigned to a given Louvain cluster. Boxplots and jitter plots show the expression score for the indicated modules per clone. D) Comparison between number of cells in the scRNA-seq dataset that were assigned to a clone (x-axis) and number of spatial spots, in which a clone-specific sequence was found (y-axis) shows an overall agreement in the relative clone sizes across both data modalities. E) Spatial outline of the tumor section highlighting spots, in which lineage barcode sequences representative of the indicated clones (derived from scRNA-seq data) were found. Clones 1 to 6 were defined as large NB cell clones in the scRNA-seq data and are also shown in Fig. 4G. The other clones shown here either had very few NB cells or were completely composed of TME cells in the scRNA-seq data. F) Heatmap showing spatial overlap of the 6 NB cell clones. The numbers indicate the number of spots, in which at least one read for each of two clonal lineage barcodes was found. The diagonal shows the total number of spots in which at least one lineage barcode of a particular clone was found. Most clones overlap in space to a certain extent with differences between clone pairs, e.g. almost 80 % of all spots with clone 6 labels also have a clone 1 label; in contrast, only about 20 % of all spots with clone 4 labels also have a clone 1 label and clone 4 and clone 6 have little overlap in space.

**Fig S13:**
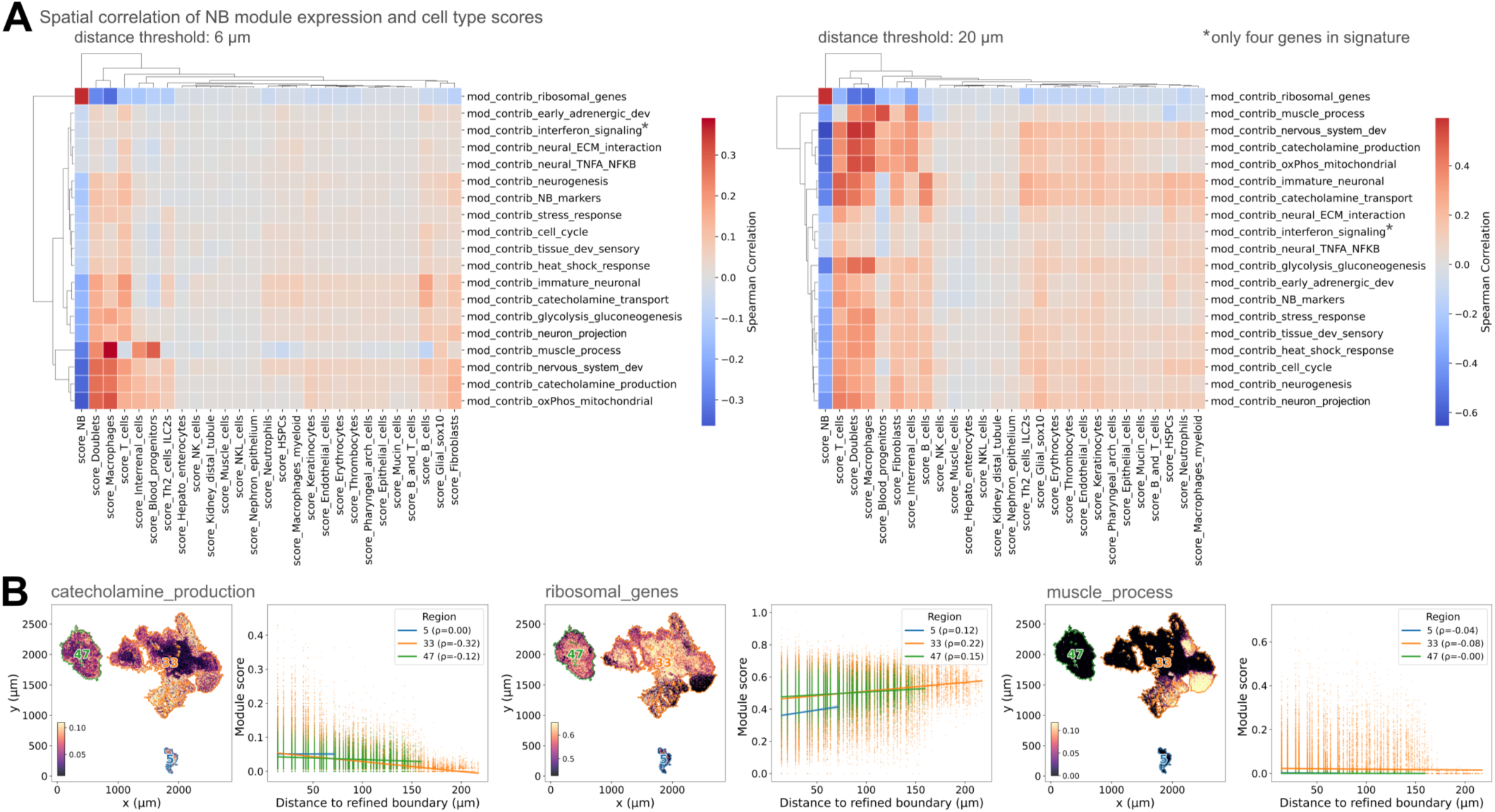
Association of NB gene module expression with tumor spatial features. A) Co-localization of NB cells expressing distinct modules and TME cell types in spatial transcriptomics: Correlation of activation of NB gene expression programs and scores for TME cell types in spots that are directly neighboring each other (left) or that are found in a slightly larger neighborhood (right). Spots directly neighboring each other (6 µm) have an average of 6.53 neighbors, while the larger neighborhood (20 µm) contains an average of 56.69 neighbors for one spot. B) Spatial distribution of module expression relative to the tumor border for three modules (*ribosomal_genes*, *catecholamine_production*, *muscle_process*). The tumor border was defined from the microscopic image and is indicated in orange, green or blue depending on tumor region. Next to expression scores on the tissue section, spots are plotted according to their distance from the tumor border and their expression of a given module. *Ribosomal_genes* expression is positively correlated with distance from the border, *catecholamine_production* is inversely correlated with it and *muscle_process* doesn’t show a systematic association.

**Fig S14:**
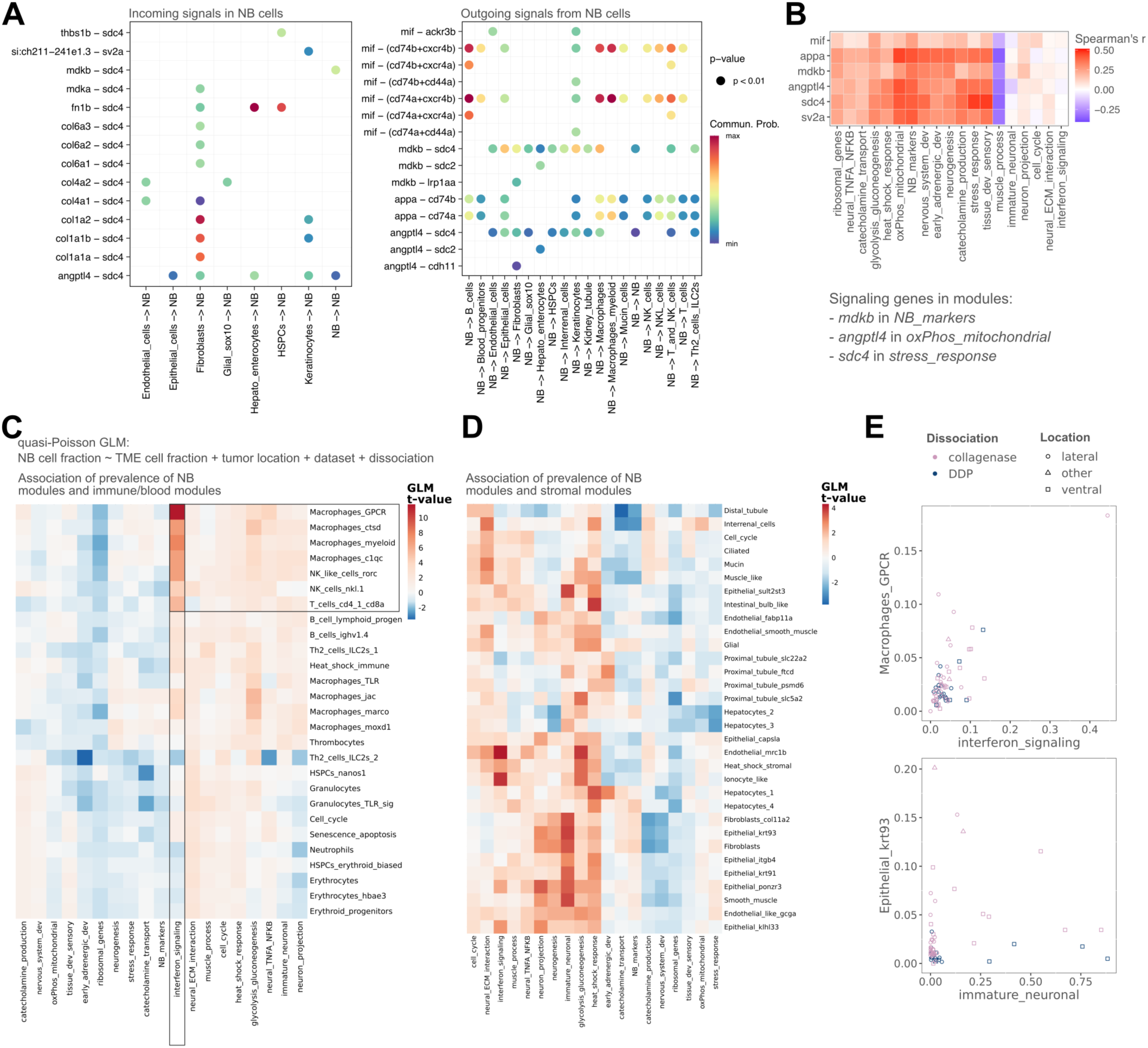
Association of NB gene module expression with presence of TME cell types in scRNA-seq data. A) CellChat-inferred NB-specific interactions. Left plot shows signals received by NB cells and the right plot shows signaling interactions with NB cells as senders. Interaction strength is a relative measure determined by Cellchat. B) Spearman correlation between expression of ligand or receptor molecules (x-axis) and gene modules (AUCell scores, y axis) across NB cells of all primary tumors. Ligand or receptor molecules that are part of one of the gene modules are indicated below the plot. C) Co-occurrence of NB cells in a specific state and individual TME cell types in scRNA-seq data: Heatmap showing t-values of association of fraction of TME cells with expression of modules representative of immune / blood cells (rows) and fraction of NB cells expressing NB modules (columns) in a generalized linear model considering the dataset, dissociation method and tumor location as covariates. Only modules with at least one significant association after correction for multiple testing are shown. Association of interferon signaling activation in NB cells and immune cells expressing macrophage-like programs is highlighted as an example of putative functional association (see scatterplot in E). D) Heatmap showing t-values of association of fraction of TME cells expressing modules representative of stromal cells (rows) and fraction of NB cells expressing NB modules (columns) in a generalized linear model (as in C). Only modules with at least one significant association are shown. E) Scatter plot showing fraction of NB cells expressing the module *interferon_signaling* and fraction of TME cells expressing the module ‘Macrophages_GPCR’ per tumor (top) shows a visible association of the two values. Tumors are colored by dissociation protocol used and shaped according to the tumor location. Association of TME cells expressing the module ‘Epithelial_krt93’ and NB cells with *immature_neuronal* activation looks less clear (bottom).

**Fig S15:**
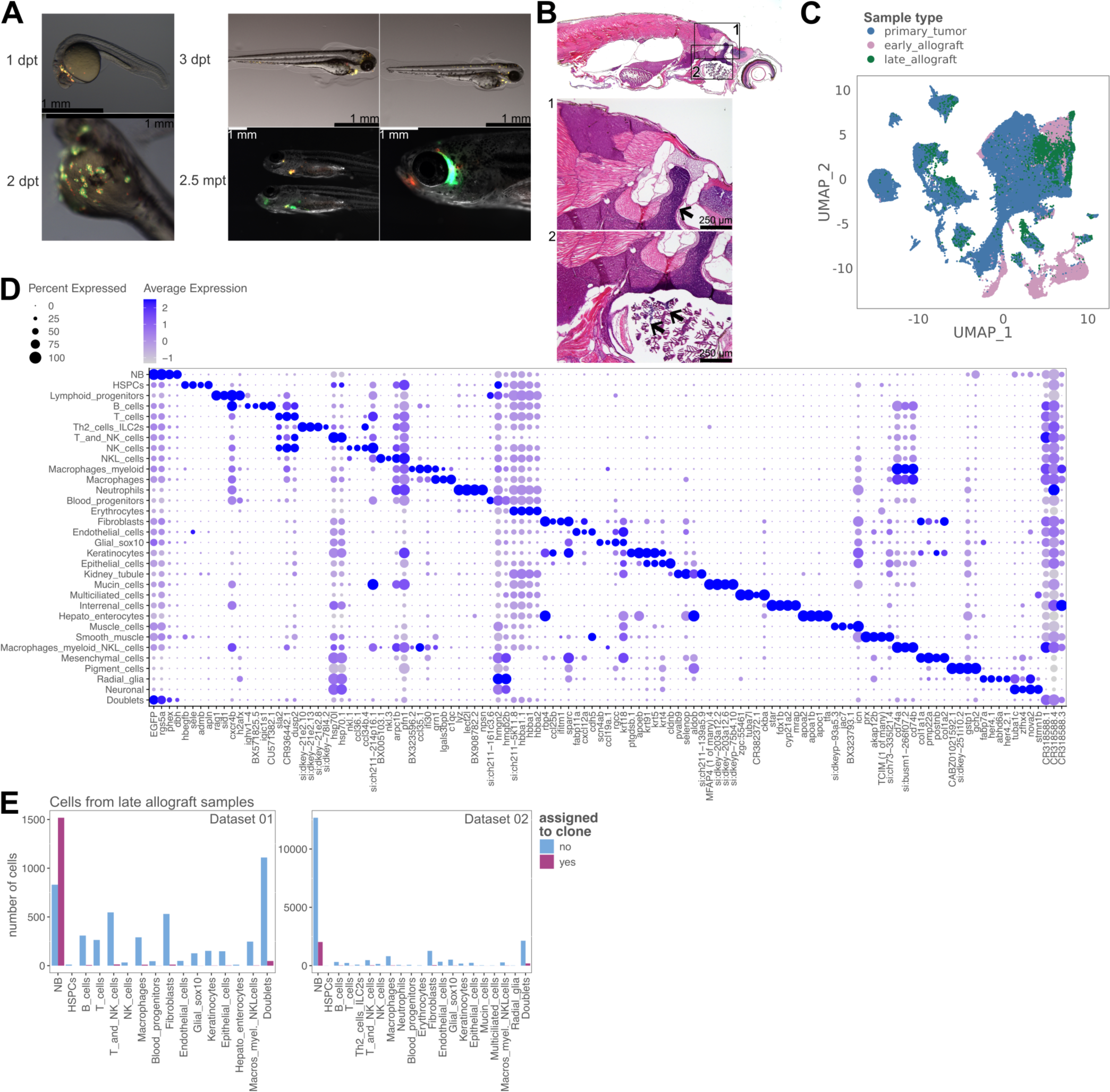
Tumor cell allogeneic transplantation and scRNA-seq of allograft tumors. A) Example images of fluorescent tumor cells (mCherry- and GFP-channels overlaid on the light microscopic image) in host fish from 1 day post transplantation (dpt) to 2.5 months post transplantation (mpt). B) H&E-stained sagittal section of a 3 months old host fish with graft tumors. The insets highlight (1) a large lateral tumor mass close to the superior cervical ganglion and (2) tumor cell foci along the gills. C) UMAPs of all cells from all timepoints colored by sampling time point, showing that a large population of putative NB cells from the early and late allograft stage cluster together with primary tumor NB cells. D) Dotplot showing top four marker genes determined by differential expression for each cell type in the dataset with cells from all timepoints (shown in C). E) Detection of NB-clone-specific lineage barcodes (derived from primary tumors) in cells of late allograft tumors in two separate experiments, in which data from all three timepoints was collected (primary tumor, early allograft, late allograft). Mainly graft cells identified as NB cells based on gene expression were assigned to primary tumor clones, while relatively few other cell types carried lineage barcodes also found in the primary tumor.

**Fig S16:**
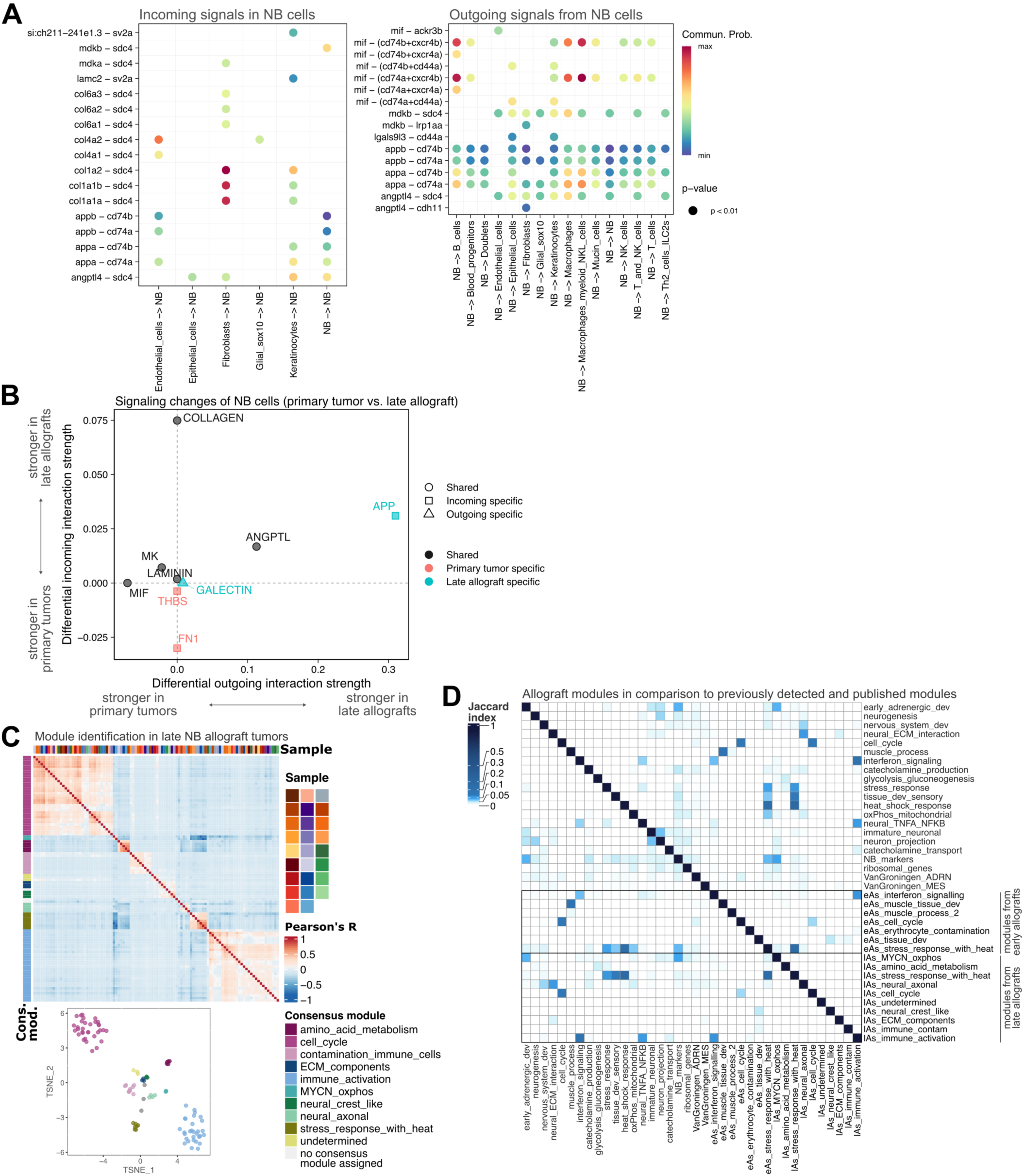
Cell-cell communication and expression program activation in allografted NB cells. A) CellChat-inferred NB-specific interactions in all late allograft tumors. Left plot shows signaling interactions received by NB cells and the right plot shows signaling interactions with NB cells as senders. Interaction strength is a relative measure determined by CellChat. B) Differential interaction strength measured for NB cells in primary tumors compared to late allograft tumors with CellChat. C) Heatmap and T-SNE showing modules detected in the analysis of individual late allograft tumors plotted according to their pairwise correlation to each other (Pearson). Modules in the heatmap are annotated by the individual late graft tumor sample they were derived from (color legend in top row) and by the consensus module they were assigned to after clustering with HDBScan (color legend on the left side). Modules in the T-SNE are colored according to the consensus module they were assigned to after clustering with HDBScan. Modules that were not assigned to any consensus module are labelled NA and colored in grey. D) Summary of final list of modules derived from the primary tumor samples, the published human ADRN/adrenergic and MES/mesenchymal gene signatures, and the modules derived from early graft NB cells (modules with prefix ‘eAs’) or late graft tumors (modules with prefix ‘lAs’). Jaccard index shows gene content overlap for all modules.

**Fig. S17:**
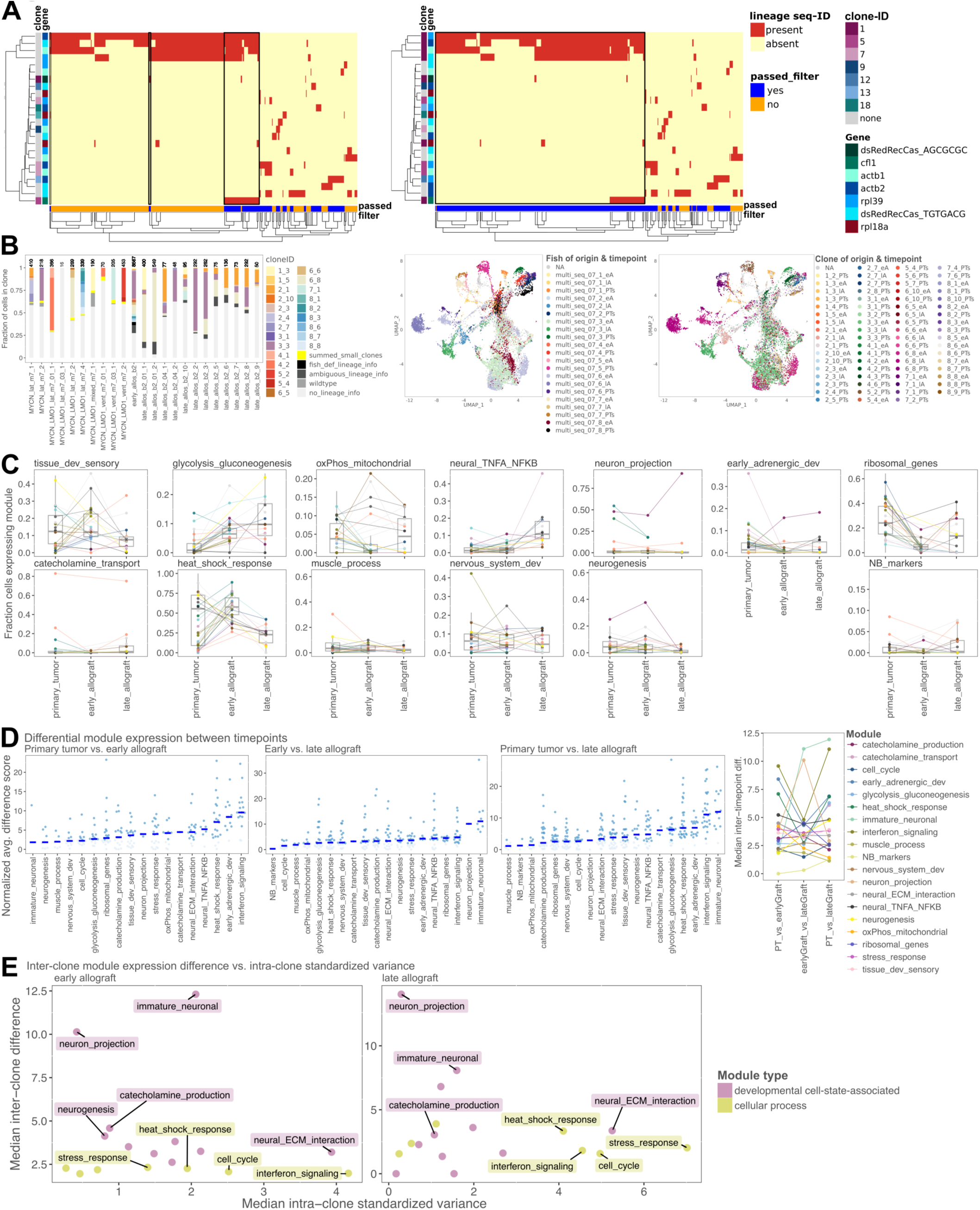
Clonal dynamics and gene module expression changes across allografting timepoints. A) Heatmaps of all cells from one tumor (columns) that passed filtering for at least one target gene versus alleles (for transgene) or allele combinations (for endogenous targets) from all target genes (rows). Alleles are colored by the clone they are definitive of left of the heatmap. Alleles that did not pass filtering are shown in grey. Cells that could unambiguously be assigned to a single clone are highlighted in orange (left heatmap) or blue (right heatmap). The left heatmap represents clone calling, which maximizes resolution in the assignment of cells to clones. This leads to the exclusion of cells, which only carry lineage barcodes that were created early, in favor of splitting subgroups of this population into smaller clones based on lineage barcodes created later (black boxes). The heatmap on the right shows the result of lower resolution clone calling as used for the analyses in Fig. 5 and 6. Here, retaining the large group of cells excluded in the high-resolution analysis is favored (black box). Many cells are placed into one clone regardless of the additional lineage barcodes they may have obtained later. B) Clonal dynamics for one experiment with cells from all three timepoints (primary tumor, early allograft, late allograft). Barplot shows composition of primary tumors, early graft samples and late graft tumors by clone. Very small clones were grouped (‘summed_small_clones’). Cells that could not be assigned to a clone, but to a primary tumor fish of origin are shown in black (‘fish_def_lineage_info’). Cells that only carry lineage barcodes that are ambiguous in terms of their origin are shown in dark grey (‘ambiguous_lineage_info’). Cells with only uncut lineage reads or lacking lineage info entirely are shown in lighter grey hues. Left UMAP shows cells colored according to fish of origin (for primary tumors) or assigned fish of origin (for graft samples) and sampling time point (PTs = primary tumor, eA = early allograft, lA = late allograft). Right UMAP shows cells colored according to clone/fish of origin (for primary tumors) or assigned clone/fish of origin (for graft samples) and sampling time point. G) Fraction of cells per clone and time point that express a given module. Each dot shows a clone in a specific time point and groups of cells from the same clone in different time points are connected via lines (as in Fig. 6A). C) Differential module expression scores for comparisons between cells from one clone found in two different time points (as indicated above the plots) with the blue bars indicating the median. Summary plot on the right shows median values of differential expression for each timepoint and module. D) Scatterplots as in Fig. 4F for early allograft (left) and late allograft (right) timepoints. Plots show median inter-clone difference in module expression (between clones within one early allograft sample or late allograft tumor) and median intra-clone variance for cells from individual clones found in one early or late graft sample.

**Fig. S18:**
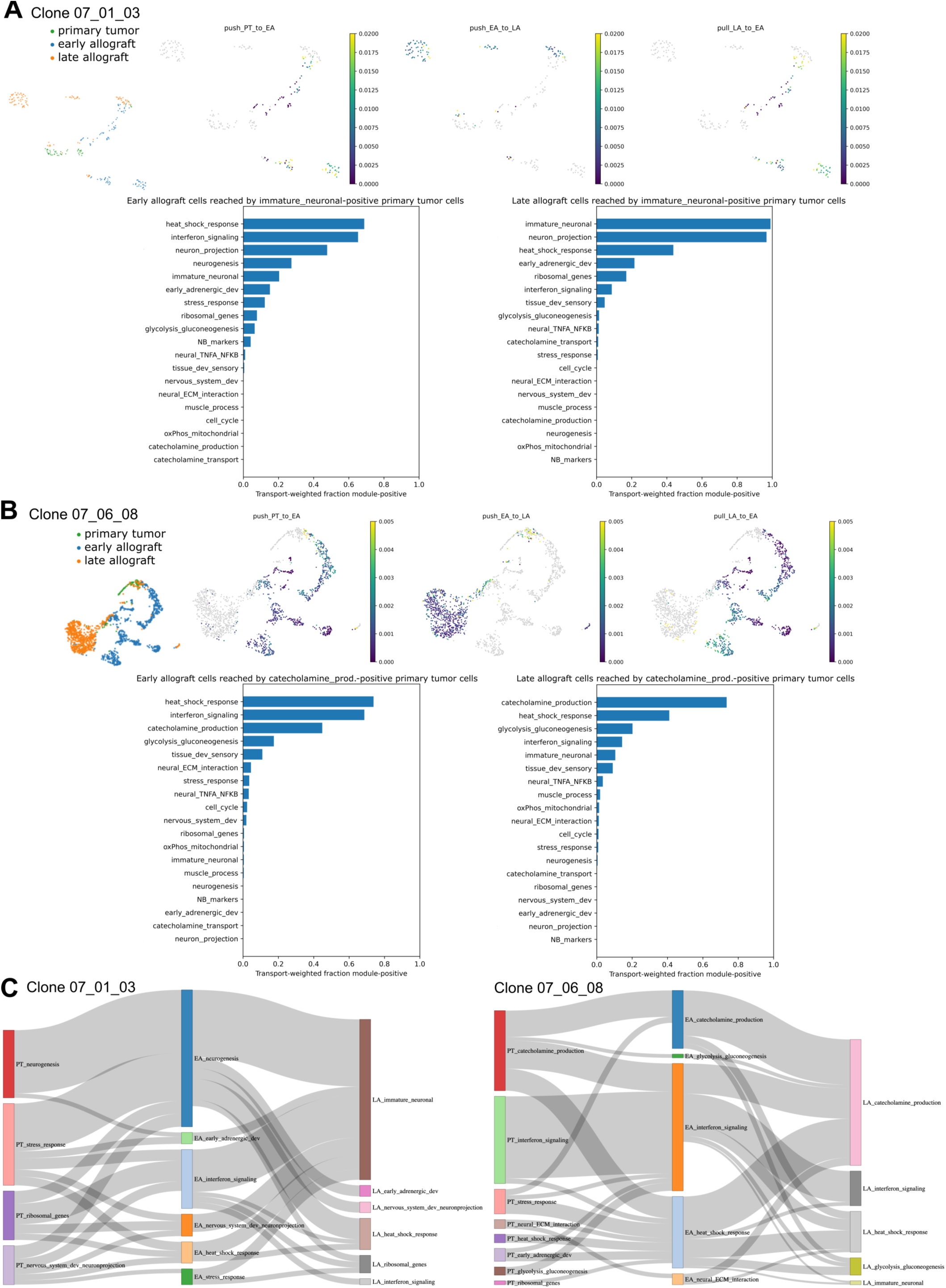
Cell state transition inference shows that developmental cell state-associated modules can be activated or deactivated upon transplantation. A) UMAP of NB cells from a single clone detected in primary tumor multi_seq_07_01 (MYCN;LMO1), early allograft and late allograft samples. UMAPs are colored by push forward from primary tumor to early allograft, early allograft to late allograft or pull back from late allograft to early allograft (from left to right). Barplots show the fraction of cells in the early allograft timepoint (left) or late allograft timepoint (right) with a high probability of being derived from primary tumor cells with *immature_neuronal* expression that express a given module. B) UMAP of cells from a single clone detected in primary tumor multi_seq_07_06 (MYCN;LMO1), early allograft and late allograft samples. UMAPs are colored by push forward from primary tumor to early allograft, early allograft to late allograft or pull back from late allograft to early allograft (from left to right). Barplots show the fraction of cells in the early allograft timepoint (left) or late allograft timepoint (right) derived from primary tumor cells with *catecholamine_production* expression that express a given module. C) Sankey plots for the two clones shown in A and B showing inferred transitions between cells with expression of different modules. Cells were assigned to the module, which they express most highly.

## References

1. Marin-Bejar, O. et al. Evolutionary predictability of genetic versus nongenetic resistance to anticancer drugs in melanoma. Cancer Cell 39, 1135–1149.e8 (2021).

2. Moorman, A. et al. Progressive plasticity during colorectal cancer metastasis. Nature 637, 947–954 (2025).

3. Barkley, D. et al. Cancer cell states recur across tumor types and form specific interactions with the tumor microenvironment. Nat. Genet. 54, 1192–1201 (2022).

4. Gavish, A. et al. Hallmarks of transcriptional intratumour heterogeneity across a thousand tumours. Nature 618, 598–606 (2023).

5. Ciriello, G. et al. Cancer Evolution: A Multifaceted Affair. Cancer Discov. 14, 36– 48 (2024).

6. Moiso, E. et al. Developmental Deconvolution for Classification of Cancer Origin. Cancer Discov. 12, 2566–2585 (2022).

7. Yang, D. et al. Lineage tracing reveals the phylodynamics, plasticity, and paths of tumor evolution. Cell 185, 1905–1923.e25 (2022).

8. Quinn, J. J. et al. Single-cell lineages reveal the rates, routes, and drivers of metastasis in cancer xenografts. Science 371, eabc1944 (2021).

9. Simeonov, K. P. et al. Single-cell lineage tracing of metastatic cancer reveals selection of hybrid EMT states. Cancer Cell 39, 1150–1162.e9 (2021).

10. Filbin, M. & Monje, M. Developmental origins and emerging therapeutic opportunities for childhood cancer. Nat. Med. 25, 367–376 (2019).

11. Pugh, T. J. et al. The genetic landscape of high-risk neuroblastoma. Nat. Genet. 45, 279–284 (2013).

12. Schulte, J. H. & Eggert, A. Neuroblastoma. Crit. Rev. Oncog. 20, 245–270 (2015).

13. Körber, V. et al. Neuroblastoma arises in early fetal development and its evolutionary duration predicts outcome. Nat. Genet. 55, 619–630 (2023).

14. Maris, J. M., Hogarty, M. D., Bagatell, R. & Cohn, S. L. Neuroblastoma. The Lancet 369, 2106–2120 (2007).

15. Westermann, F. et al. Distinct transcriptional MYCN/c-MYC activities are associated with spontaneous regression or malignant progression in neuroblastomas. Genome Biol. 9, (2008).

16. Rickman, D. S., Schulte, J. H. & Eilers, M. The Expanding World of N-MYC– Driven Tumors. Cancer Discov. 8, 150–163 (2018).

17. Zhu, S. et al. Activated ALK Collaborates with MYCN in Neuroblastoma Pathogenesis. Cancer Cell 21, 362–373 (2012).

18. Olsen, R. R. et al. MYCN induces neuroblastoma in primary neural crest cells. Oncogene 36, 5075–5082 (2017).

19. Van Groningen, T. et al. Neuroblastoma is composed of two super-enhancer-associated differentiation states. Nat. Genet. 49, 1261–1266 (2017).

20. Van Groningen, T. et al. A NOTCH feed-forward loop drives reprogramming from adrenergic to mesenchymal state in neuroblastoma. Nat. Commun. 10, 1530 (2019).

21. Boeva, V. et al. Heterogeneity of neuroblastoma cell identity defined by transcriptional circuitries. Nat. Genet. 49, 1408–1413 (2017).

22. Wang, L. et al. ASCL1 is a MYCN- and LMO1-dependent member of the adrenergic neuroblastoma core regulatory circuitry. Nat. Commun. 10, (2019).

23. Jansky, S. et al. Single-cell transcriptomic analyses provide insights into the developmental origins of neuroblastoma. Nat. Genet. 53, 683–693 (2021).

24. Dong, R. et al. Single-Cell Characterization of Malignant Phenotypes and Developmental Trajectories of Adrenal Neuroblastoma. Cancer Cell 38, 716–733.e6 (2020).

25. Kildisiute, G. et al. Tumor to normal single-cell mRNA comparisons reveal a pan-neuroblastoma cancer cell. Sci. Adv. (2021).

26. Olsen, T. K. et al. Joint single-cell genetic and transcriptomic analysis reveal pre-malignant SCP-like subclones in human neuroblastoma. Mol. Cancer 23, 180 (2024).

27. Yuan, X. et al. Single-cell profiling of peripheral neuroblastic tumors identifies an aggressive transitional state that bridges an adrenergic-mesenchymal trajectory. Cell Rep. 41, 111455 (2022).

28. Kameneva, P. et al. Single-cell transcriptomics of human embryos identifies multiple sympathoblast lineages with potential implications for neuroblastoma origin. Nat. Genet. 53, 694–706 (2021).

29. Van Haver, S. et al. Human iPSC modeling recapitulates in vivo sympathoadrenal development and reveals an aberrant developmental subpopulation in familial neuroblastoma. iScience 27, 108096 (2024).

30. Saldana-Guerrero, I. M. et al. A human neural crest model reveals the developmental impact of neuroblastoma-associated chromosomal aberrations. Nat. Commun. 15, 3745 (2024).

31. Tao, T. et al. The pre-rRNA processing factor DEF is rate limiting for the pathogenesis of MYCN-driven neuroblastoma. Oncogene 36, 3852–3867 (2017).

32. Zhu, S. et al. LMO1 Synergizes with MYCN to Promote Neuroblastoma Initiation and Metastasis. Cancer Cell 32, 310–323.e5 (2017).

33. Spanjaard, B. et al. Simultaneous lineage tracing and cell-type identification using CRISPR–Cas9-induced genetic scars. Nat. Biotechnol. 36, 469–473 (2018).

34. Mitic, N. et al. Dissecting the spatiotemporal diversity of adult neural stem cells. Mol. Syst. Biol. 20, 321–337 (2024).

35. Chan, M. M. et al. Molecular recording of mammalian embryogenesis. Nature 570, 77–82 (2019).

36. McGinnis, C. S. et al. MULTI-seq: sample multiplexing for single-cell RNA sequencing using lipid-tagged indices. Nat. Methods 16, 619–626 (2019).

37. Pei, D. et al. Distinct Neuroblastoma-associated Alterations of PHOX2B Impair Sympathetic Neuronal Differentiation in Zebrafish Models. PLoS Genet. 9, e1003533 (2013).

38. Guo, S. et al. Mutations in the Zebrafish Unmask Shared Regulatory Pathways Controlling the Development of Catecholaminergic Neurons. Dev. Biol. 208, 473–487 (1999).

39. Finegold, M. J., Triche, T. J. & Askin, F. B. Neuroblastoma and the differential diagnosis of small-, round-, blue-cell tumors. Hum. Pathol. 14, 569–595 (1983).

40. Bei, Y. et al. Passenger Gene Coamplifications Create Collateral Therapeutic Vulnerabilities in Cancer. Cancer Discov. 14, 492–507 (2024).

41. Tang, Q. et al. Dissecting hematopoietic and renal cell heterogeneity in adult zebrafish at single-cell resolution using RNA sequencing. J. Exp. Med. 214, 2875–2887 (2017).

42. Stöber, M. C. et al. Intercellular extrachromosomal DNA copy-number heterogeneity drives neuroblastoma cell state diversity. Cell Rep. 43, 114711 (2024).

43. Hald, Ø. H. et al. Inhibitors of ribosome biogenesis repress the growth of MYCN-amplified neuroblastoma. Oncogene 38, 2800–2813 (2019).

44. Chapple, R. H. et al. An integrated single-cell RNA-seq map of human neuroblastoma tumors and preclinical models uncovers divergent mesenchymal-like gene expression programs. Genome Biol. 25, 161 (2024).

45. Sur, A. et al. Single-cell analysis of shared signatures and transcriptional diversity during zebrafish development. Dev. Cell 58, 3028–3047.e12 (2023).

46. Lange, M. et al. A multimodal zebrafish developmental atlas reveals the state-transition dynamics of late-vertebrate pluripotent axial progenitors. Cell 187, 6742–6759.e17 (2024).

47. Kamenev, D. et al. Schwann cell precursors generate sympathoadrenal system during zebrafish development. J. Neurosci. Res. 99, 2540–2557 (2021).

48. Setty, M. et al. Characterization of cell fate probabilities in single-cell data with Palantir. Nat. Biotechnol. 37, 451–460 (2019).

49. Weiler, P., Lange, M., Klein, M., Pe’er, D. & Theis, F. CellRank 2: unified fate mapping in multiview single-cell data. Nat. Methods 21, 1196–1205 (2024).

50. Kotliar, D. et al. Identifying gene expression programs of cell-type identity and cellular activity with single-cell RNA-Seq. eLife 8, e43803 (2019).

51. Duijkers, F. A. M. et al. High Anaplastic Lymphoma Kinase Immunohistochemical Staining in Neuroblastoma and Ganglioneuroblastoma Is an Independent Predictor of Poor Outcome. Am. J. Pathol. 180, 1223–1231 (2012).

52. Schulte, J. H. et al. High *ALK* Receptor Tyrosine Kinase Expression Supersedes *ALK* Mutation as a Determining Factor of an Unfavorable Phenotype in Primary Neuroblastoma. Clin. Cancer Res. 17, 5082–5092 (2011).

53. Mossé, Y. P. et al. Identification of ALK as a major familial neuroblastoma predisposition gene. Nature 455, 930–935 (2008).

54. Ryl, T. et al. Cell-Cycle Position of Single MYC-Driven Cancer Cells Dictates Their Susceptibility to a Chemotherapeutic Drug. Cell Syst. 5, 237–250.e8 (2017).

55. Sengupta, S. et al. Mesenchymal and adrenergic cell lineage states in neuroblastoma possess distinct immunogenic phenotypes. Nat. Cancer 3, 1228–1246 (2022).

56. Costa, A. et al. Single-cell transcriptomics reveals shared immunosuppressive landscapes of mouse and human neuroblastoma. J. Immunother. Cancer 10, e004807 (2022).

57. Patel, A. G. et al. A spatial cell atlas of neuroblastoma reveals developmental, epigenetic and spatial axis of tumor heterogeneity. Preprint at 10.1101/2024.01.07.574538 (2024).

58. Verhoeven, B. M. et al. The immune cell atlas of human neuroblastoma. Cell Rep. Med. 3, 100657 (2022).

59. Brady, S. W. et al. Pan-neuroblastoma analysis reveals age- and signature-associated driver alterations. Nat. Commun. 11, 5183 (2020).

60. SEQC/MAQC-III Consortium. A comprehensive assessment of RNA-seq accuracy, reproducibility and information content by the Sequencing Quality Control Consortium. Nat. Biotechnol. 32, 903–914 (2014).

61. Heath, A. P. et al. The NCI Genomic Data Commons. Nat. Genet. 53, 257–262 (2021).

62. Schott, M. et al. Open-ST: High-resolution spatial transcriptomics in 3D. Cell 187, 3953–3972.e26 (2024).

63. Zhou, Q., Yan, X., Gershan, J., Orentas, R. J. & Johnson, B. D. Expression of Macrophage Migration Inhibitory Factor by Neuroblastoma Leads to the Inhibition of Antitumor T Cell Reactivity In Vivo. J. Immunol. 181, 1877–1886 (2008).

64. DuBois, S. G. et al. Metastatic Sites in Stage IV and IVS Neuroblastoma Correlate With Age, Tumor Biology, and Survival. J. Pediatr. Hematol. Oncol. 21, (1999).

65. Rocha, M., Singh, N., Ahsan, K., Beiriger, A. & Prince, V. E. Neural crest development: insights from the zebrafish. Dev. Dyn. 249, 88–111 (2020).

66. Klein, D. et al. Mapping cells through time and space with moscot. Nature 638, 1065–1075 (2025).

67. Villalard, B. et al. Neuroblastoma plasticity during metastatic progression stems from the dynamics of an early sympathetic transcriptomic trajectory. Nat. Commun. 15, 9570 (2024).

68. Grossmann, L. D. et al. Identification and Characterization of Chemotherapy-Resistant High-Risk Neuroblastoma Persister Cells. Cancer Discov. 14, 2387–2406 (2024).

69. Westerfield, M. A Guide for the Laboratory Use of Zebrafish (Danio Rerio). (University of Oregon Press, Eugene, 2007).

70. Pan, Y. A. et al. Zebrabow: multispectral cell labeling for cell tracing and lineage analysis in zebrafish. Development 140, 2835–2846 (2013).

71. Wallace, C. K. et al. Effectiveness of Rapid Cooling as a Method of Euthanasia for Young Zebrafish (*Danio rerio*). J Am Assoc Lab Anim Sci 57, 58–63 (2018).

72. Replogle, J. M. et al. Combinatorial single-cell CRISPR screens by direct guide RNA capture and targeted sequencing. Nat. Biotechnol. 38, 954–961 (2020).

73. McKenna, A. et al. The Genome Analysis Toolkit: A MapReduce framework for analyzing next-generation DNA sequencing data. Genome Res. 20, 1297–1303 (2010).

74. Talevich, E., Shain, A. H., Botton, T. & Bastian, B. C. CNVkit: Genome-Wide Copy Number Detection and Visualization from Targeted DNA Sequencing. PLOS Comput. Biol. 12, e1004873 (2016).

75. Olshen, A. B. et al. Parent-specific copy number in paired tumor–normal studies using circular binary segmentation. Bioinformatics 27, 2038–2046 (2011).

76. Venkatraman, E. S. & Olshen, A. B. A faster circular binary segmentation algorithm for the analysis of array CGH data. Bioinformatics 23, 657–663 (2007).

77. Rubin, S. A. et al. Single-cell analyses reveal early thymic progenitors and pre-B cells in zebrafish. J Exp Med 219, e20220038 (2022).

78. Hahsler, M., Piekenbrock, M. & Doran, D. **dbscan**: Fast Density-Based Clustering with *R*. J. Stat. Softw. 91, (2019).

79. Aibar, S. et al. SCENIC: single-cell regulatory network inference and clustering. Nat. Methods 14, 1083–1086 (2017).

80. Chen, L. et al. CRISPR-Cas9 screen reveals a MYCN-amplified neuroblastoma dependency on EZH2. J. Clin. Invest. 128, 446–462 (2017).

81. Bonine, N. et al. NBAtlas: A harmonized single-cell transcriptomic reference atlas of human neuroblastoma tumors. Cell Rep. 43, 114804 (2024).

82. Love, M. I., Huber, W. & Anders, S. Moderated estimation of fold change and dispersion for RNA-seq data with DESeq2. Genome Biol. 15, 550 (2014).

83. Decoene, I., Herpelinck, T., Geris, L., Luyten, F. P. & Papantoniou, I. Engineering bone-forming callus organoid implants in a xenogeneic-free differentiation medium. Front. Chem. Eng. 4, 892190 (2022).

84. Tirosh, I. et al. Dissecting the multicellular ecosystem of metastatic melanoma by single-cell RNA-seq. Science 352, 189–196 (2016).

85. Sztanka-Toth, T. R., Jens, M., Karaiskos, N. & Rajewsky, N. Spacemake: processing and analysis of large-scale spatial transcriptomics data.

86. Korsunsky, I. et al. Fast, sensitive and accurate integration of single-cell data with Harmony. Nat. Methods 16, 1289–1296 (2019).

